# Leveraging Endogenous ADAR for Programmable Editing on RNA

**DOI:** 10.1101/605972

**Authors:** Liang Qu, Zongyi Yi, Shiyou Zhu, Chunhui Wang, Zhongzheng Cao, Zhuo Zhou, Pengfei Yuan, Ying Yu, Feng Tian, Zhiheng Liu, Ying Bao, Yanxia Zhao, Wensheng Wei

**Affiliations:** Biomedical Pioneering Innovation Center, Beijing Advanced Innovation Center for Genomics, Peking-Tsinghua Center for Life Sciences, Peking University Genome Editing Research Center, State Key Laboratory of Protein and Plant Gene Research, School of Life Sciences, Peking University, Beijing 100871, China; Peking University-Tsinghua University-National Institute of Biological Sciences Joint Graduate Program (PTN), Peking University, 100871, China; Academy for Advanced Interdisciplinary Studies, Peking University, Beijing 100871, China; EdiGene Inc, Life Science Park, 22 KeXueYuan Road, Changping District, Beijing 102206, China

## Abstract

Nucleic acid editing carries enormous potential for biological research and the development of therapeutics. Current tools for DNA or RNA editing rely on introducing exogenous proteins into living organisms, which is subject to potential risks or technical barriers due to possible aberrant effector activity, delivery limits and immunogenicity. Here, we report a programmable approach that employs a short RNA to leverage endogenous ADAR (Adenosine Deaminase Acting on RNA) proteins for targeted RNA editing. We engineered an RNA that is partially complementary to the target transcript to recruit native ADAR1 or ADAR2 to change adenosine to inosine at a specific site. We designated this new method as LEAPER (<u>L</u>everaging <u>E</u>ndogenous <u>A</u>DAR for <u>P</u>rogrammable <u>E</u>diting on <u>R</u>NA) and the ADAR-recruiting RNA as arRNA. arRNA, either expressed from plasmid or viral vector, or synthesized as an oligonucleotide, could achieve desirable editing. LEAPER has a manageable off-target rate on the targeted transcripts and rare global off-targets. We demonstrated that LEAPER could restore p53 function by repairing a specific cancer-relevant point mutation. Moreover, LEAPER could apply to a broad spectrum of cell types including multiple human primary cells, and it restored the α-L-iduronidase catalytic activity in Hurler syndrome patient-derived primary fibroblasts without evoking innate immune responses. As a single molecule system akin to RNAi, LEAPER enables precise and efficient RNA editing, offering the transformative potential for basic research and therapeutics.

## Introduction

Genome editing technologies are revolutionizing biomedical research. Highly active nucleases, such as zinc finger nucleases (ZFNs)^1^, transcription activator-like effector nucleases (TALENs)^2–4^, and Cas proteins of CRISPR (Clustered Regularly Interspaced Short Palindromic Repeats) system^5–7^ have been successfully engineered to manipulate the genome in a myriad of organisms. Recently, deaminases have been harnessed to precisely change the genetic code without breaking double-stranded DNA. By coupling a cytidine or an adenosine deaminase with the CRISPR-Cas9 system, researchers created programmable base editors that enable the conversion of C•G to T•A or A•T to G•C in genomic DNA^8–10^, offering novel opportunities for correcting disease-causing mutations.

Aside from DNA, RNA is an attractive target for genetic correction because RNA modification could alter the protein function without generating any permanent changes to the genome. The ADAR adenosine deaminases are currently exploited to achieve precise base editing on RNAs. Three kinds of ADAR proteins have been identified in mammals, ADAR1 (isoforms p110 and p150), ADAR2 and ADAR3 (catalytic inactive)^11,12^, whose substrates are double-stranded RNAs, in which an adenosine (A) mismatched with a cytosine (C) is preferentially deaminated to inosine (I). Inosine is believed to mimic guanosine (G) during translation^13,14^. To achieve targeted RNA editing, the ADAR protein or its catalytic domain was fused with a λN peptide^15–17^, a SNAP-tag^18–22^ or a Cas protein (dCas13b)^23^, and a guide RNA was designed to recruit the chimeric ADAR protein to the specific site. Alternatively, overexpressing ADAR1 or ADAR2 proteins together with an R/G motif-bearing guide RNA was also reported to enable targeted RNA editing^24–27^.

All these reported nucleic acid editing methods in mammalian system rely on ectopic expression of two components: an enzyme and a guide RNA. Although these binary systems work efficiently in most studies, some inherited obstacles limit their broad applications, especially in therapies. Because the most effective *in vivo* delivery for gene therapy is through viral vectors^28^, and the highly desirable adeno-associated virus (AAV) vectors are limited with cargo size (∼4.5 kb), making it challenging for accommodating both the protein and the guide RNA^29,30^. Over-expression of ADAR1 has recently been reported to confer oncogenicity in multiple myelomas due to aberrant hyper-editing on RNAs^31^, and to generate substantial global off-targeting edits^32^. In addition, ectopic expression of proteins or their domains of non-human origin has potential risk of eliciting immunogenicity^30,33^. Moreover, pre-existing adaptive immunity and p53-mediated DNA damage response may compromise the efficacy of the therapeutic protein, such as Cas9^34–38^. Although it has been attempted to utilize endogenous mechanism for RNA editing, this was tried only by injecting pre-assembled target transcript:RNA duplex into *Xenopus* embryos^39^. Alternative technologies for robust nucleic acid editing that don’t rely on ectopic expression of proteins are much needed. Here, we developed a novel approach that leverages endogenous ADAR for RNA editing. We showed that expressing a deliberately designed guide RNA enables efficient and precise editing on endogenous RNAs, and corrects pathogenic mutations. This unary nucleic acid editing platform may open new avenues for therapeutics and research.

## Results

### Leveraging endogenous ADAR for RNA editing

In an attempt to explore an efficient RNA editing platform, we fused the deaminase domain of the hyperactive E1008Q mutant ADAR1 (ADAR1_DD_)^40^ to the catalytic inactive LbuCas13 (dCas13a), an RNA-guided RNA-targeting CRISPR effector^41^ (Extended Data Fig. 1a). To assess RNA editing efficiency, we constructed a surrogate reporter harbouring *mCherry* and *EGFP* genes linked by a sequence comprising a 3× GGGGS-coding region and an in-frame UAG stop codon (Reporter-1, Extended Data Fig. 1b). The reporter-transfected cells only expressed mCherry protein, while targeted editing on the UAG of the reporter transcript could convert the stop codon to UIG and consequently permit the downstream EGFP expression. Such a reporter allows us to measure the A-to-I editing efficiency through monitoring EGFP level. We then designed hU6 promoter-driven crRNAs (CRISPR RNAs) containing 5’ scaffolds subjected for Cas13a recognition and variable lengths of spacer sequences for targeting (crRNA^Cas13a^, Supplementary Table 1). The sequences complementary to the target transcripts all contain CCA opposite to the UAG codon so as to introduce a cytidine (C) mis-pairing with the adenosine (A) (Extended Data Fig. 1b) because adenosine deamination preferentially occurs in the A-C mismatch site^13,14^. To test the optimal length of the crRNA, non-targeting or targeting crRNAs of different lengths were co-expressed with dCas13a-ADAR1_DD_ proteins in HEK293T cells stably expressing the Reporter-1. Evident RNA editing effects indicated by the appearance of EGFP expression were observed with crRNAs containing matching sequences at least 40-nt long, and the longer the crRNAs the higher the EGFP positive percentage (Extended Data Fig. 1c). Surprisingly, expression of long crRNA^Cas13a^ alone appeared sufficient to activate EGFP expression, and the co-expression of dCas13a-ADAR1_DD_ rather decreased crRNA activity (Extended Data Fig. 1c, d). The EGFP expression was clearly sequence-dependent because the 70-nt (exclusive of the 5’ scaffold for the length calculation) control RNA could not activate EGFP expression (Extended Data Fig. 1c, d).

With the surprising finding that certain long engineered crRNA^Cas13a^ enabled RNA editing independent of dCas13a-ADAR1_DD_, we decided to remove the Cas13a-recruiting scaffold sequence from the crRNA. Because the crRNA_70_ had the highest activity to trigger EGFP expression (Extended Data Fig. 1c, d), we chose the same 70-nt long guide RNA without the Cas13a-recruiting scaffold for further test (Fig. 1a and Supplementary Table 2). It turned out that this linear guide RNA induced strong EGFP expression in close to 40% of total cells harboring the Reporter-1 (Fig. 1b, upper). Because endogenous ADAR proteins could edit double-stranded RNA (dsRNA) substrates^12^, we reasoned that the long guide RNAs could anneal with the target transcripts to form dsRNA substrates that in turn recruit endogenous ADAR proteins for targeted editing. We thus designated such guide RNA as arRNA (ADAR-recruiting RNA).

**Figure 1.**
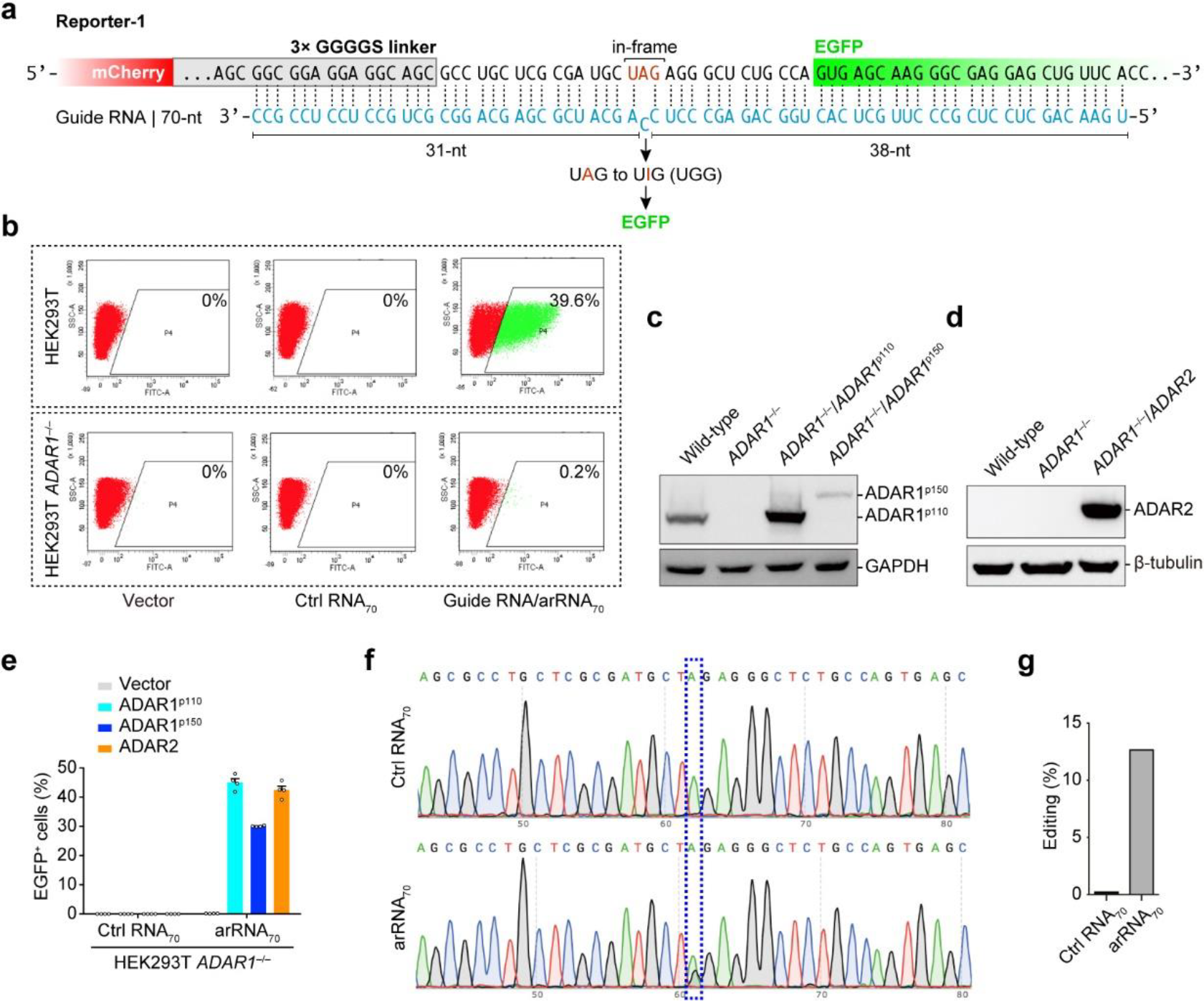
Leveraging endogenous ADAR1 protein for targeted RNA editing. **a**, Schematic of the Reporter-1 and the 70-nt arRNA. **b**, Representative FACS analysis of arRNA-induced EGFP expression in wild-type (HEK293T, upper) or ADAR1 knockout (HEK293T *ADAR1*^−/−^, lower) cells stably expressing the Repoter-1. **c**, Western blot analysis showing expression levels of ADAR1 proteins in wild-type and HEK293T *ADAR1*^−/−^ cells, as well as those in HEK293T *ADAR1*^−/−^ cells transfected with ADAR1 isoforms (p110 and p150). **d**, Western blot analysis showing expression levels of ADAR2 proteins in wild-type and HEK293T *ADAR1*^−/−^ cells, as well as those in HEK293T *ADAR1*^−/−^ cells transfected with ADAR2. **e**, Quantification of the EGFP positive (EGFP^+^) cells. Reporter-1 and indicated ADAR-expressing constructs were co-transfected into HEK293T *ADAR1*^−/−^ cells, along with the Ctrl RNA_70_ or with the targeting arRNA_70_, followed by FACS analysis. EGFP^+^ percentages were normalized by transfection efficiency, which was determined by mCherry^+^. Data are mean values ± s.e.m. (n = 4). **f**, The Electropherograms showing Sanger sequencing results in the Ctrl RNA_70_ (upper) or the arRNA_70_ (lower)-targeted region. **g**, Quantification of the A to I conversion rate at the targeted site by deep sequencing.

To verify if endogenous ADAR proteins are indeed responsible for above observation, we set out to examine the arRNA-mediated RNA editing in ADAR-deficient cells. Since *ADAR2* mRNA was barely detectable in HEK293T cells (Extended Data Fig. 2a), we generated HEK293T *ADAR1*^−/−^ cells, rendering this cell line deficient in both ADAR1 and ADAR2 (Fig. 1c, d). Indeed, the depletion of ADAR1 abrogated arRNA_70_-induced EGFP signals (Fig. 1b, lower). Moreover, exogenous expression of ADAR1^p110^, ADAR1^p150^ or ADAR2 in HEK293T *ADAR1*−/^−^ cells (Fig. 1c, d) successfully rescued the loss of EGFP induction by arRNA_70_ (Fig. 1e, Extended Data Fig. 2b), demonstrating that arRNA-induced EGFP reporter expression solely depended on native ADAR1, whose activity could be reconstituted by its either isoforms (p110 and p150) or ADAR2. Sanger sequencing analysis on the arRNA_70_-targeting region showed an A/G overlapping peak at the predicted adenosine site within UAG, indicating a significant A to I (G) conversion (Fig. 1f). The next-generation sequencing (NGS) further confirmed that the A to I conversion rate was about 13% of total reporter transcripts (Fig. 1g). The quantitative PCR analysis showed that arRNA_70_ did not reduce the expression of targeted transcripts (Extended Data Fig. 3), ruling out the possible RNAi effect of the arRNA. Collectively, our data demonstrated that the arRNA is capable of generating significant level of editing on the targeted transcripts through the engineered A-C mismatch. We thus designate this novel RNA editing method as **LEAPER** (**L**everaging **E**ndogenous **A**DAR for **P**rogrammable **E**diting on **R**NA).

### LEAPER enables RNA editing in multiple cell lines

Because the expression of endogenous ADAR proteins is a prerequisite for LEAPER-mediated RNA editing, we tested the performance of LEAPER in a panel of cell lines originated from distinct tissues, including HT29, A549, HepG2, RD, SF268, SW13 and HeLa. We first examined the endogenous expression of all three kinds of ADAR proteins using Western blotting analyses. ADAR1 was highly expressed in all tested cell lines, and its identity in the Western blots was confirmed by the negative control, HEK293T *ADAR1*^−/−^ line (Fig. 2a, b). ADAR3 was detected only in HepG2 and HeLa cells (Fig. 2a, b). ADAR2 was non-detectable in any cells, a result that was not due to the failure of Western blotting because ADAR2 protein could be detected from ADAR2-overexpressing HEK293T cells (Fig. 2a, b). These findings are in consistent with previous reports that ADAR1 is ubiquitously expressed, while the expressions of ADAR2 and ADAR3 are restricted to certain tissues^11^.

**Figure 2.**
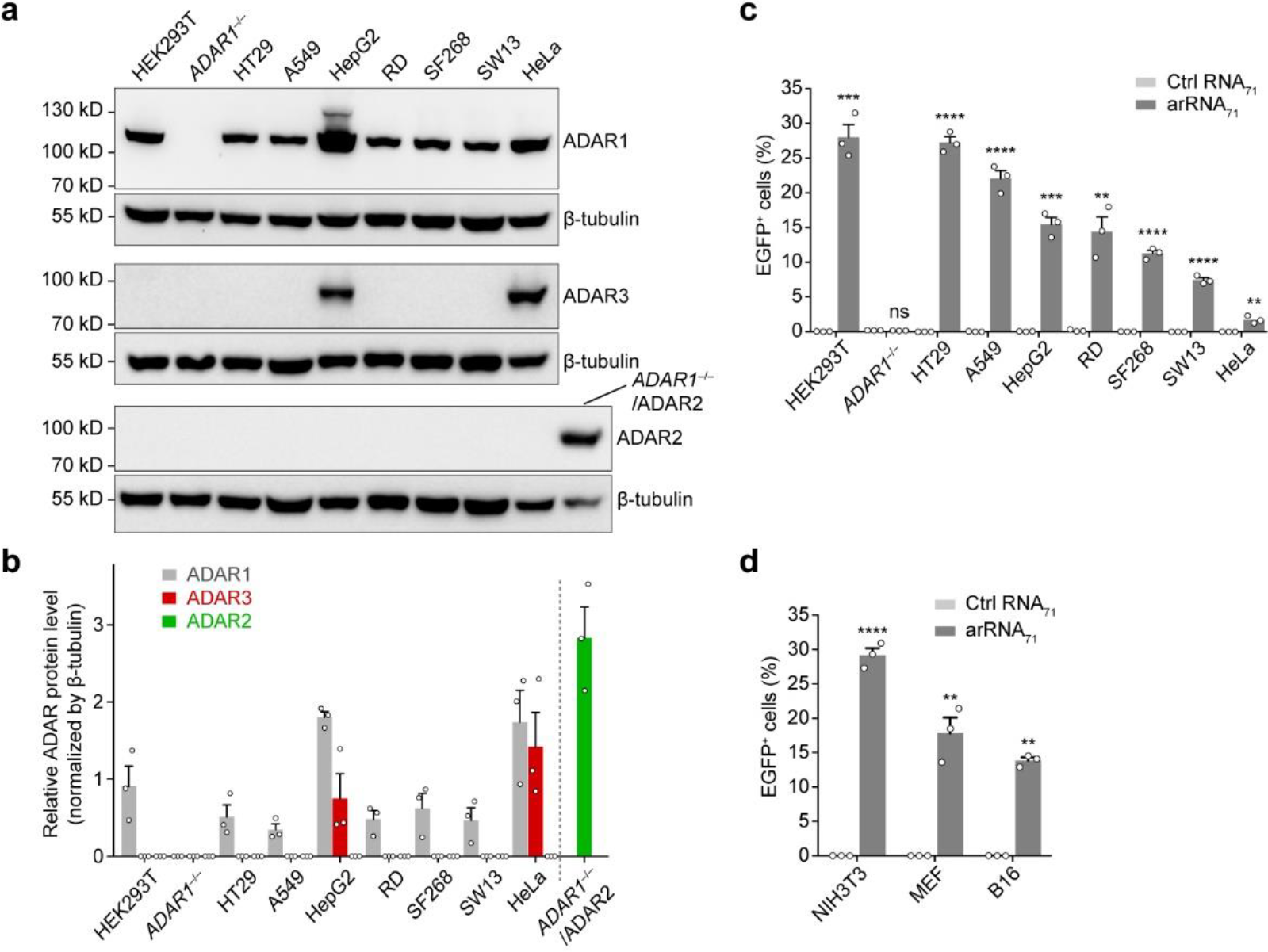
Targeted RNA editing with LEAPER in multiple cell lines. **a**, Western-blot results showing the expression levels of ADAR1, ADAR2 and ADAR3 in indicated human cell lines. β-tubulin was used as a loading control. Data shown is the representative of three independent experiments. *ADAR1*^−/−^/ADAR2 represents ADAR1-knockout HEK293T cells overexpressing ADAR2. **b**, Relative ADAR protein expression levels normalized by β-tubulin expression. **c**, Indicated human cells were transfected with Reporter-1, along with the 71-nt control arRNA (Ctrl RNA_71_) or with the 71-nt targeting arRNA (arRNA_71_) followed by FACS analysis. **d**, Indicated mouse cell lines were analyzed as described in (**c**). EGFP^+^ percentages were normalized by transfection efficiency, which was determined by mCherry^+^. Error bars in (**b**, **c**, **d**) all indicate the mean ± s.e.m. (n = 3); unpaired two-sided Student’s *t*-test, \**P* < 0.05; \*\**P* < 0.01; \*\*\**P* < 0.001; \*\*\*\**P* < 0.0001; ns, not significant.

We then set out to test the editing efficiencies of a re-designed 71-nt arRNA (arRNA_71_) targeting the Reporter-1 (Extended Data Fig. 4a and Supplementary Table 2) in these cell lines. LEAPER worked in all tested cells for this arRNA_71_, albeit with varying efficiencies (Fig. 2c). These results were in agreement with the prior report that the ADAR1/2 protein levels correlate with the RNA editing yield^42^, with the exception of HepG2 and HeLa cells. The suboptimal correlations of editing efficiencies with ADAR1 levels were likely due to the abundant ADAR3 expressions in these two lines (Fig. 2a, b) because it has been reported that ADAR3 plays an inhibitory role in RNA editing^43^. Importantly, LEAPER also worked in three different cell lines of mouse origin (NIH3T3, Mouse Embryonic Fibroblast (MEF) and B16) (Fig. 2d), paving the way for testing its therapeutics potential through animal and disease models. Collectively, we conclude that LEAPER is a versatile tool for wide-spectrum of cell types, and for different organisms.

### Characterization and optimization of LEAPER

To better characterize and optimize LEAPER, we investigated the choices of nucleotide opposite to the adenosine within the UAG triplet of the targeted transcript. In HEK293T cells, Reporter-1-targeting arRNA_71_ showed variable editing efficiencies with a changed triplet (5’-CNA, N denotes one of A/U/C/G) opposite to the targeted UAG (Supplementary Table 2). A-C mismatch resulted in the highest editing efficiency, and the A-G mismatch yielded the least but evident edits (Fig. 3a). We then investigated the preference of nucleotides flanking the A-C mismatch in arRNA. We tested all 16 combinations of 5’ and 3’ neighbor sites surrounding the cytidine (5’-N^1^CN^2^) (Supplementary Table 2), and found that the 3’ neighboring adenosine was required for the efficient editing, while adenosine is the least favorable nucleotide at the 5’ site (Fig. 3b, c). We thus concluded that CCA motif on the arRNA confers the highest editing efficiency targeting the UAG site. It is worthwhile to note that the 3’ neighboring guanosine (5’-N^1^CG) in arRNA showed a dramatic inhibitory effect (Fig. 3b, c).

**Figure 3.**
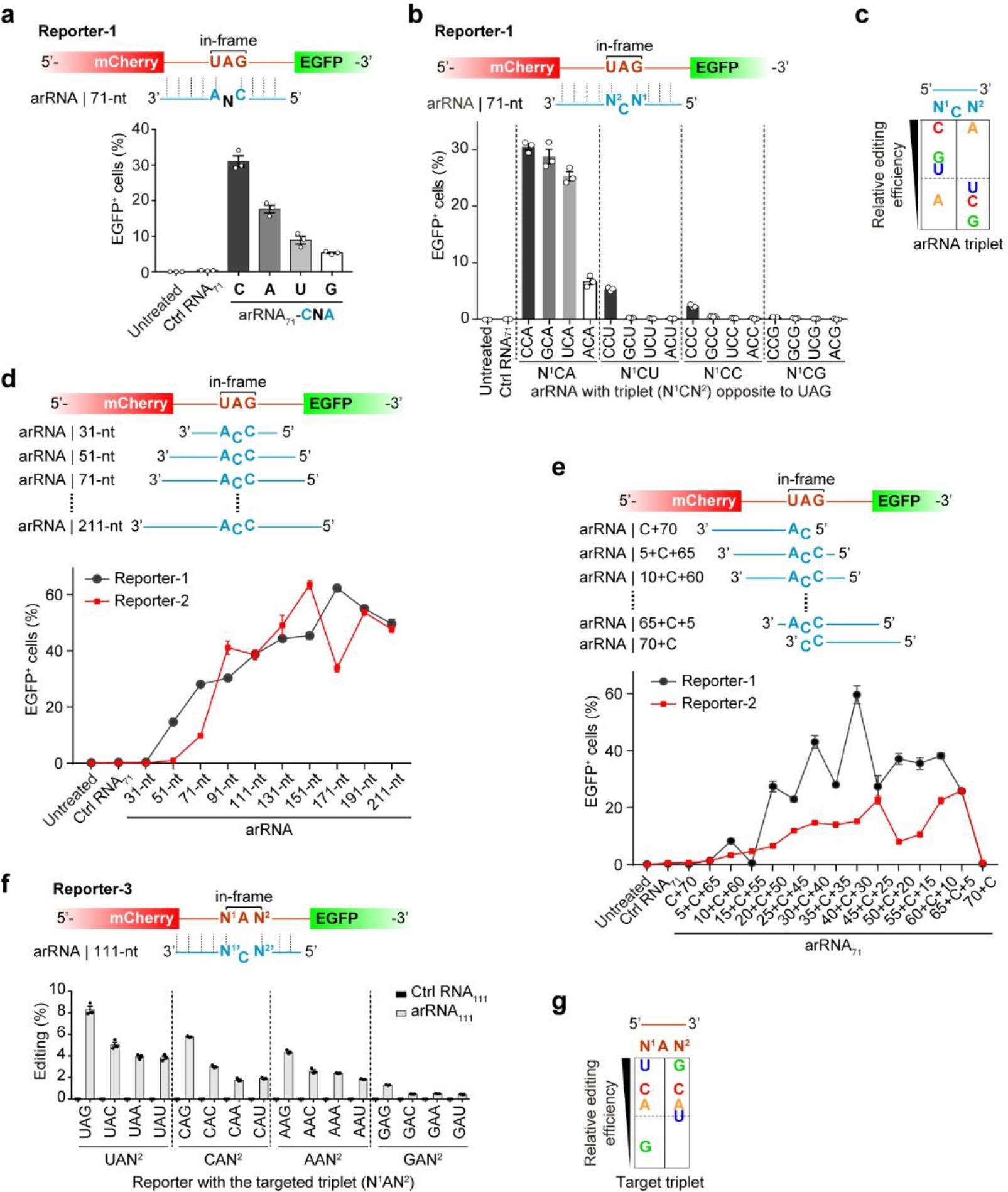
Characterization and optimization of LEAPER. **a**, Top, schematic of the design of arRNAs with changed triplet (5’-CNA, N denotes A, U, C or G) opposite to the target UAG. Bottom, EGFP^+^ percent showing the effects of variable bases opposite to the targeted adenosine on RNA editing efficiency. **b**, Top, the design of arRNAs with changed neighboring bases flanking the cytidine in the A-C mismatch (5’-N^1^CN^2^). Bottom, the effects of 16 different combinations of N^1^CN^2^ on RNA editing efficiency. **c**, Summary of the preference of 5’ and 3’ nearest neighboring sites of the cytidine in the A-C mismatch. **d**, Top, the design of arRNAs with variable length. Bottom, the effect of arRNA length on RNA editing efficiency based on Reporter-1 and Reporter-2. **e**, Top, the design of arRNAs with variable A-C mismatch position. Bottom, the effect of A-C mismatch position on RNA editing efficiency based on Reporter 1 and Reporter-2. **f**, Top, the design of the triplet motifs in the reporter-3 with variable nearest neighboring bases surrounding the targeting adenosine (5’-N^1^AN^2^) and the opposite motif (5’-N^2^CN^1^) on the 111-nt arRNA (arRNA_111_). Bottom, deep sequencing results showing the editing rate on targeted adenosine in the 5’-N^1^AN^2^ motif. **g**, Summary of the 5’ and 3’ base preferences of LEAPER-mediated editing at the Reporter-3. Error bars in (**a**, **b**, **d**, **e** and **f**) all indicate mean values ± s.e.m. (n = 3).

Length of RNA appeared relevant to arRNA efficiency in directing the editing on the targeted transcripts (Extended Data Fig. 1c), consistent with a previous report^42^. To fully understand this effect, we tested arRNAs with variable lengths targeting two different reporter transcripts - Reporter-1 and Reporter-2 (Extended Data Fig. 4a, b). For either reporter targeting, arRNAs of 10 different sizes were designed and tested, ranging from 31-nt to 211-nt, with CCA triplet (for UAG targeting) right in the middle (Supplementary Table 2). Based on the reporter EGFP activities, the length of arRNA correlated positively with the editing efficiency, for both reporters, peaking at 111-to191-nt (Fig. 3d). Although one arRNA_51_ appeared working, 71-nt was the minimal length for arRNA to work for both reporters (Fig. 3d).

Next, we investigated the effect of the A-C mismatch position within an arRNA on editing efficiency. We fixed the lengths of all arRNAs for testing to 71-nt, and slided the UAG-targeting ACC triplet from 5’ to 3’ within arRNAs (Supplementary Table 2). It turned out that placing the A-C mismatch in the middle region resulted in high editing yield, and arRNAs with the mismatch sites close to the 3’ end outperformed those close to the 5’ end in both reporters (Fig. 3e). For convenience, we placed the A-C mismatch at the center of arRNAs for all of our subsequent studies.

We also tested the targeting flexibility of LEAPER and tried to determine whether UAG on target is the only motif subjected to RNA editing. For all 16 triplet combinations (5’-N^1^AN^2^) on Reporter-3 (Extended Data Fig. 4c), we used the corresponding arRNAs with the fixed lengths (111-nt) and ensured the perfect sequencing match for arRNA and the reporter except for the editing site (A-C mismatch) (Fig. 3f and Supplementary Table 2). NGS results showed that all N^1^AN^2^ motifs could be edited. The UAN^2^ and GAN^2^ are the most and the least preferable motifs, respectively (Fig. 3f, g). Collectively, the nearest neighbor preference of the target adenosine is 5’ U>C≈A>G and 3’ G>C>A≈U (Fig. 3g).

### Editing endogenous transcripts using LEAPER

Next, we examined if LEAPER could enable effective editing on endogenous transcripts. Using arRNAs of different lengths, we targeted the UAG motifs in the transcripts of *PPIB*, *KRAS* and *SMAD4* genes, and an UAC motif in *FANCC* gene transcript (Fig. 4a, Supplementary Table 2). Encouragingly, targeted adenosine sites in all four transcripts were edited by their corresponding arRNAs with all four sizes, albeit with variable efficiencies according to NGS results (Fig. 4b). In consistent with our prior observation, longer arRNAs tended to yield higher editing rates. Of note, the 151-nt arRNA^PPIB^ edited ~50% of total transcripts of *PPIB* gene (Fig. 4b). No arRNAs showed RNAi effects on their targeted transcripts (Extended Data Fig. 5a) or ultimate protein level (e.g. KRAS, Extended Data Fig. 5b). Besides, LEAPER is able to achieve desirable editing rate on non-UAN sites (Fig. 4c and Supplementary Table 2), showing the flexibility of LEAPER on editing endogenous transcripts. To further explore the power of LEAPER, we tested whether it could simultaneously target multiple sites. We observed multiplex editing of both *TARDBP* and *FANCC* transcripts by co-expression of two arRNAs (Supplementary Table 2), with the efficiency even higher than those with individual arRNAs (Fig. 4d), indicating that LEAPER is well suited for editing multiple targets in parallel.

**Figure 4.**
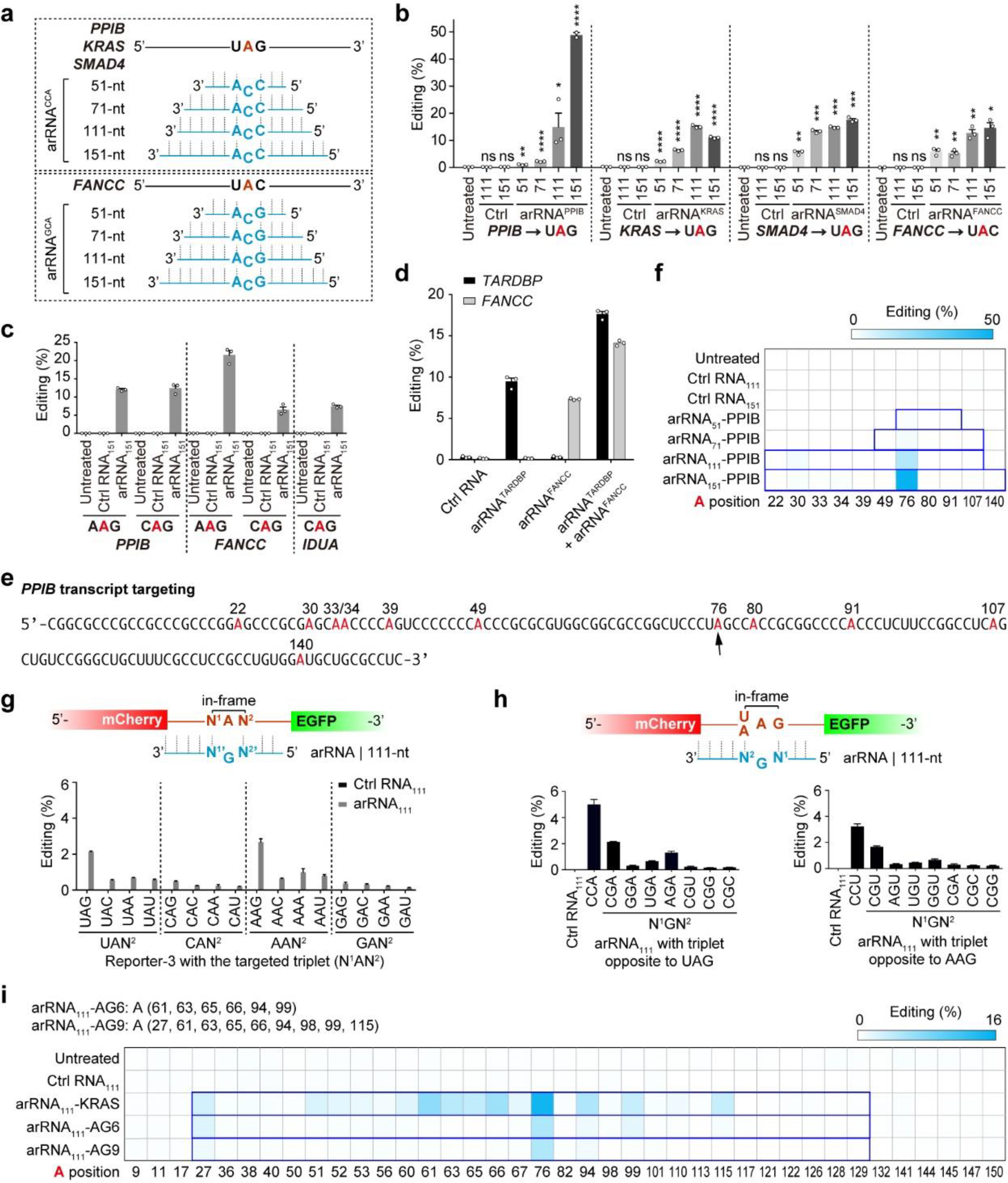
Editing endogenous transcripts with LEAPER. **a**, Schematic of the targeting endogenous transcripts of four disease-related genes (*PPIB*, *KRAS*, *SMAD4* and *FANCC*) and the corresponding arRNAs. **b**, Deep sequencing results showing the editing rate on targeted adenosine of the *PPIB*, *KRAS*, *SMAD4* and *FANCC* transcripts by introducing indicated lengths of arRNAs. **c**, Deep sequencing results showing the editing rate on non-UAN sites of endogenous PPIB, FANCC and IDUA transcripts. **d**, Multiplex editing rate by two 111-nt arRNAs. Indicated arRNAs were transfected alone or were co-transfected into the HEK293T cells. The targeted editing at the two sites was measured from co-transfected cells. **e**, Schematic of the *PPIB* transcript sequence covered by the 151-nt arRNA. The black arrow indicates the targeted adenosine. All adenosines were marked in red. **f**, Heatmap of editing rate on adenosines covered by indicated lengths of arRNAs targeting the *PPIB* gene (marked in bold frame in blue). For the 111-nt arRNA or arRNA_151_-PPIB covered region, the editing rates of A22, A30, A33, and A34 were determined by RNA-seq because of the lack of effective PCR primers for amplifying this region. Otherwise the editing rate was determined by targeted deep-sequencing analysis. **g**, Top, the design of the triplet motifs in the reporter-3 with variable nearest neighboring bases surrounding the targeting adenosine (5’-N^1^AN^2^) and the opposite motif (5’-N^2’^GN^1’^) in the 111-nt arRNA (arRNA_111_). Bottom, deep sequencing results showing the editing rate. **h**, Top, the design of arRNAs with two consecutive mismatches in the 5’-N^1^GN^2^ motif opposite to the 5’-UAG or the 5’-AAG motifs. Deep sequencing results showing the editing rate by an arRNA_111_ with two consecutive mismatches in the 5’-N1GN2 motif opposite to the 5’-UAG motif (bottom left) or the 5’-AAG motif (bottom right). **i**, Heatmap of the editing rate on adenosines covered by engineered arRNA_111_ variants targeting the *KRAS* gene. Data in (**b**, **c**, **d**, **g** and **h**) are presented as the mean ± s.e.m. (n = 3); unpaired two-sided Student’s *t*-test, \**P* < 0.05; \*\**P* < 0.01; \*\*\**P* < 0.001; \*\*\*\**P* < 0.0001; NS, not significant. Data in (**f** and **i**) are presented as the mean (n = 3).

It is noteworthy that ADAR1/2 tend to promiscuously deaminate multiple adenosines in an RNA duplex^44^ and the A-C mismatch is not the only motif to guide the A-to-I switch (Fig. 3a). It is therefore reasonable to assume that all adenosines on target transcripts within the arRNA coverages are subjected to variable levels of editing, major sources of unwanted modifications. The longer the arRNA, the higher the possibility of such off-targets. We therefore examined all adenosine sites within the arRNA covering regions in these targeted transcripts. For *PPIB* transcripts, very little off-target editing was observed throughout the sequencing window for variable sizes of arRNAs (Fig. 4e, f). However, in the cases of targeting *KRAS*, *SMAD4* and *FANCC* genes, multiple off-target edits were detected (Extended Data Fig. 6a-f). For *KRAS* in particular, 11 out of 30 adenosines underwent substantial A to I conversions in the sequencing window of arRNA_111_ (Extended Data Fig. 6a, b).

We next attempted to develop strategies to minimize such unwanted off-target effects. Because an A-G mismatch suppressed editing for UAG targeting (Fig. 3a), we postulated that pairing a guanosine with a non-targeting adenosine might reduce undesirable editing. We then tested the effect of the A-G mismatch on adenosine in all possible triplet combinations (5’-N^1^AN^2^) as in Reporter-3 (Extended Data Fig. 4c and Supplementary Table 2). A-G mismatch indeed decreased the editing on adenosine in all tested targets, except for UAG or AAG targeting (~2%) (Fig. 4g), in comparison with A-C mismatch (Fig. 3f). To further reduce editing rates at unwanted sites, we went on testing the effect of two consecutive mismatches. It turned out that the additional mismatch at the 3’ end nucleotide of the triplet opposite to either UAG or AAG, abolished its corresponding adenosine editing (Fig. 4h and Supplementary Table 2). In light of these findings, we attempted to apply this rule to reduce off-targets in *KRAS* transcripts (Extended Data Fig. 6a). We first designed an arRNA (arRNA_111_-AG6) that created A-G mismatches on all “editing-prone” motifs covered by arRNA_111_ (Fig. 4i, Extended Data Fig. 6a and Supplementary Table 2), including AAU (the 61^st^), UAU (the 63^rd^), UAA (the 65^th^), AAA (the 66^th^), UAG (the 94^th^) and AAG (the 99^th^). This arRNA_111_-AG6 eliminated most of the off-target editing, while maintained an on-target editing rate of ~ 5%. In consistent with the findings in Fig. 4g, the single A-G mismatch could not completely minimize editing in AAG motif (99^th^) (Fig. 4i and Extended Data Fig. 6a). We then added more mismatches on arRNA_111_-AG6, including a dual mismatch (5’-CGG opposite to the targeted motif 5’-AAG), plus three additional A-G mismatches to mitigate editing on the 27^th^, 98^th^ and the 115^th^ adenosines (arRNA_111_-AG9) (Supplementary Table 2). Consequently, we achieved a much improved specificity for editing, without additional loss of editing rate on the targeted site (A76) (Fig. 4i). In summary, engineered LEAPER incorporating additional rules enables efficient and more precise RNA editing on endogenous transcripts.

### RNA editing specificity of LEAPER

In addition to the possible off-target effects within the arRNA-covered dsRNA region, we were also concerned about the potential off-target effects on other transcripts through partial base pairing of arRNA. We then performed a transcriptome-wide RNA-sequencing analysis to evaluate the global off-target effects of LEAPER. Cells were transfected with a Ctrl RNA_151_ or a *PPIB*-specific arRNA (arRNA_151_-PPIB) expressing plasmids before subjected to RNA-seq analysis. We identified six potential off-targets in the Ctrl RNA_151_ group (Fig. 5a) and five in the arRNA_151_-PPIB group (Fig. 5b), and the *PPIB* on-target rate based on NGS analysis was ~37% (Fig. 5b). Further analysis revealed that all sites, except for the two sites from *EIF2AK2* transcripts, were located in either SINE (Alu) or LINE regions (Fig. 5a, b), both are prone to ADAR-mediated editing^45^, suggesting that these off-targets may not be derived from pairing between the target transcripts and the arRNA or control RNA. Of note, two off-targeting transcripts, *WDR73* and *SMYD4*, appeared in both groups, suggesting they are unlikely sequence-dependent RNA editing. Indeed, minimum free energy analysis suggested that all these possible off-target transcripts failed to form a stable duplex with either Ctrl RNA_151_ or arRNA_151_-PPIB (Fig. 5c). To further test if arRNA generates sequence-dependent off-targets, we selected potential off-target sites by comparing sequence similarity using NCBI BLAST for both arRNA_151_-PPIB and arRNA_111_-FANCC. *TRAPPC12* transcripts for arRNA_151_-PPIB and three sites in the *ST3GAL1*, *OSTM1-AS1* and *EHD2* transcripts for arRNA_111_-FANCC were top candidates (Fig. 5d and Extended Data Fig. 7a). NGS analysis revealed that no editing could be detected in any of these predicted off-target sites (Fig. 5d and Extended Data Fig. 7b). These results indicate that LEAPER empowers efficient editing at the targeted site, while maintaining transcriptome-wide specificity without detectable sequence-dependent off-target edits.

**Figure 5.**
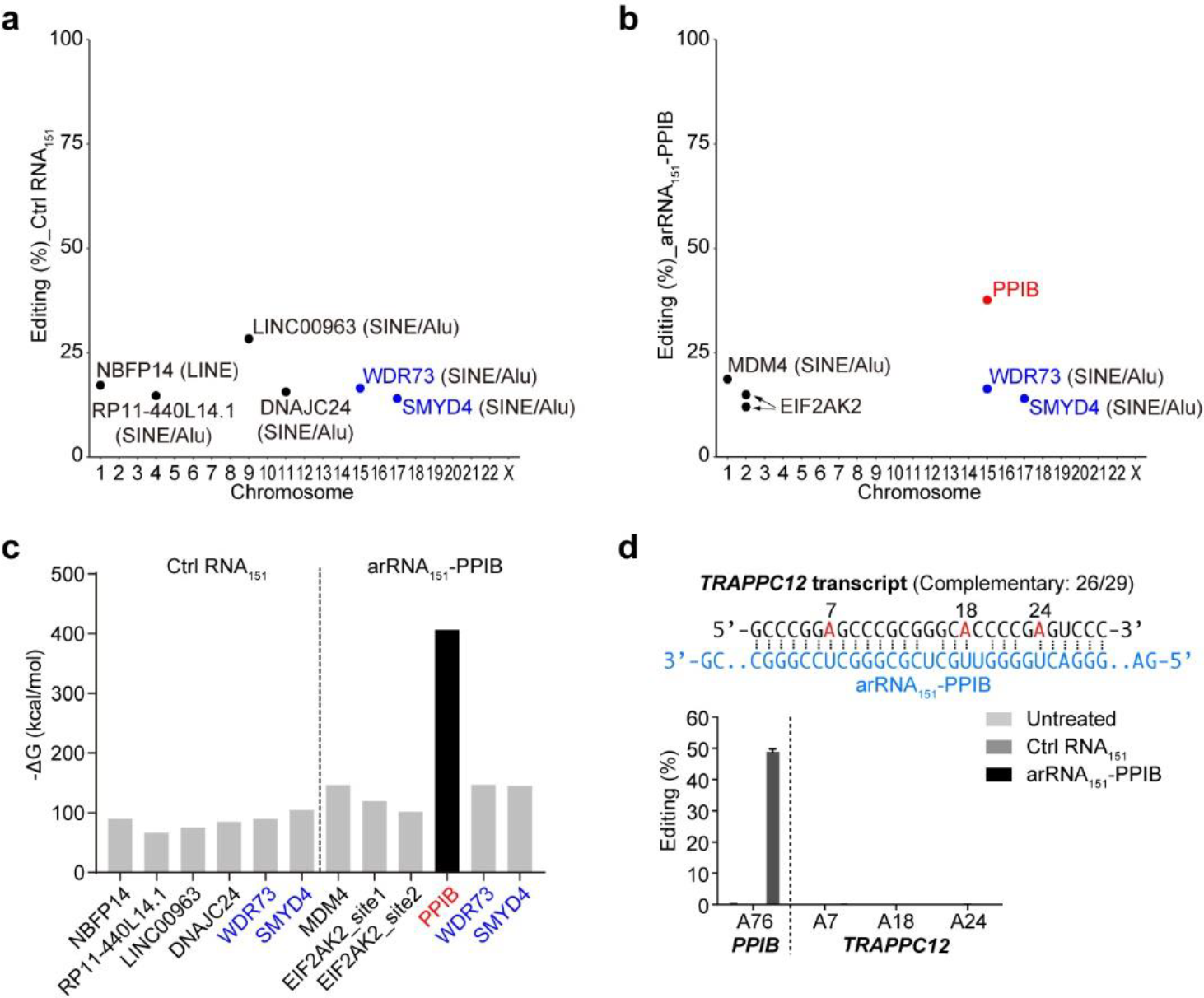
Transcriptome-wide specificity of RNA editing by LEAPER. **a** and **b**, Transcriptome-wide off-targeting analysis of Ctrl RNA_151_ and arRNA_151_-PPIB. The on-targeting site (PPIB) is highlighted in red. The potential off-target sites identified in both Ctrl RNA and PPIB-targeting RNA groups are labeled in blue. **c**, The predicted annealing affinity between off-target sites and the corresponding Ctrl RNA_151_ or arRNA_151_-PPIB. The minimum free energy (ΔG) of double-stranded RNA formed by off-target sites (150-nt upstream and downstream of the editing sites) and the corresponding Ctrl RNA_151_ or arRNA_151_-PPIB was predicted with RNAhybrid, an online website tool. **d**, Top, schematic of the highly complementary region between arRNA_151_-PPIB and the indicated potential off-target sites, which were predicted by searching homologous sequences through NCBI-BLAST. Bottom, Deep sequencing showing the editing rate on the on-target site and all predicted off-target sites of arRNA_151_-PPIB. Data are presented as the mean ± s.e.m. (n = 3).

### Safety assessment of LEAPER in mammalian cells

Because arRNAs rely on endogenous ADAR proteins for editing on target transcripts, we wondered if the addition of exogenous arRNAs affects native RNA editing events by occupying too much of ADAR1 or ADAR2 proteins. Therefore, we analyzed the A-to-I RNA editing sites shared by mock group and arRNA_151_-PPIB group from the transcriptome-wide RNA-sequencing results, and the comparison between the mock group and Ctrl RNA_151_ group was also analyzed. Neither Ctrl RNA_151_ group nor arRNA_151_-PPIB group showed a significant difference compared to the mock group (Fig. 6a, b), indicating that LEAPER had little impact on the normal function of endogenous ADAR1 to catalyze the native A-to-I editing events.

**Figure 6.**
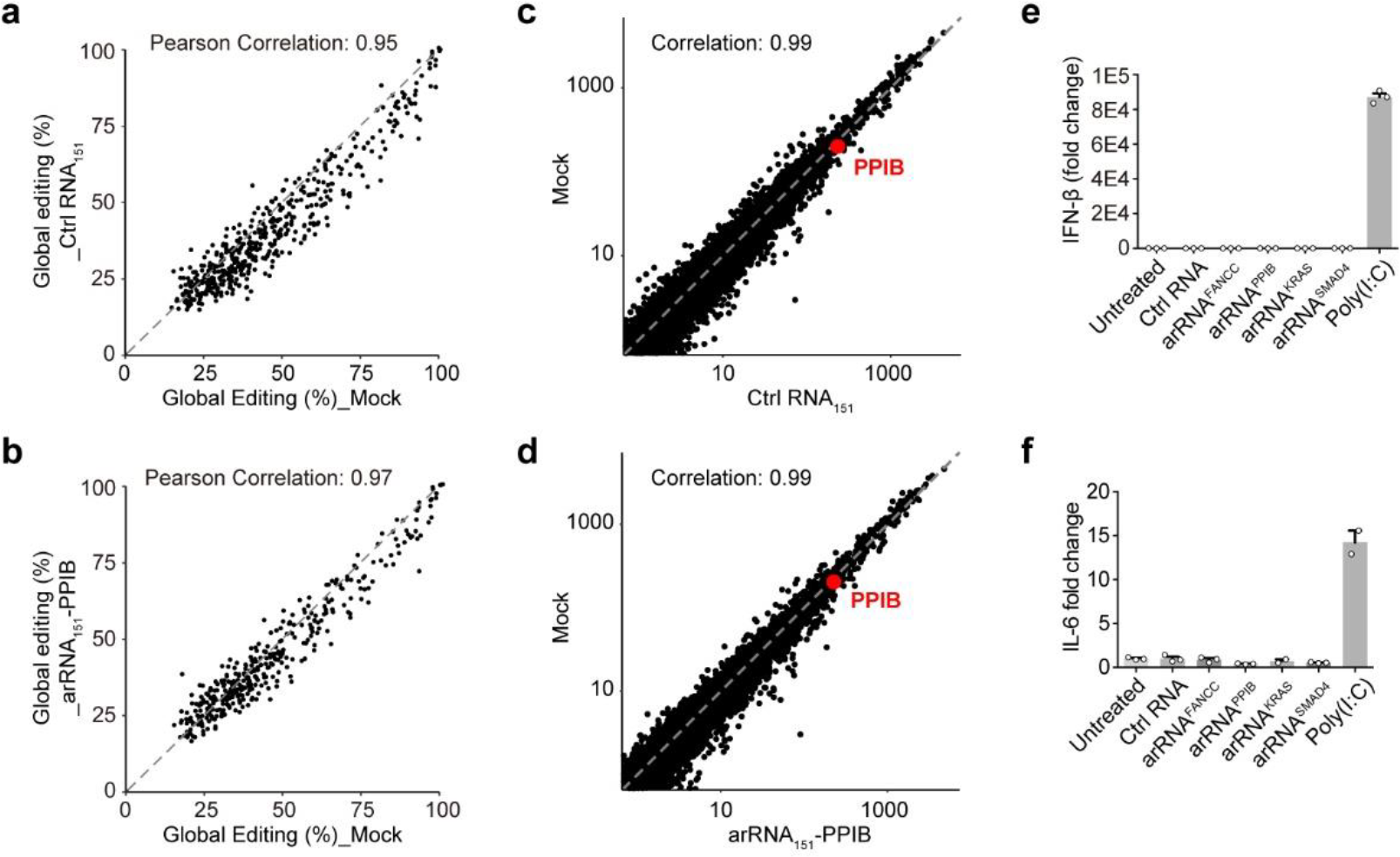
Safety evaluation of applying LEAPER in mammalian cells. **a** and **b**, Transcriptome-wide analysis of the effects of Ctrl RNA_151_ (**a**) arRNA_151_-PPIB (**b**) on native editing sites by transcriptome-wide RNA-sequencing. Pearson’s correlation coefficient analysis was used to assess the differential RNA editing rate on native editing sites. **c** and **d**, Differential gene expression analysis of the effects of Ctrl RNA_151_ (**c**) arRNA_151_-PPIB (**d**) with RNA-seq data at the transcriptome level. Pearson’s correlation coefficient analysis was used to assess the differential gene expression. **e** and **f**, Effect of arRNA transfection on innate immune response. The indicated arRNAs or the poly(I:C) were transfected into HEK293T cells. Total RNA was then analyzed using quantitative PCR to determine expression levels of IFN-β (**e**) and IL-6 (**f**). Data (**e** and **f**) are presented as the mean ± s.e.m. (n = 3).

Meanwhile, we performed differential gene expression analysis using RNA-seq data to verify whether arRNA affects global gene expression. We found that neither Ctrl RNA_151_ nor arRNA_151_-PPIB affected the global gene expression in comparison with the mock group (Fig. 6c, d). In consistent with our prior observation (Extended Data Fig. 5a), arRNAs did not show any RNAi effect on the expression of *PPIB* (Fig. 6c, d).

Considering that the arRNA forms RNA duplex with the target transcript and that RNA duplex might elicit innate immune response, we investigated if the introduction of arRNA has such an effect. To test this, we selected arRNAs targeting four gene transcripts that had been proven effective. We did not observe any mRNA induction of interferon-β (IFN-β) (Fig. 6e) or interleukin-6 (IL-6) (Fig. 6f), which are two hallmarks of innate immune activation. As a positive control, a synthetic analog of double-stranded RNA - poly(I:C) induced strong IFN-β and IL-6 expression (Fig. 6e, f). LEAPER does not seem to induce immunogenicity in target cells, a feature important for safe therapeutics.

### Recovery of transcriptional regulatory activity of p53 by LEAPER

Now that we have established a novel method for RNA editing without the necessity of introducing foreign proteins, we attempted to demonstrate its therapeutic utility. We first targeted the tumor suppressor gene *TP53*, which is known to play a vital role in the maintenance of cellular homeostasis, but undergo frequent mutations in >50% of human cancers^46^. The c.158G>A mutation in *TP53* is a clinically-relevant nonsense mutation (Trp53Ter), resulting in a non-functional truncated protein^46^. We designed one arRNA_111_ and two alternative arRNAs (arRNA_111_-AG1 and arRNA_111_-AG4) (Supplementary Table 2), all targeting *TP53*^W53X^ transcripts (Fig. 7a), with the latter two being designed to minimize potential off-targets. We generated HEK293T *TP53*^−/−^ cell line to eliminate the effects of native p53 protein. All three forms of *TP53*^W53X^-targeting arRNAs converted ~25-35% of *TP53*^W53X^ transcripts on the mutated adenosine site (Fig. 7b), with variable reductions of unwanted edits for arRNA_111_-AG1 and arRNA_111_-AG4 (Extended Data Fig. 8). Western blot showed that arRNA_111_, arRNA_111_-AG1 and arRNA_111_-AG4 could all rescue the production of full-length p53 protein based on the *TP53*^W53X^ transcripts in HEK293T *TP53*^−/−^ cells, while the Ctrl RNA_111_ could not (Fig. 7c). To verify whether the repaired p53 proteins are fully functional, we tested the transcriptional regulatory activity of p53 with a p53-luciferase *cis*-reporting system^47,48^. All three versions of arRNAs could restore p53 activity, and the optimized version arRNA_111_-AG1 performed the best (Fig. 7d). In conclusion, we demonstrated that LEAPER is capable of repairing the cancer-relevant pre-mature stop codon of *TP53* and restoring its function.

**Figure 7.**
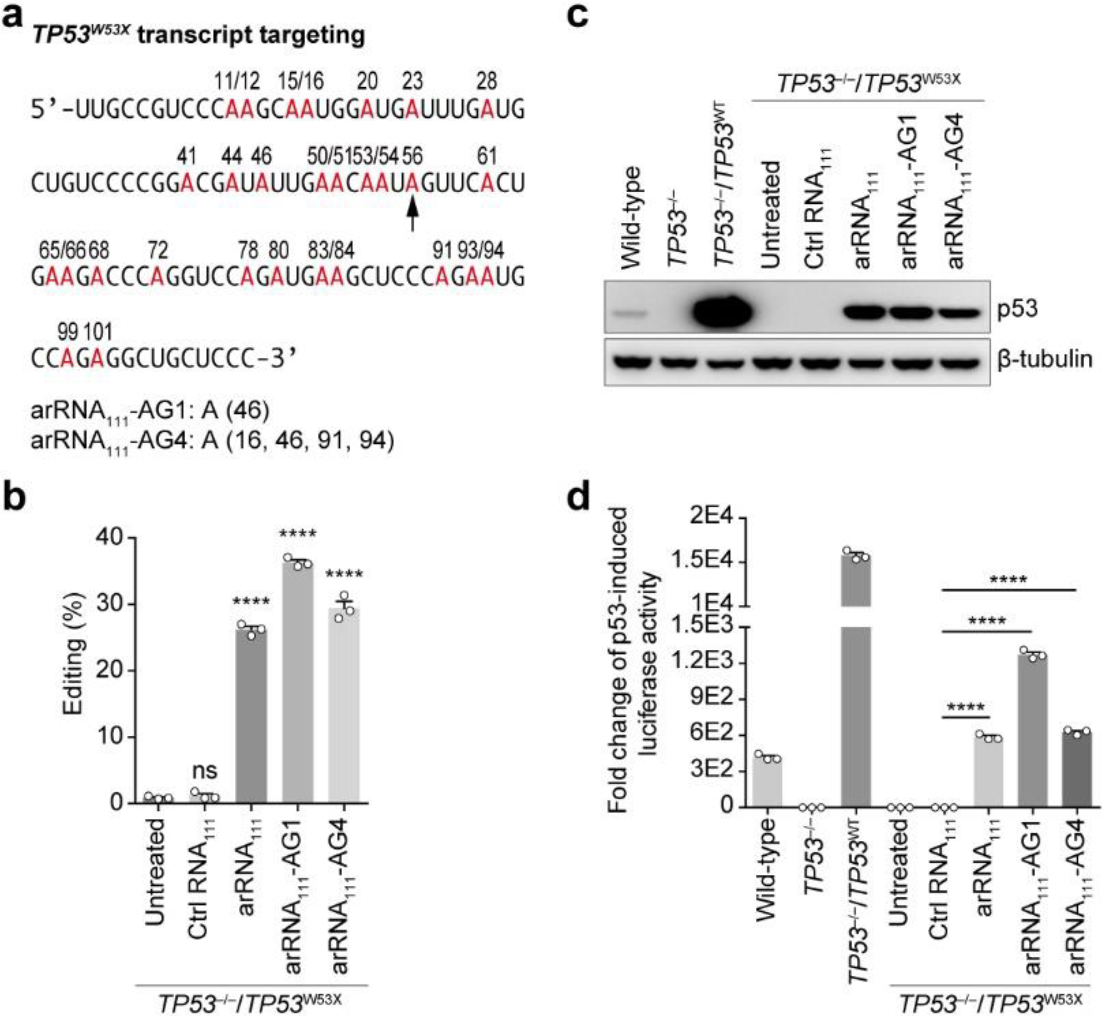
Recovery of transcriptional regulatory activity of mutant *TP53*^W53X^ by LEAPER. **a**, Top, Schematic of the *TP53* transcript sequence covered by the 111-nt arRNA containing c.158G>A clinical-relevant non-sense mutation (Trp53Ter). The black arrow indicates the targeted adenosine. All adenosines were marked in red. Bottom, the design of two optimized arRNAs targeting *TP53*^W53X^ transcripts with A-G mismatch on A^46th^ for arRNA_111_-AG1, and on A^16th^, A^46th^, A^91th^ and A^94th^ together for arRNA_111_-AG4 to minimize the potential off-targets on “editing-prone” motifs. **b**, Deep sequencing results showing the targeted editing on *TP53*^W53X^ transcripts by arRNA_111_, arRNA_111_-AG1 and arRNA_111_-AG4. **c**, Western blot showing the recovered production of full-length p53 protein from the *TP53*^W53X^ transcripts in the HEK293T *TP53*^−/−^ cells. **d**, Detection of the transcriptional regulatory activity of restored p53 protein using a p53-Firefly-luciferase reporter system, normalized by co-transfected Renilla-luciferase vector. Data (**b**, **c** and **d**) are presented as the mean ± s.e.m. (n = 3); unpaired two-sided Student’s *t*-test, \**P* < 0.05; \*\**P* < 0.01; \*\*\**P* < 0.001; \*\*\*\**P* < 0.0001; ns, not significant.

### Corrections of pathogenic mutations by LEAPER

We next investigated whether LEAPER could be used to correct more pathogenic mutations. Aiming at clinically relevant mutations from six pathogenic genes, *COL3A1* of Ehlers-Danlos syndrome, *BMPR2* of Primary pulmonary hypertension, *AHI1* of Joubert syndrome, *FANCC* of Fanconi anemia, *MYBPC3* of Primary familial hypertrophic cardiomyopathy and *IL2RG* of X-linked severe combined immunodeficiency, we designed 111-nt arRNAs for each of these genes carrying corresponding pathogenic G>A mutations (Extended Data Fig. 9 and Supplementary Tables 2, 3). By co-expressing arRNA/cDNA pairs in HEK293T cells, we identified significant amounts of target transcripts with A>G corrections in all tests (Fig. 8). Because G>A mutations account for nearly half of known disease-causing point mutations in humans^10,49^, the A>G conversion by LEAPER may offer immense opportunities for therapeutics.

**Figure 8.**
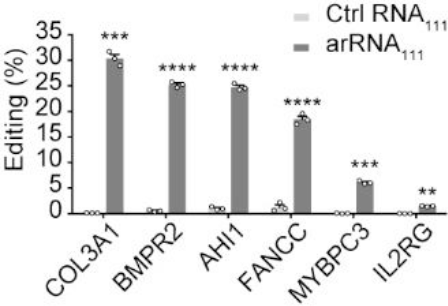
Correction of pathogenic mutations by LEAPER. A to I correction of disease-relevant G>A mutation from ClinVar data by the corresponding 111-nt arRNA, targeting clinical-related mutations from six pathogenic genes as indicated (Extended Data Fig. 9 and Supplementary Tables 2, 3). Data are presented as the mean ± s.e.m. (n = 3); unpaired two-sided Student’s *t*-test, \**P* < 0.05; \*\**P* < 0.01; \*\*\**P* < 0.001; \*\*\*\**P* < 0.0001; ns, not significant.

### RNA editing in multiple human primary cells by LEAPER

To further explore the clinical utility of LEAPER, we set out to test the method in multiple human primary cells. First, we tested LEAPER in human primary pulmonary fibroblasts and human primary bronchial epithelial cells with 151-nt arRNA (Supplementary Table 2) to edit the Reporter-1 (Extended Data Fig. 4a). 35-45% of EGFP positive cells could be obtained by LEAPER in both human primary cells (Fig. 9a). We then tested LEAPER in editing endogenous gene *PPIB* in these two primary cells and human primary T cells, and found that arRNA_151_-PPIB could achieve >40%, >80% and >30% of editing rates in human primary pulmonary fibroblasts, primary bronchial epithelial cells (Fig. 9b) and primary T cells (Fig. 9c), respectively. The high editing efficiency of LEAPER in human primary cells is particularly encouraging for its potential application in therapeutics.

**Figure 9.**
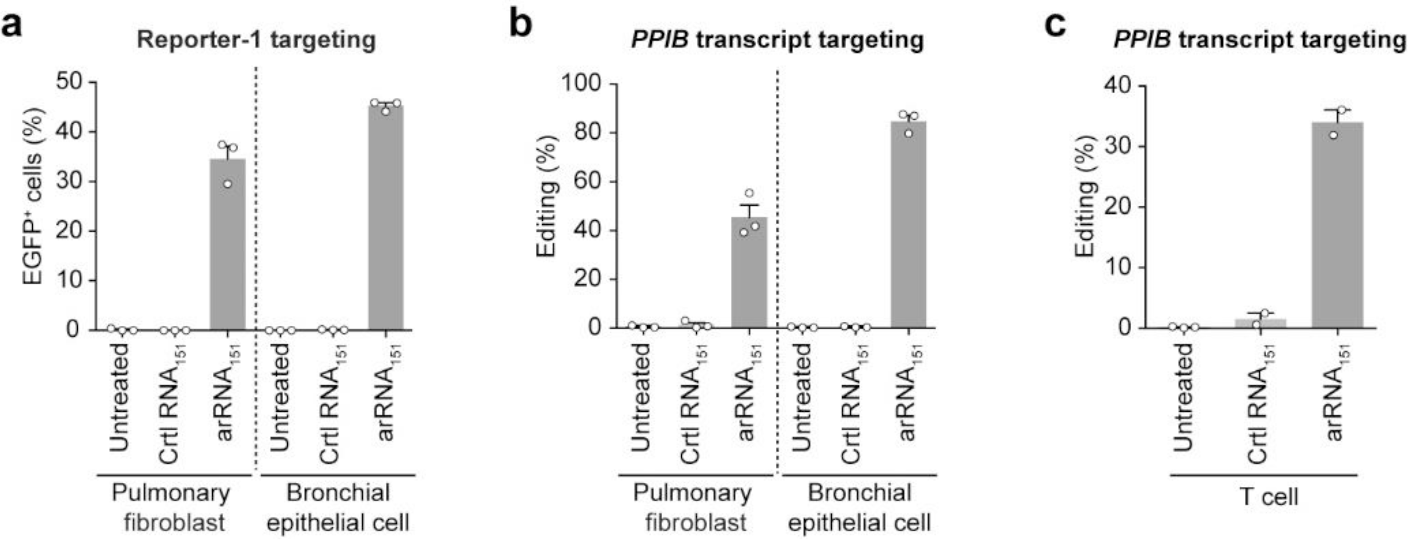
RNA editing in multiple human primary cells by LEAPER. **a**, Quantification of the EGFP positive (EGFP^+^) cells induced by LEAPER-mediated RNA editing. Human primary pulmonary fibroblasts and human primary bronchial epithelial cells were transfected with Reporter-1, along with the 151-nt control RNA (Ctrl RNA_151_) or the 151-nt targeting arRNA (arRNA_151_) followed by FACS analysis. **b** and **c**, Deep sequencing results showing the editing rate on *PPIB* transcripts in human primary pulmonary fibroblasts, human primary bronchial epithelial cells (**b**), and human primary T cells (**c**). Data in **a**, **b** and Untreated group (**c**) are presented as the mean ± s.e.m. (n = 3); data of Ctrl RNA_151_ and arRNA_151_ (**c**) are presented as the mean ± s.e.m. (n = 2).

### Efficient editing by lentiviral expression and chemical synthesis of arRNAs

We then investigated if LEAPER could be delivered by more clinically-relevant methods. We first tested the effect of arRNA through lentivirus-based expression. Reporter-1-targeting arRNA_151_ induced strong EGFP expression in more than 40% of total cells harboring the Reporter-1 in HEK293T cells 2 days post infection (dpi). At 8 dpi, the EGFP ratio maintained at a comparable level of ~38% (Fig. 10a and Supplementary Table 2), suggesting that LEAPER could be tailored to therapeutics that require continuous administration. For native gene editing, we delivered *PPIB*-targeting arRNA_151_ through lentiviral transduction in HEK293T cells and observed over 6% of target editing at 6 dpi (Fig. 10b).

**Figure 10.**
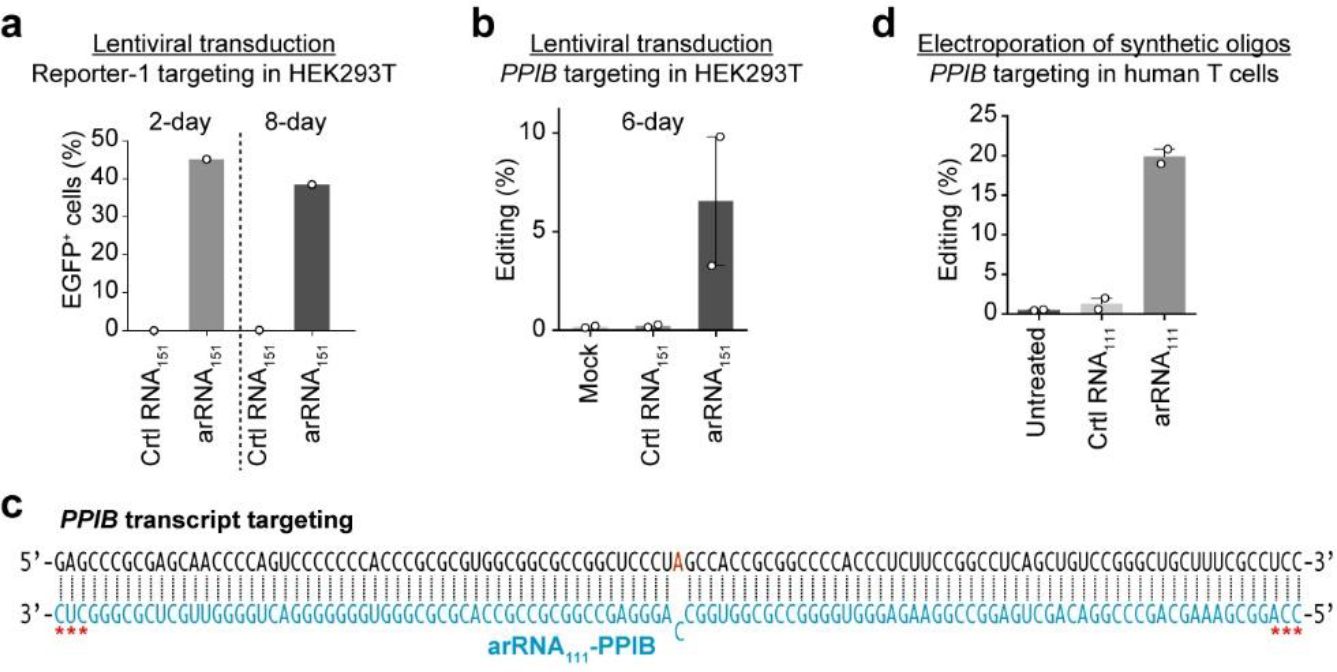
Targeted editing by lentiviral transduction of arRNA and electroporation of synthesized arRNA oligonucleotides. **a**, Quantification of the EGFP^+^ cells. HEK293T cells stably expressing the Repoter-1 were infected with lentivirus expressing 151-nt of Ctrl RNA or the targeting arRNA. FACS analyses were performed 2 days and 8 days post infection. The ratios of EGFP^+^ cells were normalized by lentiviral transduction efficiency (BFP^+^ ratios). **b**, Deep sequencing results showing the editing rate on the *PPIB* transcripts upon lentiviral transduction of 151-nt arRNAs into HEK293T cells. **c**, Schematic of the *PPIB* sequence and the corresponding 111-nt targeting arRNA. *(in red) represents nucleotide with 2’-O-methyl and phosphorothioate linkage modifications. **d**, Deep sequencing results showing the editing rate on the *PPIB* transcripts upon electroporation of 111-nt synthetic arRNA oligonucleotides into human primary T cells.

We next tested synthesized arRNA oligonucleotides and electroporation delivery method for LEAPER. The 111-nt arRNA targeting *PPIB* transcripts as well as Ctrl RNA were chemically synthesized with 2’-O-methyl and phosphorothioate linkage modifications at the first three and last three residues of arRNAs (Fig. 10c). After introduced into T cells through electroporation, arRNA_111_-PPIB oligos achieved ~20% of editing on *PPIB* transcripts (Fig. 10d), indicating that LEAPER holds promise for the development of oligonucleotide drugs.

### Restoration of α-L-iduronidase activity in Hurler syndrome patient-derived primary fibroblast by LEAPER

Finally, we examined the potential of LEAPER in treating a monogenic disease - Hurler syndrome, the most severe subtype of Mucopolysaccharidosis type I (MPS I) due to the deficiency of α-L-iduronidase (IDUA), a lysosomal metabolic enzyme responsible for the degradation of mucopolysaccharides^50^. We chose a primary fibroblast GM06214 that was originally isolated from Hurler syndrome patient. The GM06214 cells contain a homozygous TGG>TAG mutation in exon 9 of the *IDUA* gene, resulting in a Trp402Ter mutation in the protein. We designed two versions of arRNAs by synthesized RNA oligonucleotides with chemical modifications, arRNA_111_-IDUA-V1 and arRNA_111_-IDUA-V2, targeting the mature mRNA and the pre-mRNA of *IDUA*, respectively (Fig. 11a and Supplementary Table 2). After introduction of arRNA_111_-IDUA-V1 or arRNA_111_-IDUA-V2 into GM06214 cells via electroporation, we measured the targeted RNA editing rates via NGS analysis and the catalytic activity of α-L-iduronidase with 4-MU-α-L-iduronidase substrate at different time points. Both arRNA_111_-IDUA-V1 and arRNA_111_-IDUA-V2 significantly restored the IDUA catalytic activity in *IDUA*-deficient GM06214 cells progressively with time after electroporation, and arRNA_111_-IDUA-V2 performed much better than arRNA_111_-IDUA-V1, while no α-L-iduronidase activity could be detected in three control groups (Fig. 11b).

**Figure 11.**
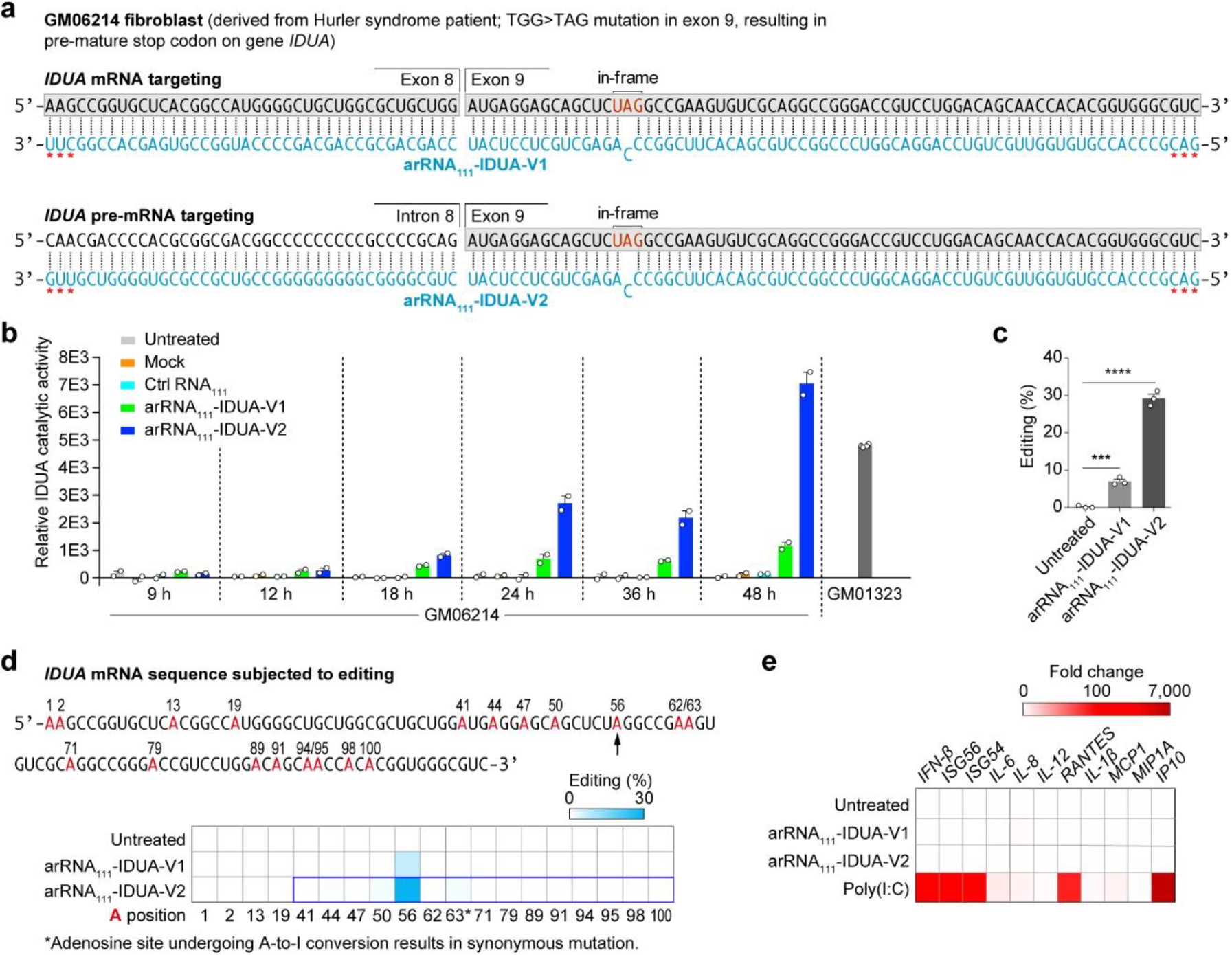
Restoration of α-L-iduronidase activity in Hurler syndrome patient-derived primary fibroblast by LEAPER. **a**, Top, genetic information of pathogenic mutation in patient-derived fibroblast GM06214; Medium, schematic of the *IDUA* mature mRNA sequence of GM06214 cells (Black) containing a homozygous TGG>TAG mutation in exon 9 of the *IDUA* gene (Trp402Ter), and the corresponding 111-nt targeting arRNA_111_-IDUA-V1 (Blue); Bottom, schematic of the *IDUA* pre-mRNA sequence of GM06214 cells (Black) and the corresponding 111-nt targeting arRNA_111_-IDUA-V2 (Blue). *(in red) represents nucleotides with 2’-O-methyl and phosphorothioate linkage modifications. **b**, Measuring the catalytic activity of α-L-iduronidase with 4-methylumbelliferyl α-L-iduronidase substrate at different time points. Data are presented as the mean ± s.e.m. (n = 2). **c**, Deep sequencing results showing the targeted editing rate on *IDUA* transcripts in GM06214 cells, 48 hours post electroporation. **d**, Top, schematic of the *IDUA* transcript sequence covered by the 111-nt arRNAs. The arrow indicates the targeted adenosine. All adenosines were marked in red. Bottom, a heatmap of editing rate on adenosines covered by indicated arRNAs in the *IDUA* transcript (marked in the bold frame in blue). **e**, Quantitative PCR showing the expressions of type I interferon, interferon-stimulated genes, and pro-inflammatory genes upon arRNA or poly(I:C) electroporation. Data are presented as the mean (n = 3).

To further evaluate the extent to which the restored IDUA activity in GM06214 by LEAPER relieves the Hurler syndrome, we examined the IDUA activity in GM01323 cells, another primary fibroblasts from patient with Scheie syndrome, a much milder subtype of MPS I than Hurler syndrome due to the remnant IDUA activity resulting from heterozygous genotype on *IDUA* gene. We found that the catalytic activity of IDUA in GM06214 cells harbouring arRNA_111_-IDUA-V2 was higher than GM01323 cells 48 hr post electroporation (Fig. 11b). Consistent with these results, NGS analysis indicated that arRNA_111_-IDUA-V2 converted nearly 30% of A to I editing, a much higher rate than arRNA_111_-IDUA-V1 (Fig. 11c). Further analysis revealed that minimal unwanted edits were detected within the arRNA covered regions of *IDUA* transcripts (Fig. 11d). Importantly, LEAPER did not trigger immune responses in primary cells as we demonstrated that, unlike the RNA duplex poly(I:C) serving as a positive control, neither arRNA_111_-IDUA-V1 nor arRNA_111_-IDUA-V2 induced expressions of a panel of genes involved in type-I interferon and pro-inflammatory responses (Fig. 11e). These results showed the therapeutic potential of LEAPER in targeting certain monogenetic diseases.

## Discussion

In this report, we showed that expression of a linear arRNA with adequate length is capable of guiding endogenous ADAR proteins to edit adenosine to inosine on the targeted transcripts. This system, referred to as LEAPER, utilizes endogenous ADAR proteins to achieve programmable nucleic acid editing, thus possessing advantages over existing approaches.

The rare quality of LEAPER is its simplicity because it only relies on a small size of RNA molecule to direct the endogenous proteins for RNA editing. This is reminiscent of RNAi, in which a small dsRNA could invoke native mechanism for targeted RNA degradation^51^. Because of the small size, arRNA could be readily delivered by a variety of viral and non-viral vehicles. Different from RNAi, LEAPER catalyzes the precise A to I switch without generating cutting or degradation of targeted transcripts (Extended Data Fig. 5a). Although the length requirement for arRNA is longer than RNAi, it neither induces immune-stimulatory effects at the cellular level (Fig. 6e, f and Fig. 11e) nor affects the function of endogenous ADAR proteins (Fig. 6a, b), making it a safe strategy for RNA targeting. Remarkably, it has been reported that ectopic expression of ADAR proteins or their catalytic domains induces substantial global off-target edits^32^ and possibly triggers cancer^31^.

Recently, several groups reported that cytosine base editor could generate substantial off-target single-nucleotide variants in mouse embryos, rice or human cell lines due to the expression of an effector protein, which illustrates the advantage of LEAPER for potential therapeutic application^52–54^. Gratifyingly, LEAPER empowers efficient editing while elicits rare global off-target editing (Fig. 5 and Extended Data Fig. 7). In addition, LEAPER could minimize potential immunogenicity or surmount delivery obstacles commonly shared by other methods that require the introduction of foreign proteins.

For LEAPER, we would recommend using arRNA with a minimal size above 70-nt to achieve desirable activity. In the native context, ADAR proteins non-specifically edit Alu repeats which have a duplex of more than 300-nt^55^. Of note, Alu repeats form stable intramolecular duplex, while the LEAPER results in an intermolecular duplex between arRNA and mRNA or pre-mRNA, which is supposed to be less stable and more difficult to form. Therefore, we hypothesized that an RNA duplex longer than 70-nt is stoichiometrically important for recruiting or docking ADAR proteins for effective editing. Indeed, longer arRNA resulted in higher editing yield in both ectopically expressed reporters and endogenous transcripts (Fig. 3d and Fig. 4b). However, because ADAR proteins promiscuously deaminate adenosine base in the RNA duplex, longer arRNA may incur more off-targets within the targeting window.

While LEAPER could effectively target native transcripts, their editing efficiencies and off-target rates varied. For *PPIB* transcript-targeting, we could convert 50% of targeted adenosine to inosine without evident off-targets within the covering windows (Fig. 4b, f). The off-targets became more severe for other transcripts. We have managed to reduce off-targets such as introducing A-G mismatches or consecutive mismatches to repress undesired editing. However, too many mismatches could decrease on-target efficiency. Weighing up the efficiency and potential off-targets, we would recommend arRNA with the length ranging from 100-to 150-nt for editing on endogenous transcripts. If there is a choice, it’s better to select regions with less adenosine to minimize the chance of unwanted edits. Encouragingly, we have not detected any off-targets outside of the arRNA-targeted-transcript duplexes (Fig. 5).

We have optimized the design of the arRNA to achieve improved editing efficiency and demonstrated that LEAPER could be harnessed to manipulate gene function or correct pathogenic mutation. We have also shown that LEAPER is not limited to only work on UAG, instead that it works with possibly any adenosine regardless of its flanking nucleotides (Fig. 3f, g and Fig. 4c). Such flexibility is advantageous for potential therapeutic correction of genetic diseases caused by certain single point mutations. Interestingly, in editing the *IDUA* transcripts, the arRNA targeting pre-mRNA is more effective than that targeting mature RNA, indicating that nuclei are the main sites of action for ADAR proteins and LEAPER could be leveraged to manipulate splicing by modifying splice sites within pre-mRNAs. What’s more, LEAPER has demonstrated high efficiency for simultaneously targeting multiple gene transcripts (Fig. 4d). This multiplexing capability of LEAPER might be developed to cure certain polygenetic diseases in the future.

It is beneficial to perform genetic correction at the RNA level. First, editing on targeted transcripts would not permanently change the genome or transcriptome repertoire, making RNA editing approaches safer for therapeutics than means of genome editing. In addition, transient editing is well suited for temporal control of treating diseases caused by occasional changes in a specific state. Second, LEAPER and other RNA editing methods would not introduce DSB on the genome, avoiding the risk of generating undesirable deletions of large DNA fragments^37^. DNA base editing methods adopting nickase Cas9 could still generate indels in the genome^8^. Furthermore, independent of native DNA repair machinery, LEAPER should also work in post-mitosis cells such as cerebellum cells with high expression of ADAR2^11^.

We have demonstrated that LEAPER could apply to a broad spectrum of cell types such as human cell lines (Fig. 2c), mouse cell lines (Fig. 2d) and human primary cells including primary T cells (Fig. 9 and Fig. 10d). Efficient editing through lentiviral delivery or synthesized oligo provides increased potential for therapeutic development (Fig. 10). Moreover, LEAPER could produce phenotypic or physiological changes in varieties of applications including recovering the transcriptional regulatory activity of p53 (Fig. 7), correcting pathogenic mutations (Fig. 8), and restoring the α-L-iduronidase activity in Hurler syndrome patient-derived primary fibroblasts (Fig. 11). It can thus be envisaged that LEAPER has enormous potential in disease treatment.

While our manuscript was under revision, Stafforst and colleagues reported a new and seemingly similar RNA editing method, named RESTORE, which works through recruiting endogenous ADARs using synthetic antisense oligonucleotides^56^. The fundamental difference between RESTORE and LEAPER lies in the distinct nature of the guide RNA for recruiting endogenous ADAR. The guide RNA of RESTORE is limited to chemosynthetic antisense oligonucleotides (ASO) depending on complex chemical modification, while arRNA of LEAPER can be generated in a variety of ways, chemical synthesis and expression from viral or non-viral vectors (Fig. 10 and Fig. 11). Importantly, being heavily chemically modified, ASOs is restricted to act transiently in disease treatment. In contrast, arRNA could be produced through expression, a feature particularly important for the purpose of constant editing.

There are still rooms for improvements regarding LEAPER’s efficiency and specificity. Because LEAPER relies on the endogenous ADAR, the expression level of ADAR proteins in target cells is one of the determinants for successful editing. According to previous report^57^ and our observations (Fig. 2a, b), the ADAR1^p110^ is ubiquitously expressed across tissues, assuring the broad applicability of LEAPER. The ADAR1^p150^ is an interferon-inducible isoform^58^, and has proven to be functional in LEAPER (Fig. 1e, Extended Data Fig. 2b). Thus, co-transfection of interferon stimulatory RNAs with the arRNA might further improve editing efficiency under certain circumstances. Alternatively, as ADAR3 plays inhibitory roles, inhibition of ADAR3 might enhance editing efficiency in ADAR3-expressing cells. Moreover, additional modification of arRNA might increase its editing efficiency. For instance, arRNA fused with certain ADAR-recruiting scaffold may increase local ADAR protein concentration and consequently boost editing yield. So far, we could only leverage endogenous ADAR1/2 proteins for the A to I base conversion. It is exciting to explore whether more native mechanisms could be harnessed similarly for the modification of genetic elements, especially to realize potent nucleic acid editing.

Altogether, we provided a proof of principle that the endogenous machinery in cells could be co-opted to edit RNA transcripts. We demonstrated that LEAPER is a simple, efficient and safe system, shedding light on a novel path for gene editing-based therapeutics and research.

## Supporting information

Supplementary Figures and Sequences

## Acknowledgements

We acknowledge the staff of the BIOPIC High-throughput Sequencing Center (Peking University) and Genetron Health for their assistance in NGS analysis, the National Center for Protein Sciences (Beijing) and the core facilities at the School of Life Sciences (Peking University, X. Zhang, F. Wang and L. Du) for help with Fluorescence Activated Cell Sorting. We thank High-performance Computing Platform of Peking University for providing platforms of NGS data analysis. We thank M. Mo for technical assistance. We thank J. Wang for providing plasmids encoding disease-relevant genes and primary cells. We also thank Z. Jiang for providing the mouse melanoma cell line B16. This project was supported by funds from Beijing Municipal Science & Technology Commission (Z181100001318009), the National Science Foundation of China (31430025), Beijing Advanced Innovation Center for Genomics at Peking University, and the Peking-Tsinghua Center for Life Sciences (to W.W.); the National Science Foundation of China (31870893) and the National Major Science & Technology Project for Control and Prevention of Major Infectious Diseases in China (2018ZX10301401, to Z.Z.); and Beijing Nova Program (Z181100006218042, to P.Y.).

## Author Contributions

W.W. conceived and supervised the project. W.W., L.Q., Z.Y., S.Z., C.W., Z.C. and Z.Z. designed the experiments. L.Q., Z.Y., C.W., S.Z., Z.C. and P.Y. performed the experiments with the help from F.T., Y.B. and Y.Z.. Y.Y. conducted all the sample preparation for NGS. Z.Y. and Z.L. performed the data analysis. L.Q., S.Z., Z.Z. and W.W. wrote the manuscript with the help of all other authors.

## Methods

### Plasmids construction

For the three versions of dual fluorescence reporters (Reporter-1, -2 and -3), *mCherry* and *EGFP* (the start codon ATG of EGFP was deleted) coding sequences were PCR amplified, digested using BsmBI (Thermo Fisher Scientific, ER0452), followed by T4 DNA ligase (NEB, M0202L)-mediated ligation with GGGGS linkers. The ligation product was subsequently inserted into the pLenti-CMV-MCS-PURO backbone.

For the dLbuCas13-ADAR _DD_ (E1008Q) expressing construct, the ADAR1_DD_ gene was amplified from the ADAR1^p150^ construct (a gift from Jiahuai Han’s lab, Xiamen University). The dLbuCas13 gene was amplified by PCR from the Lbu_C2c2_R472A_H477A_R1048A_ H1053A plasmid (Addgene #83485). The ADAR1_DD_ (hyperactive E1008Q variant) was generated by overlap-PCR and then fused to dLbuCas13. The ligation products were inserted into the pLenti-CMV-MCS-BSD backbone.

For arRNA-expressing construct, the sequences of arRNAs were synthesized and golden-gate cloned into the pLenti-sgRNA-lib 2.0 (Addgene #89638) backbone, and the transcription of arRNA was driven by hU6 promoter. For the ADAR expressing constructs, the full length ADAR1^p110^ and ADAR1^p150^ were PCR amplified from the ADAR1^p150^ construct, and the full length ADAR2 were PCR amplified from the ADAR2 construct (a gift from Jiahuai Han’s lab, Xiamen University). The amplified products were then cloned into the pLenti-CMV-MCS-BSD backbone.

For the constructs expressing genes with pathogenic mutations, full length coding sequences of *TP53* (ordered from Vigenebio) and other 6 disease-relevant genes (*COL3A1*, *BMPR2*, *AHI1*, *FANCC*, *MYBPC3* and *IL2RG*, gifts from Jianwei Wang’s lab, Institute of pathogen biology, Chinese Academy of Medical Sciences) were amplified from the constructs encoding the corresponding genes with introduction of G>A mutations through mutagenesis PCR. The amplified products were cloned into the pLenti-CMV-MCS-mCherry backbone through Gibson cloning method^59^.

### Cell culture

The HeLa and B16 cell lines were from Z. Jiang’s laboratory (Peking University). And the HEK293T cell line was from C. Zhang’s laboratory (Peking University). RD cell line was from J Wang’s laboratory (Institute of Pathogen Biology, Peking Union Medical College & Chinese Academy of Medical Sciences). SF268 cell lines were from Cell Center, Institute of Basic Medical Sciences, Chinese Academy of Medical Sciences. A549 and SW13 cell lines were from EdiGene Inc. HepG2, HT29, NIH3T3, and MEF cell lines were maintained in our laboratory at Peking University. These mammalian cell lines were cultured in Dulbecco’s Modified Eagle Medium (Corning, 10-013-CV) with 10% fetal bovine serum (CellMax, SA201.02), additionally supplemented with 1% penicillin–streptomycin under 5% CO_2_ at 37°C. Unless otherwise described, cells were transfected with the X-tremeGENE HP DNA transfection reagent (Roche, 06366546001) according to the manufacturer’s instruction.

The human primary pulmonary fibroblasts (#3300) and human primary bronchial epithelial cells (#3210) were purchased from ScienCell Research Laboratories, Inc. and were cultured in Fibroblast Medium (ScienCell, #2301) and Bronchial Epithelial Cell Medium (ScienCell, #3211), respectively. Both media were supplemented with 15% fetal bovine serum (BI) and 1% penicillin–streptomycin. The primary GM06214 and GM01323 cells were ordered from Coriell Institute for Medical Research and cultured in Dulbecco’s Modified Eagle Medium (Corning, 10-013-CV) with 15% fetal bovine serum (BI) and 1% penicillin–streptomycin. All cells were cultured under 5% CO_2_at 37°C.

### Isolation and culture of human primary T cells

Primary human T cells were isolated from leukapheresis products from healthy human donor. Briefly, Peripheral blood mononuclear cells (PBMCs) were isolated by Ficoll centrifugation (Dakewei, AS1114546), and T cells were isolated by magnetic negative selection using an EasySep Human T Cell Isolation Kit (STEMCELL, 17951) from PBMCs. After isolation, T cells were cultured in X-vivo15 medium, 10% FBS and IL2 (1000 U/ml) and stimulated with CD3/CD28 DynaBeads (ThermoFisher, 11131D) for 2 days. Leukapheresis products from healthy donors were acquired from AllCells LLC China. All healthy donors provided informed consent.

### Cell line construction

For the stable reporter cell lines, the reporter constructs (pLenti-CMV-MCS-PURO backbone) were co-transfected into HEK293T cells, together with two viral packaging plasmids, pR8.74 and pVSVG. 72 hours later, the supernatant virus was collected and stored at −80°C. The HEK293T cells were infected with lentivirus, then mCherry-positive cells were sorted via FACS and cultured to select a single clone cell lines stably expressing dual fluorescence reporter system without detectable EGFP background. The HEK293T *ADAR1*^−/−^ and *TP53*^−/−^ cell lines were generated according to a previously reported method^60^. ADAR1-targeting sgRNA and PCR amplified donor DNA containing CMV-driven puromycin resistant gene were co-transfected into HEK293T cells. Then cells were treated with puromycin 7 days after transfection. Single clones were isolated from puromycin resistant cells followed by verification through sequencing and Western blot.

### RNA editing of endogenous or exogenous-expressed transcripts

For assessing RNA editing on the dual fluorescence reporter, HEK293T cells or HEK293T *ADAR1*^−/−^ cells were seeded in 6-well plates (6×10 ^5^ cells/well). 24 hours later, cells were co-transfected with 1.5 μg reporter plasmids and 1.5 μg arRNA plasmids. To examine the effect of ADAR1^p110^, ADAR1^p150^ or ADAR2 protein expression, the editing efficiency was assayed by EGFP positive ratio and deep sequencing.

HEK293T *ADAR1*^−/−^ cells were seeded in 12-well plates (2.5×10 ^5^ cells/well). 24 hours later, cells were co-transfected with 0.5 μg of reporter plasmids, 0.5 μg arRNA plasmids and 0.5 μg ADAR1/2 plasmids (pLenti backbone as control). The editing efficiency was assayed by EGFP positive ratio and deep sequencing.

To assess RNA editing on endogenous mRNA transcripts, HEK293T cells were seeded in 6-well plates (6×10 ^5^ cells/well). Twenty-four hours later, cells were transfected with 3 μg of arRNA plasmids. The editing efficiency was assayed by deep sequencing.

To assess RNA editing efficiency in multiple cell lines, 8-9×10 ^4^ (RD, SF268, HeLa) or 1.5×10 ^5^ (HEK293T) cells were seeded in 12-well plates. For cells difficult to transfect, such as HT29, A549, HepG2, SW13, NIH3T3, MEF and B16, 2-2.5×10 ^5^ cells were seeded in 6-well plate. Twenty-four hours later, reporters and arRNAs plasmid were co-transfected into these cells. The editing efficiency was assayed by EGFP positive ratio.

To evaluate EGFP positive ratio, at 48 to 72 hrs post transfection, cells were sorted and collected by Fluorescence-activated cell sorting (FACS) analysis. The mCherry signal was served as a fluorescent selection marker for the reporter/arRNA-expressing cells, and the percentages of EGFP^+^/mCherry^+^ cells were calculated as the readout for editing efficiency. For NGS quantification of the A to I editing rate, at 48 to 72 hr post transfection, cells were sorted and collected by FACS assay and were then subjected to RNA isolation (TIANGEN, DP420). Then, the total RNAs were reverse-transcribed into cDNA via RT-PCR (TIANGEN, KR103-04), and the targeted locus was PCR amplified with the corresponding primers (listed in the Supplementary Table 4). The PCR products were purified for Sanger sequencing or NGS (Illumina HiSeq X Ten).

### RNA editing analysis for targeted sites

For deep sequencing analysis, an index was generated using the targeted site sequence (upstream and downstream 20-nt) of arRNA covering sequences. Reads were aligned and quantified using BWA version 0.7.10-r789. Alignment BAMs were then sorted by Samtools, and RNA editing sites were analyzed using REDitools version 1.0.4. The parameters are as follows: -U [AG or TC] -t 8 -n 0.0 -T 6-6 -e -d -u. All the significant A>G conversion within arRNA targeting region calculated by Fisher’s exact test (*p*-value < 0.05) were considered as edits by arRNA. The conversions except for targeted adenosine were off-target edits. The mutations that appeared in control and experimental groups simultaneously were considered as SNP.

### Transcriptome-wide RNA-sequencing analysis

The Ctrl RNA_151_ or arRNA_151_-PPIB-expressing plasmids with BFP expression cassette were transfected into HEK293T cells. The BFP^+^ cells were enriched by FACS 48 hours after transfection, and RNAs were purified with RNAprep Pure Micro kit (TIANGEN, DP420). The mRNA was then purified using NEBNext Poly(A) mRNA Magnetic Isolation Module (New England Biolabs, E7490), processed with the NEBNext Ultra II RNA Library Prep Kit for Illumina (New England Biolabs, E7770), followed by deep sequencing analysis using Illumina HiSeq X Ten platform (2× 150-bp paired end; 30G for each sample). To exclude nonspecific effect caused by transfection, we included the mock group in which we only treated cells with transfection reagent. Each group contained four replications.

The bioinformatics analysis pipeline was referred to the work by Vogel *et al*^22^. The quality control of analysis was conducted by using FastQC, and quality trim was based on Cutadapt (the first 6-bp for each reads were trimmed and up to 20-bp were quality trimmed). AWK scripts were used to filtered out the introduced arRNAs. After trimming, reads with lengths less than 90-nt were filtered out. Subsequently, the filtered reads were mapped to the reference genome (GRCh38-hg38) by STAR software^61^. We used the GATK Haplotypcaller^62^ to call the variants. The raw VCF files generated by GATK were filtered and annotated by GATK VariantFiltration, bcftools and ANNOVAR^63^. The variants in dbSNP, 1000 Genome^64^, EVS were filtered out. The shared variants in four replicates of each group were then selected as the RNA editing sites. The RNA editing level of Mock group was viewed as the background, and the global targets of Ctrl RNA_151_ and arRNA_151_-PPIB were obtained by subtracting the variants in the Mock group.

To assess if LEAPER perturbs natural editing homeostasis, we analyzed the global editing sites shared by Mock group and arRNA_151_-PPIB group (or Ctrl RNA_151_ group). The differential RNA editing rates at native A-to-I editing sites were assessed with Pearson’s correlation coefficient analysis. Pearson correlations of editing rate between Mock group and arRNA_151_-PPIB group (or Ctrl RNA_151_ group) were calculated and annotated in Fig. 6.

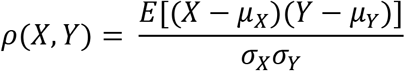

*X* means the editing rate of each site in the Mock group; *Y* means the editing rate of each site in the Ctrl RNA_151_ group (Fig. 6a) or arRNA_151_-PPIB group (Fig. 6b); *σx* is the standard deviation of *X; σY* is the standard deviation of *Y; μX* is the mean of *X; μY* is the mean of *Y; E* is the expectation.

The RNA-Seq data were analysed for the interrogation of possible transcriptional changes induced by RNA editing events. The analysis of transcriptome-wide gene expression was performed using HISAT2 and STRINGTIE software^65^. We used Cutadapt and FastQC for the quality control of the sequencing data. The sequencing reads were then mapped to reference genome (GRCh38-hg38) using HISAT2, followed by Pearson’s correlation coefficient analysis as mentioned above.

### Western blot

We used the mouse monoclonal primary antibodies respectively against ADAR1 (Santa Cruz, sc-271854), ADAR2 (Santa Cruz, sc-390995), ADAR3 (Santa Cruz, sc-73410), p53 (Santa Cruz, sc-99), KRAS (Sigma, SAB1404011); GAPDH (Santa Cruz, sc-47724) and β-tubulin (CWBiotech, CW0098). The HRP-conjugated goat anti-mouse IgG (H+L, 115-035-003) secondary antibody was purchased from Jackson ImmunoResearch. 2×10 ^6^ cells were sorted to be lysed and an equal amount of each lysate was loaded for SDS-PAGE. Then, sample proteins were transferred onto PVDF membrane (Bio-Rad Laboratories) and immunoblotted with primary antibodies against one of the ADAR enzymes (anti-ADAR1, 1:500; anti-ADAR2, 1:100; anti-ADAR3, 1:800), followed by secondary antibody incubation (1:10,000) and exposure. The β-Tubulin were re-probed on the same PVDF membrane after stripping of the ADAR proteins with the stripping buffer (CWBiotech, CW0056). The experiments were repeated three times. The semi-quantitative analysis was done with Image Lab software.

### Cytokine expression assay

HEK293T cells were seeded on 12 wells plates (2×10 ^5^ cells/well). When approximately 70% confluent, cells were transfected with 1.5 μg of arRNA. As a positive control, 1 μg of poly(I:C) (Invitrogen, tlrl-picw) was transfected. Forty-eight hours later, cells were collected and subjected to RNA isolation (TIANGEN, DP430). Then, the total RNAs were reverse-transcribed into cDNA via RT-PCR (TIANGEN, KR103-04), and the expression of IFN-β and IL-6 were measured by quantitative PCR (TAKARA, RR820A). The sequences of the primers were listed in the Supplementary Table 4.

### Transcriptional regulatory activity assay of p53

The *TP53*^W53X^ cDNA-expressing plasmids and arRNA-expressing plasmids were co-transfected into HEK293T *TP53*^−/−^ cells, together with p53-Firefly-luciferase *cis*-reporting plasmids (YRGene, VXS0446) and Renilla-luciferase plasmids (a gift from Z. Jiang’s laboratory, Peking University) for detecting the transcriptional regulatory activity of p53. 48 hrs later, the cells were harvested and assayed with the Promega Dual-Glo Luciferase Assay System (Promega, E4030) according to the manufacturer protocol. Briefly, 150 μL Dual-Glo Luciferase Reagent was added to the harvested cell pellet, and 30 minutes later, the Firefly luminescence was measured by adding 100 μL Dual-Glo Luciferase Reagent (cell lysis) to 96-well white plate by Infinite M200 reader (TECAN). 30 min later, 100 μL Dual-Glo stop and Glo Reagent were sequentially added to each well to measure the Renilla luminescence and calculate the ratio of Firefly luminescence to Renilla luminescence.

### Electroporation in primary cells

For arRNA-expressing plasmids electroporation in the human primary pulmonary fibroblasts or human primary bronchial epithelial cells, 20 μg plasmids were electroporated with Nucleofector^TM^ 2b Device (Lonza) and Basic Nucleofector^TM^ Kit (Lonza, VPI-1002), and the electroporation program was U-023. For arRNA-expressing plasmids electroporation in human primary T cells, 20 μg plasmids were electroporated into human primary T with Nucleofector^TM^ 2b Device (Lonza) and Human T cell Nucleofector^TM^ Kit (Lonza, VPA-1002), and the electroporation program was T-024. Forty-eight hours post-electroporation, cells were sorted and collected by FACS assay and were then subjected to the following deep-sequencing for targeted RNA editing assay. The electroporation efficiency was normalized according to the fluorescence marker.

For the chemosynthetic arRNA or control RNA electroporation in human primary T cells or primary GM06214 cells, RNA oligo was dissolved in 100 μL opti-MEM medium (Gbico, 31985070) with the final concentration 2 μM. Then 1×10E6 GM06214 cells or 3×10E6 T cells were resuspended with the above electroporation mixture and electroporated with Agile Pulse In Vivo device (BTX) at 450 V for 1 ms. Then the cells were transferred to warm culture medium for the following assays.

### α-L-iduronidase (IDUA) catalytic activity assay

The harvested cell pellet was resuspended and lysed with 28 μL 0.5% Triton X-100 in 1×PBS buffer on ice for 30 minutes. And then 25 μL of the cell lysis was added to 25 μL 190 μM 4-methylumbelliferyl-α-L-iduronidase substrate (Cayman, 2A-19543-500), which was dissolved in 0.4 M sodium formate buffer containing 0.2% Triton X-100, pH 3.5, and incubated for 90 minutes at 37°C in the dark. The catalytic reaction was quenched by adding 200 μL 0.5M NaOH/Glycine buffer, pH 10.3, and then centrifuged for 2 minutes at 4°C. The supernatant was transferred to a 96-well plate, and fluorescence was measured at 365 nm excitation wavelength and 450 nm emission wavelength with Infinite M200 reader (TECAN).

