## Supplementary Figures and Sequences for "Leveraging Endogenous ADAR for Programmable Editing on RNA"

#### Extended Data Figure Legends

**Extended Data Figure 1 | Exploration of an efficient RNA editing platform. a**, Schematic of dLbuCas13a-ADAR1$_{DD}$ (E1008Q) fusion protein and the corresponding crRNA. The catalytic inactive LbuCas13a was fused to the deaminase domain of ADAR1 (hyperactive E1008Q variant) using 3× GGGGS linker. The crRNA (crRNA$^{Cas13a}$) consisted of Lbu-crRNA scaffold and a spacer which was complementary to the targeting RNA with an A-C mismatch as indicated. **b**, Schematic of dual fluorescent reporter system and the Lbu-crRNA with various lengths of spacers as indicated. **c**, Quantification of the EGFP positive (EGFP$^{+}$) cells. HEK293T cells stably expressing the Repoter-1 were transfected with indicated lengths of crRNA$^{Cas13a}$, with or without co-expression of the dLbuCas13a-ADAR1$_{DD}$ (E1008Q), followed by FACS analysis. Data are presented as the mean ± s.e.m. (n = 3). **d**, Representative FACS result from the experiment performed with the control (Ctrl crRNA$_{70}$) or the targeting spacer (crRNA$_{70}$).

**Extended Data Figure 2 | mRNA expression level of ADAR1/ADAR2 and arRNA-mediated RNA editing. a**, Quantitative PCR showing the mRNA levels of *ADAR1* and *ADAR2* in HEK293T cells. Data are presented as the mean ± s.e.m. (n = 3). **b**, Representative FACS results from Fig. 1e.

**Extended Data Figure 3 | Quantitative PCR showing the effects of LEAPER on the expression levels of targeted Reporter-1 transcripts by 111-nt arRNA or control RNA in HEK293T cells.** Data are presented as the mean ± s.e.m. (n = 3); unpaired two-sided Student's *t*-test, ns, not significant.

**Extended Data Figure 4 | Schematic of Reporter-1 (a), -2 (b), and -3 (c), as well as their corresponding arRNAs.**

**Extended Data Figure 5 | Effects of LEAPER on the expression levels of targeted transcripts and protein products. a**, Quantitative PCR showing the expression levels of targeted transcripts from PPIB, KRAS, SMAD4 and FANCC by the corresponding 151-nt arRNA or Control RNA in HEK293T cells. Data are presented as the mean ± s.e.m. (n = 3); unpaired two-sided Student's *t*-test, **P* < 0.05; ***P* < 0.01; ****P* < 0.001; *****P* < 0.0001; ns, not significant. **b**, Western blot results showing the effects on protein products of targeted KRAS gene by 151-nt arRNA in HEK293T cells. β-tubulin was used as a loading control.

**Extended Data Figure 6 | Editing endogenous transcripts with LEAPER. a**, Schematic of the *KARS* transcript sequence covered by the 151-nt arRNA. The arrow indicates the targeting adenosine. All adenosines were marked in red. **b**, Heatmap of editing rate on adenosines covered by indicated arRNAs in the *KARS* transcript (marked in the bold frame in blue). **c**, Schematic of the *SMAD4* transcript covered by the 151-nt arRNA. **d**, Heatmap of editing rate on adenosines covered by indicated arRNAs in the *SMAD4* transcript. **e**, Schematic of the *FANCC* transcript covered by the 151-nt arRNA. **f**, Heatmap of editing rate on adenosines covered by indicated arRNAs in the *FANCC* transcript. For each arRNA, the region of duplex RNA is highlighted with bold frame in blue. Data (**b**, **d**, and **f**) are presented as the mean (n = 3).

**Extended Data Figure 7 | Evaluation of potential off-targets. a**, Schematic of the highly complementary region of $arRNA_{111}$-FANCC and the indicated potential off-target sequence, which were predicted by searching homologous sequences through NCBI-BLAST. **b**, Deep sequencing showing the editing rate on the on-target site and all predicted off-target sites of $arRNA_{111}$-FANCC. All data are presented as the mean ± s.e.m. (n = 3).

**Extended Data Figure 8 | Editing mutant *TP53*$^{W53X}$ transcripts by LEAPER.** Top, schematic of the *TP53* transcript sequence covered by the 111-nt arRNAs. The arrow indicates the targeted adenosine. All adenosines were marked in red. Bottom, a heatmap of editing rate on adenosines covered by indicated arRNAs in the *TP53* transcript.

**Extended Data Figure 9 | Schematic representation of the selected disease-relevant cDNA containing G to A mutation from ClinVar data and the corresponding 111-nt arRNA.**

### Supplementary Tables

**Supplementary Table 1 | LbuCas13 crRNA sequences.**

**Supplementary Table 2 | Sequences of arRNAs and control RNAs used in this study.**

**Supplementary Table 3 | Disease-relevant cDNAs used in this study.**

**Supplementary Table 4 | Primers used in this study.**

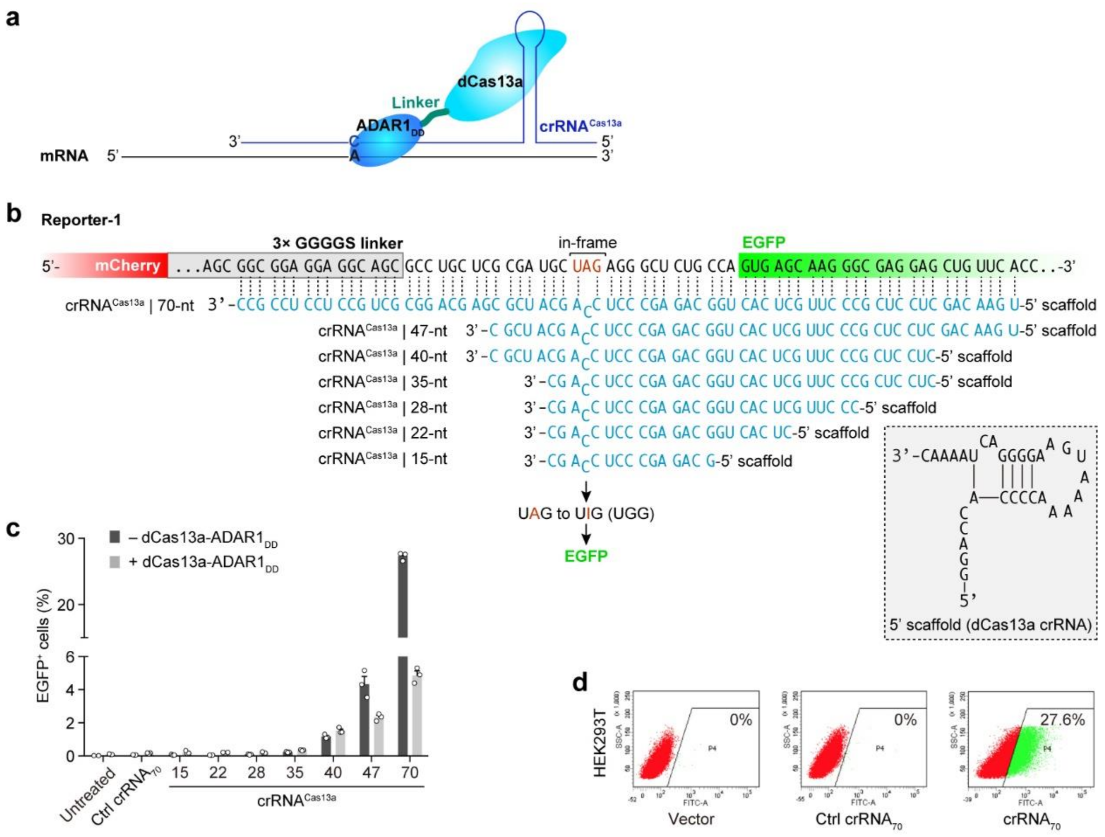

Extended Data Figure 1

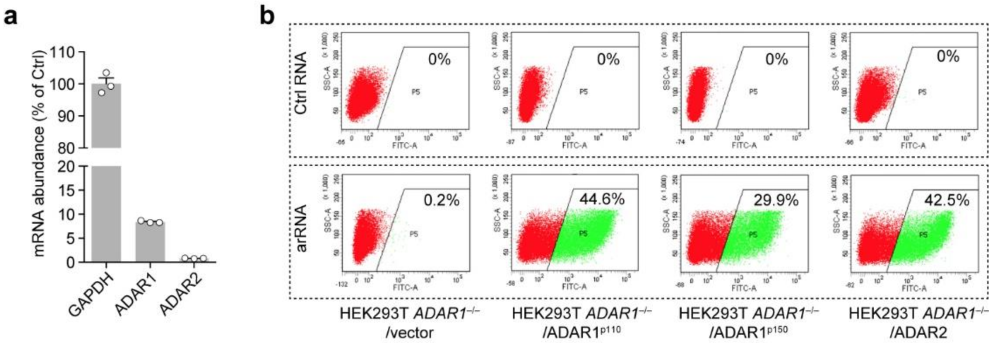


Extended Data Figure 2

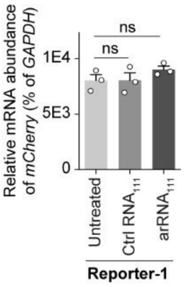


Extended Data Figure 3

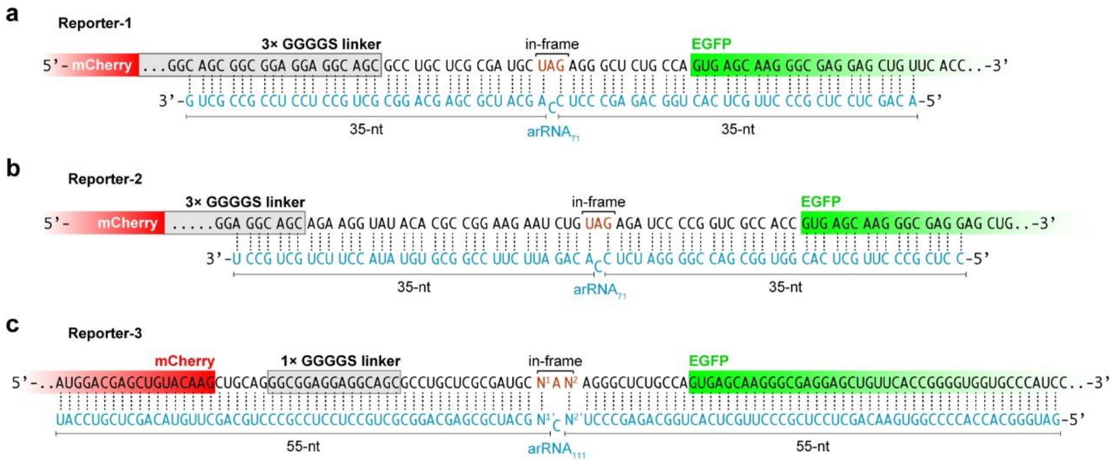


Extended Data Figure 4

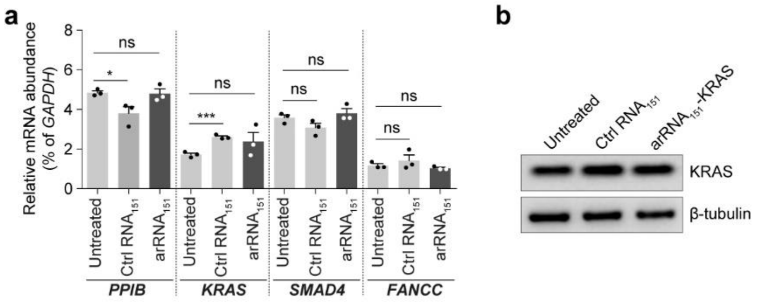


Extended Data Figure 5

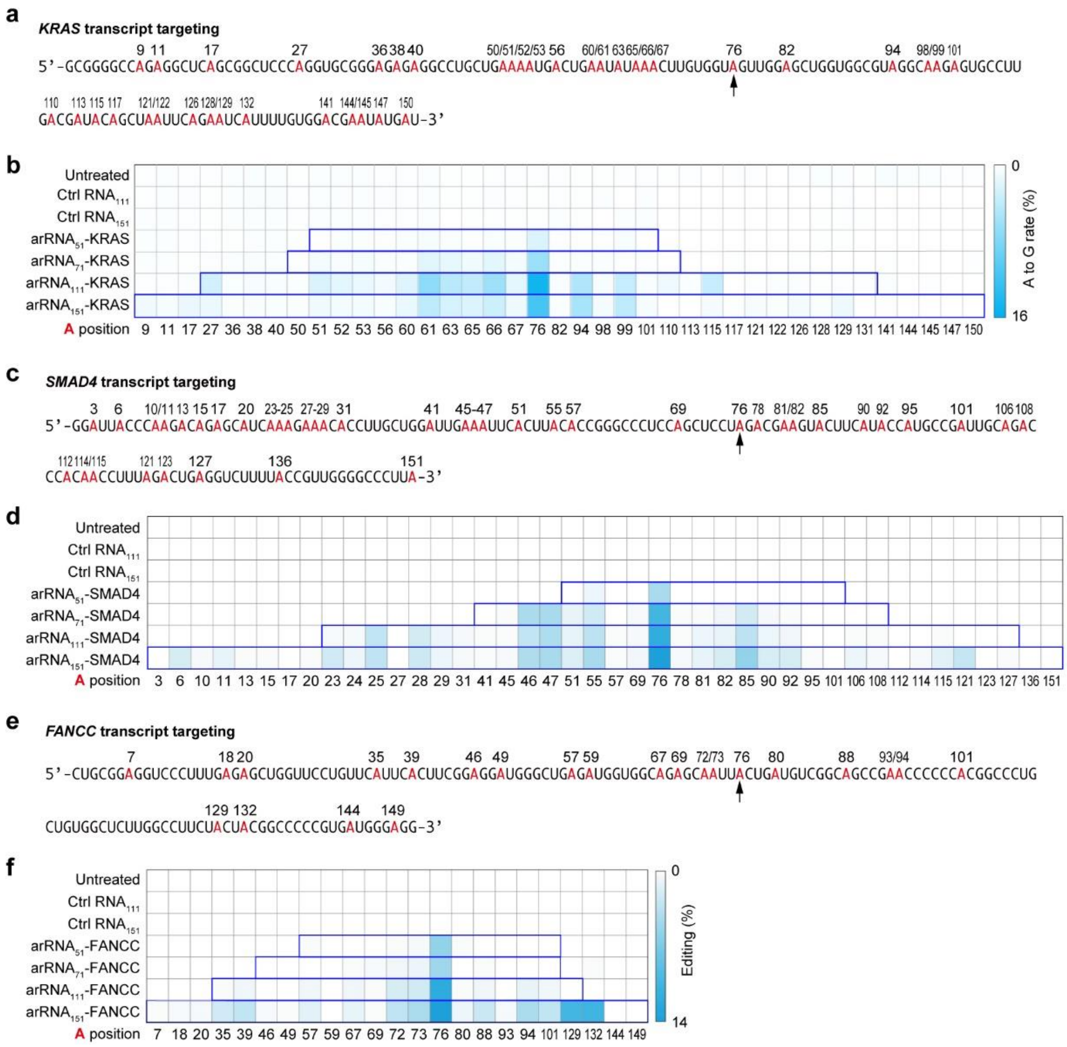


Extended Data Figure 6

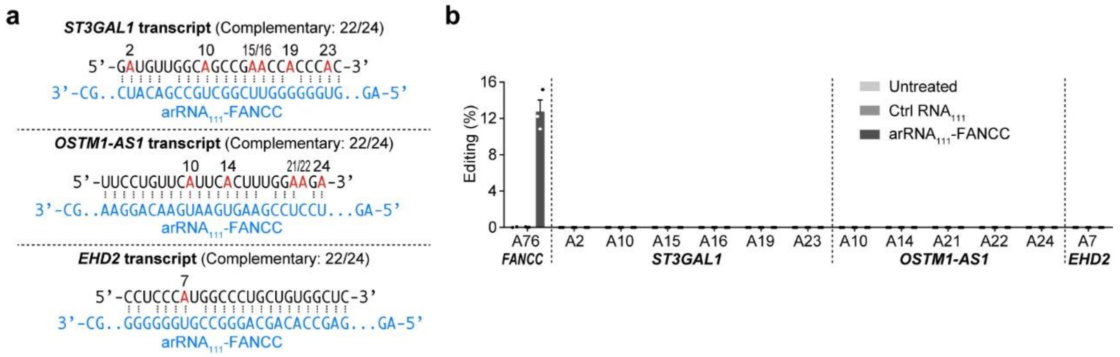


Extended Data Figure 7

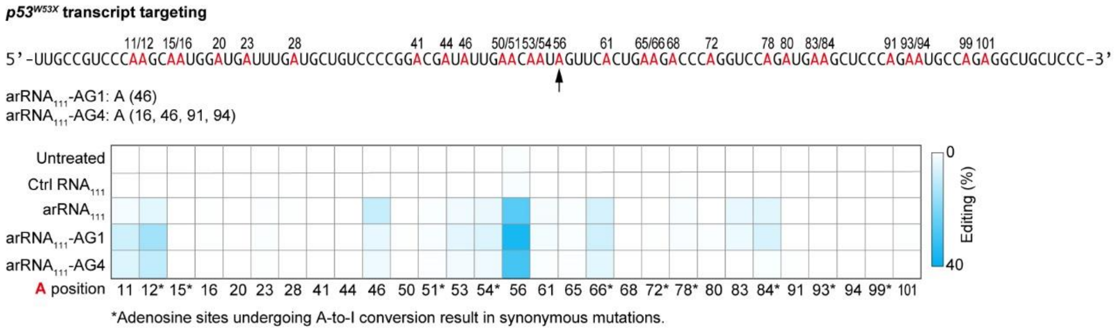


Extended Data Figure 8

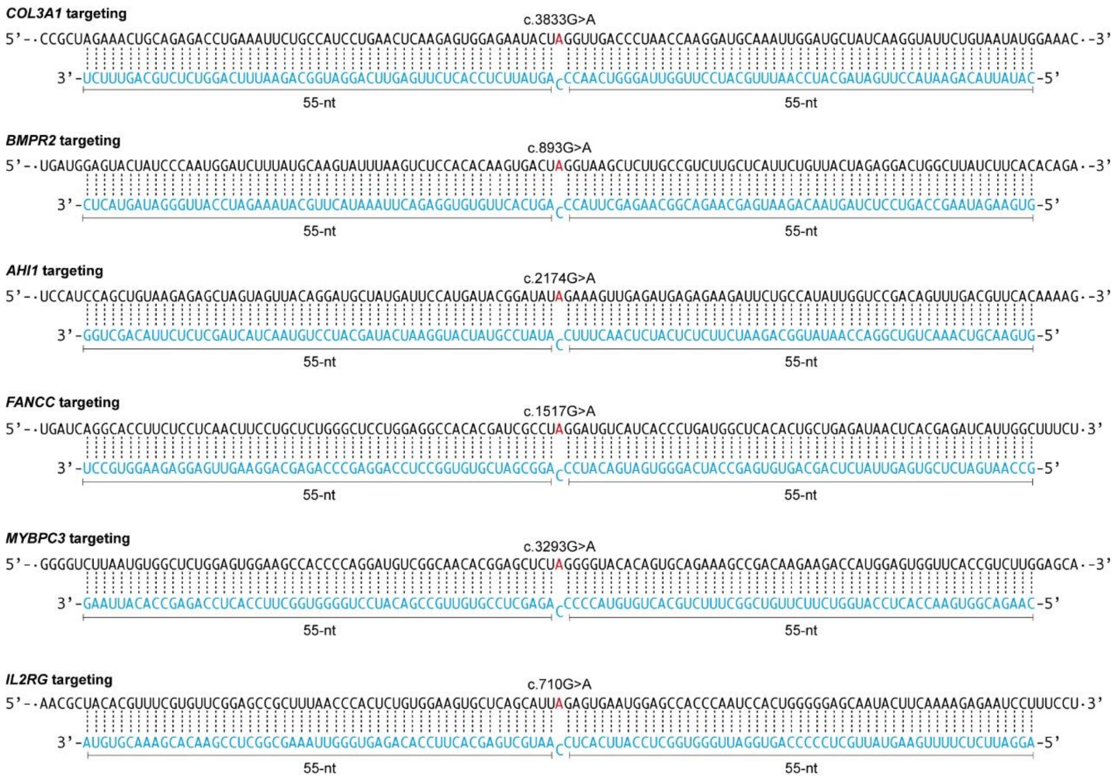


Extended Data Figure 9

**Supplementary sequences**

**Reporter-1:**

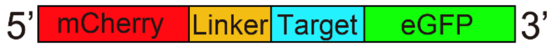


5'-
ATGGTGAGCAAGGGCGAGGAGGATAACATGGCCATCATCAAGGAGTTCAT
GCGCTTCAAGGTGCACATGGAGGGCTCCGTGAACGGCCACGAGTTCGAGA
TCGAGGGCGAGGGCGAGGGCCGCCCCTACGAGGGCACCCAGACCGCCAA
GCTGAAGGTGACCAAGGGTGGCCCCCTGCCCTTCGCCTGGGACATCCTGT
CCCCTCAGTTCATGTACGGCTCCAAGGCCTACGTGAAGCACCCCGCCGACA
TCCCCGACTACTTGAAGCTGTCCTTCCCCGAGGGCTTCAAGTGGGAGCGCG
TGATGAACTTCGAGGACGGCGGCGTGGTGACCGTGACCCAGGACTCCTCC
CTGCAGGACGGCGAGTTCATCTACAAGGTGAAGCTGCGCGGCACCAACTT
CCCCTCCGACGGCCCCGTAATGCAGAAGAAGACCATGGGCTGGGAGGCCT
CCTCCGAGCGGATGTACCCCGAGGACGGCGCCCTGAAGGGCGAGATCAAG
CAGAGGCTGAAGCTGAAGGACGGCGGCCACTACGACGCTGAGGTCAAGA
CCACCTACAAGGCCAAGAAGCCCGTGCAGCTGCCCGGCGCCTACAACGTC
AACATCAAGTTGGACATCACCTCCCACAACGAGGACTACACCATCGTGGA
ACAGTACGAACGCGCCGAGGGCCGCCACTCCACCGGCGGCATGGACGAGC
TGTACAAG CTGCAG GGCGGAGGAGGCAGC GGCGGAGGAGGCAGC
GGCGGAGGAGGCAGC GCCTGCTCGCGATGCTAGAGGGCTCTGCCA
GTGAGCAAGGGCGAGGAGCTGTTCACCGGGGTGGTGCCCATCCTGGTCGA
GCTGGACGGCGACGTAAACGGCCACAAGTTCAGCGTGTCCGGCGAGGGCG
AGGGCGATGCCACCTACGGCAAGCTGACCCTGAAGTTCATCTGCACCACC
GGCAAGCTGCCCGTGCCCTGGCCCACCCTCGTGACCACCCTGACCTACGG
CGTGCAGTGCTTCAGCCGCTACCCCGACCACATGAAGCAGCACGACTTCTT
CAAGTCCGCCATGCCCGAAGGCTACGTCCAGGAGCGCACCATCTTCTTCAA
GGACGACGGCAACTACAAGACCCGCGCCGAGGTGAAGTTCGAGGGCGAC
ACCCTGGTGAACCGCATCGAGCTGAAGGGCATCGACTTCAAGGAGGACGG
CAACATCCTGGGGCACAAGCTGGAGTACAACTACAACAGCCACAACGTCT
ATATCATGGCCGACAAGCAGAAGAACGGCATCAAGGTGAACTTCAAGATC
CGCCACAACATCGAGGACGGCAGCGTGCAGCTCGCCGACCACTACCAGCA
GAACACCCCCATCGGCGACGGCCCCGTGCTGCTGCCCGACAACCACTACC
TGAGCACCCAGTCCGCCCTGAGCAAAGACCCCAACGAGAAGCGCGATCAC
ATGGTCCTGCTGGAGTTCGTGACCGCCGCCGGGATCACTCTCGGCATGGAC
GAGCTGTACAAGTAA-3'

**Reporter-2:**

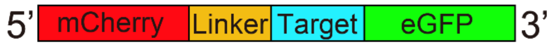


5'-
ATGGTGAGCAAGGGCGAGGAGGATAACATGGCCATCATCAAGGAGTTCAT
GCGCTTCAAGGTGCACATGGAGGGCTCCGTGAACGGCCACGAGTTCGAGA
TCGAGGGCGAGGGCGAGGGCCGCCCCTACGAGGGCACCCAGACCGCCAA

GCTGAAGGTGACCAAGGGTGGCCCCCTGCCCTTCGCCTGGGACATCCTGT
CCCCTCAGTTCATGTACGGCTCCAAGGCCTACGTGAAGCACCCCGCCGACA
TCCCCGACTACTTGAAGCTGTCCTTCCCCGAGGGCTTCAAGTGGGAGCGCG
TGATGAACTTCGAGGACGGCGGCGTGGTGACCGTGACCCAGGACTCCTCC
CTGCAGGACGGCGAGTTCATCTACAAGGTGAAGCTGCGCGGCACCAACTT
CCCCTCCGACGGCCCCGTAATGCAGAAGAAGACCATGGGCTGGGAGGCCT
CCTCCGAGCGGATGTACCCCGAGGACGGCGCCCTGAAGGGCGAGATCAAG
CAGAGGCTGAAGCTGAAGGACGGCGGCCACTACGACGCTGAGGTCAAGA
CCACCTACAAGGCCAAGAAGCCCGTGCAGCTGCCCGGCGCCTACAACGTC
AACATCAAGTTGGACATCACCTCCCACAACGAGGACTACACCATCGTGGA
ACAGTACGAACGCGCCGAGGGCCGCCACTCCACCGGCGGCATGGACGAGC
TGTACAAG CTGCAG GGCGGAGGAGGCAGC GGCGGAGGAGGCAGC
GGCGGAGGAGGCAGC
AGAAGGTATACACGCCGGAAGAATCTG<u>TAG</u>AGATCCCCGGTCGCCACC
GTGAGCAAGGGCGAGGAGCTGTTCACCGGGGTGGTGCCCATCCTGGTCGA
GCTGGACGGCGACGTAAACGGCCACAAGTTCAGCGTGTCCGGCGAGGGCG
AGGGCGATGCCACCTACGGCAAGCTGACCCTGAAGTTCATCTGCACCACC
GGCAAGCTGCCCGTGCCCTGGCCCACCCTCGTGACCACCCTGACCTACGG
CGTGCAGTGCTTCAGCCGCTACCCCGACCACATGAAGCAGCACGACTTCTT
CAAGTCCGCCATGCCCGAAGGCTACGTCCAGGAGCGCACCATCTTCTTCAA
GGACGACGGCAACTACAAGACCCGCGCCGAGGTGAAGTTCGAGGGCGAC
ACCCTGGTGAACCGCATCGAGCTGAAGGGCATCGACTTCAAGGAGGACGG
CAACATCCTGGGGCACAAGCTGGAGTACAACTACAACAGCCACAACGTCT
ATATCATGGCCGACAAGCAGAAGAACGGCATCAAGGTGAACTTCAAGATC
CGCCACAACATCGAGGACGGCAGCGTGCAGCTCGCCGACCACTACCAGCA
GAACACCCCCATCGGCGACGGCCCCGTGCTGCTGCCCGACAACCACTACC
TGAGCACCCAGTCCGCCCTGAGCAAAGACCCCAACGAGAAGCGCGATCAC
ATGGTCCTGCTGGAGTTCGTGACCGCCGCCGGGATCACTCTCGGCATGGAC
GAGCTGTACAAGTAA-3’

**Reporter-3:**

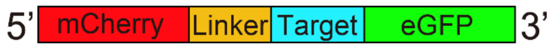


5’-
ATGGTGAGCAAGGGCGAGGAGGATAACATGGCCATCATCAAGGAGTTCAT
GCGCTTCAAGGTGCACATGGAGGGCTCCGTGAACGGCCACGAGTTCGAGA
TCGAGGGCGAGGGCGAGGGCCGCCCCTACGAGGGCACCCAGACCGCCAA
GCTGAAGGTGACCAAGGGTGGCCCCCTGCCCTTCGCCTGGGACATCCTGT
CCCCTCAGTTCATGTACGGCTCCAAGGCCTACGTGAAGCACCCCGCCGACA
TCCCCGACTACTTGAAGCTGTCCTTCCCCGAGGGCTTCAAGTGGGAGCGCG
TGATGAACTTCGAGGACGGCGGCGTGGTGACCGTGACCCAGGACTCCTCC
CTGCAGGACGGCGAGTTCATCTACAAGGTGAAGCTGCGCGGCACCAACTT
CCCCTCCGACGGCCCCGTAATGCAGAAGAAGACCATGGGCTGGGAGGCCT
CCTCCGAGCGGATGTACCCCGAGGACGGCGCCCTGAAGGGCGAGATCAAG

CAGAGGCTGAAGCTGAAGGACGGCGGCCACTACGACGCTGAGGTCAAGA
CCACCTACAAGGCCAAGAAGCCCGTGCAGCTGCCCGGCGCCTACAACGTC
AACATCAAGTTGGACATCACCTCCCACAACGAGGACTACACCATCGTGGA
ACAGTACGAACGCGCCGAGGGCCGCCACTCCACCGGCGGCATGGACGAGC
TGTACAAG CTGCAG GGCGGAGGAGGCAGC
GCCTGCTCGCGATGCTAGAGGGCTCTGCCA
GTGAGCAAGGGCGAGGAGCTGTTCACCGGGGTGGTGCCCATCCTGGTCGA
GCTGGACGGCGACGTAAACGGCCACAAGTTCAGCGTGTCCGGCGAGGGCG
AGGGCGATGCCACCTACGGCAAGCTGACCCTGAAGTTCATCTGCACCACC
GGCAAGCTGCCCGTGCCCTGGCCCACCCTCGTGACCACCCTGACCTACGG
CGTGCAGTGCTTCAGCCGCTACCCCGACCACATGAAGCAGCACGACTTCTT
CAAGTCCGCCATGCCCGAAGGCTACGTCCAGGAGCGCACCATCTTCTTCAA
GGACGACGGCAACTACAAGACCCGCGCCGAGGTGAAGTTCGAGGGCGAC
ACCCTGGTGAACCGCATCGAGCTGAAGGGCATCGACTTCAAGGAGGACGG
CAACATCCTGGGGCACAAGCTGGAGTACAACTACAACAGCCACAACGTCT
ATATCATGGCCGACAAGCAGAAGAACGGCATCAAGGTGAACTTCAAGATC
CGCCACAACATCGAGGACGGCAGCGTGCAGCTCGCCGACCACTACCAGCA
GAACACCCCCATCGGCGACGGCCCCGTGCTGCTGCCCGACAACCACTACC
TGAGCACCCAGTCCGCCCTGAGCAAAGACCCCAACGAGAAGCGCGATCAC
ATGGTCCTGCTGGAGTTCGTGACCGCCGCCGGGATCACTCTCGGCATGGAC
GAGCTGTACAAGTAA-3’

**pLenti-dCas13-ADAR1$_{DD}$**

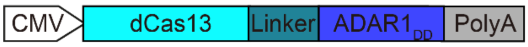


5’-ATGGTG GATTACAAGGATGACGACGATAAG(Flag tag)
ATGAAAGTGACGAAGGTAGGAGGCATTTCGCATAAGAAGTACACGTCCGA
AGGCCGCTTAGTGAAGTCAGAATCGGAAGAAAATCGCACAGACGAACGTC
TGTCGGCGTTGCTTAATATGCGCCTTGACATGTATATCAAGAATCCCAGCAG
CACGGAAACCAAGGAAAATCAAAAACGCATTGGGAAATTAAAGAAATTCT
TCTCAAACAAAATGGTCTATCTTAAAGACAATACCTTGAGTTTGAAGAATG
GGAAAAAGGAGAACATTGATCGTGAGTATTCTGAGACTGACATCCTTGAGA
GCGATGTCCGTGACAAGAAAAACTTCGCCGTGTTGAAAAAGATCTATCTGA
ATGAAAACGTGAACTCGGAGGAATTGGAAGTTTTTCGTAACGACATTAAGA
AGAAACTGAACAAAATCAACAGCCTGAAGTACTCATTTGAAAAGAATAAG
GCGAATTATCAAAAGATTAATGAGAATAACATCGAGAAGGTTGAAGGTAAG
TCAAAGCGTAACATTATTTACGATTATTATCGTGAGTCAGCGAAACGTGACG
CTTATGTAAGCAATGTGAAAGAAGCCTTTGATAAGCTTTACAAGGAAGAGG
ACATTGCAAAACTTGTTCTTGAAATTGAGAACCTTACGAAGTTAGAAAT
ACAAGATTCGCGAGTTCTACCACGAAATTATTGGACGTAAGAATGACAAGG
AAAACTTTGCAAAAATCATCTACGAAGAAATCCAGAATGTTAATAACATGA
AAGAGTTGATCGAGAAGGTACCGGACATGAGTGAGTTGAAAAAGAGCCAA
GTATTTTACAAGTATTACTTAGACAAAGAAGAGTTGAACGACAAGAACATC
AAATACGCGTTTTGTCATTTCGTGGAAATCGAAATGAGTCAGTTGCTGAAG

AACTACGTATATAAGCGCTTAAGTAATATCTCGAATGACAAAATTAAGCGTA
TCTTTGAATACCAGAACTTGAAAAAATTGATCGAAAATAAGCTGTTAAACA
AACTTGACACGTACGTCCGTAATTGTGGAAAGTATAATTATTATTTGCAAGA
CGGCGAAATTGCCACTTCAGATTTCATCGCCCGCAACCGTCAGAATGAAGC
GTTTCTTCGCAACATCATTGGGGTGTCATCTGTGGCCTACTTTTCTCTTCGC
AACATTCTTGAAACGGAGAACGAGAATGATATTACTGGGCGTATGCGCGGC
AAAACAGTTAAGAACAATAAAGGTGAAGAGAAGTACGTGTCCGGAGAAGT
TGATAAGATCTATAATGAAAATAAGAAGAACGAGGTTAAGGAGAACTTAAA
AATGTTCTATTCGTACGATTTCAATATGGACAACAAGAATGAAATCGAAGAT
TTCTTCGCCAACATCGACGAGGCGATTTCTTCCATCGCTCACGGTATTGTCG
CCTTCAACTTGGAATTAGAAGGTAAGGATATCTTTGCGTTCAAGAACATTGC
GCCATCCGAAATCTCAAAGAAGATGTTTCAGAATGAGATTAACGAGAAAA
AACTGAAATTGAAGATCTTTCGTCAACTGAACTCTGCCAACGTGTTCCGCT
ATCTCGAAAAGTATAAAATTCTGAATTACCTTAAACGTACACGCTTCGAGTT
TGTCAATAAAAATATCCCATTCGTCCCGTCTTTCACCAAATTATATTCGCGCA
TTGATGACCTGAAGAATAGTCTTGGGATTTACTGGAAAACTCCGAAAACAA
ACGACGACAATAAGACTAAGGAGATTATTGATGCCCAAATCTATTTGCTTAA
AAACATCTATTACGGGGAGTTCCTGAATTATTTCATGTCGAACAATGGTAAT
TTCTTTGAGATTTCTAAAGAAATCATCGAATTGAACAAGAACGATAAACGC
AACTTAAAGACTGGGTTTTACAAGCTGCAAAAGTTTGAAGACATCCAGGA
GAAGATTCCAAAGGAATACTTGGCGAATATCCAGTCCCTGTACATGATTAAT
GCCGGTAATCAGGACGAAGAAGAAAAGGACACTTATATTGATTTCATTCAA
AAGATCTTCTTAAAGGGATTTATGACGTATCTTGCTAATAACGGTCGTTTAA
GTCTGATTTACATCGGCTCGGATGAAGAAACAAATACGTCATTAGCAGAAA
AGAAGCAAGAGTTTGACAAGTTCTTGAAGAAGTACGAGCAGAACAATAAT
ATCAAGATCCCCTATGAGATCAATGAATTCCTGCGTGAGATCAAACTGGGA
AACATCCTGAAGTATACTGAGCGTTTAAACATGTTCTACCTTATCTTAAAGC
TTTTGAATCACAAGGAGCTGACAAATCTGAAGGGTAGTCTTGAAAAATATC
AGTCTGCCAATAAGGAAGAAGCGTTCTCTGACCAATTGGAGTTAATTAACC
TGCTTAACCTTGACAACAACCGCGTGACGGAAGACTTCGAATTAGAGGCC
GACGAGATTGGAAAATTTCTTGATTTCAATGGCAACAAAGTTAAGGATAAC
AAGGAACTGAAAAAGTTCGATACAAACAAGATCTACTTTGACGGCGAGAA
CATTATCAAACACCGTGCCTTCTACAATATTAAGAAATATGGCATGTTAAAC
TTACTGGAGAAAATTGCCGACAAGGCTGGATACAAGATCTCGATCGAAGA
GCTGAAGAAATACTCCAATAAAAAGAATGAGATCGAGAAGAACCATAAGA
TGCAGGAAAATCTGCACCGCAAATACGCTCGTCCCCGTAAAGACGAGAAG
TTTACAGATGAGGACTATGAAAGTTACAAGCAAGCTATTGAGAATATTGAG
GAGTACACCCACCTTAAGAACAAGGTAGAATTCAATGAGCTGAATTTACTG
CAGGGCCTGTTGCTGCGCATTTTACATCGTTTAGTCGGATATACCTCAATTT
GGGAACGCGATCTGCGCTTCCGCCTTAAAGGTGAGTTCCCAGAAAACCAA
TACATCGAAGAGATCTTCAACTTTGAAAATAAGAAGAACGTGAAGTACAA
AGGGGGTCAGATTGTAGAGAAATACATTAAATTCTACAAGGAATTACATCA
AAATGATGAAGTTAAGATCAACAAGTACAGTTCCGCGAATATCAAGGTGTT
GAAGCAAGAAAAGAAGGACCTTTATATTGCTAATTACATCGCCGCATTCAAT

TATATTCCTCACGCCGAGATCTCACTGCTGGAAGTCCTTGAAAATTTGCGTA
AATTGCTGTCCTACGATCGCAAACTGAAAAATGCCGTAATGAAATCAGTAG
TTGATATCCTTAAGGAGTATGGTTTTGTAGCCACATTCAAAATCGGGGCGGA
CAAGAAGATCGGTATTCAGACACTGGAGAGCGAAAAAATCGTGCATCTTA
AGAATCTTAAGAAGAAGAAGTTAATGACTGACCGCAATTCCGAGGAACTTT
GCAAATTGGTGAAGATTATGTTTGAATACAAAATGGAAGAGAAAAAGTCTG
AAAAC GGCGCGCC A GGCGGAGGAGGCAGC GGCGGAGGAGGCAGC
CTCCTCCTCTCAAGGTCCCCAGAAGCACAGCCAAAGACACTCCCTCTCACT
GGCAGCACCTTCCATGACCAGATAGCCATGCTGAGCCACCGGTGCTTCAAC
ACTCTGACTAACAGCTTCCAGCCCTCCTTGCTCGGCCGCAAGATTCTGGCC
GCCATCATTATGAAAAAAGACTCTGAGGACATGGGTGTCGTCGTCAGCTTG
GGAACAGGGAATCGCTGTGTAAAAGGAGATTCTCTCAGCCTAAAAGGAGA
AACTGTCAATGACTGCCATGCAGAAATAATCTCCCGGAGAGGCTTCATCAG
GTTTCTCTACAGTGAGTTAATGAAATACAACTCCCAGACTGCGAAGGATAG
TATATTTGAACCTGCTAAGGGAGGAGAAAAGCTCCAAATAAAAAAGACTGT
GTCATTCCATCTGTATATCAGCACTGCTCCGTGTGGAGATGGCGCCCTCTTT
GACAAGTCCTGCAGCGACCGTGCTATGGAAAGCACAGAATCCCGCCACTA
CCCTGTCTTCGAGAATCCCAAACAAGGAAAGCTCCGCACCAAGGTGGAGA
ACGGACAAGGCACAATCCCTGTGGAATCCAGTGACATTGTGCCTACGTGGG
ATGGCATTCGGCTCGGGGAGAGACTCCGTACCATGTCCTGTAGTGACAAAA
TCCTACGCTGGAACGTGCTGGGCCTGCAAGGGGCACTGTTGACCCACTTCC
TGCAGCCCATTTATCTCAAATCTGTCACATTGGGTTACCTTTTCAGCCAAGG
GCATCTGACCCGTGCTATTTGCTGTCGTGTGACAAGAGATGGGAGTGCATT
TGAGGATGGACTACGACATCCCTTTATTGTCAACCACCCCAAGGTTGGCAG
AGTCAGCATATATGATTCCAAAAGGCAATCCGGGAAGACTAAGGAGACAA
GCGTCAACTGGTGTCTGGCTGATGGCTATGACCTGGAGATCCTGGACGGTA
CCAGAGGCACTGTGGATGGGCCACGGAATGAATTGTCCCGGGTCTCCAAA
AAGAACATTTTTCTTCTATTTAAGAAGCTCTGCTCCTTCCGTTACCGCAGGG
ATCTACTGAGACTCTCCTATGGTGAGGCCAAGAAAGCTGCCCGTGACTACG
AGACGGCCAAGAACTACTTCAAAAAAGGCCTGAAGGATATGGGCTATGGG
AACTGGATTAGCAAACCCCAGGAGGAAAAGAACTTTTATCTCTGCCCAGTA
TAG-3’

**ADAR1(p110) cDNA**

5’-
ATGGCCGAGATCAAGGAGAAAATCTGCGACTATCTCTTCAATGTGTCTGAC
TCCTCTGCCCTGAATTTGGCTAAAAATATTGGCCTTACCAAGGCCCGAGATA
TAAATGCTGTGCTAATTGACATGGAAAGGCAGGGGGATGTCTATAGACAAG
GGACAACCCCTCCCATATGGCATTTGACAGACAAGAAGCGAGAGAGGATG
CAAATCAAGAGAAATACGAACAGTGTTCCTGAAACCGCTCCAGCTGCAAT
CCCTGAGACCAAAAGAAACGCAGAGTTCCTCACCTGTAATATACCCACATC
AAATGCCTCAAATAACATGGTAACCACAGAAAAAGTGGAGAATGGGCAGG
AACCTGTCATAAAGTTAGAAAACAGGCAAGAGGCCAGACCAGAACCAGC
AAGACTGAAACCACCTGTTCATTACAATGGCCCCTCAAAAGCAGGGTATGT

TGACTTTGAAAATGGCCAGTGGGCCACAGATGACATCCCAGATGACTTGAA
TAGTATCCGCGCAGCACCAGGTGAGTTTCGAGCCATCATGGAGATGCCCTC
CTTCTACAGTCATGGCTTGCCACGGTGTTCACCCTACAAGAAACTGACAGA
GTGCCAGCTGAAGAACCCCATCAGCGGGCTGTTAGAATATGCCCAGTTCGC
TAGTCAAACCTGTGAGTTCAACATGATAGAGCAGAGTGGACCACCCCATGA
ACCTCGATTTAAATTCCAGGTTGTCATCAATGGCCGAGAGTTTCCCCCAGCT
GAAGCTGGAAGCAAGAAAGTGGCCAAGCAGGATGCAGCTATGAAAGCCAT
GACAATTCTGCTAGAGGAAGCCAAAGCCAAGGACAGTGGAAAATCAGAA
GAATCATCCCACTATTCCACAGAGAAAGAATCAGAGAAGACTGCAGAGTC
CCAGACCCCCACCCCTTCAGCCACATCCTTCTTTTCTGGGAAGAGCCCCGT
CACCACACTGCTTGAGTGTATGCACAAATTGGGGAACTCCTGCGAATTCCG
TCTCCTGTCCAAAGAAGGCCCTGCCCATGAACCCAAGTTCCAATACTGTGT
TGCAGTGGGAGCCCAAACTTTCCCCAGTGTGAGTGCTCCCAGCAAGAAAG
TGGCAAAGCAGATGGCCGCAGAGGAAGCCATGAAGGCCCTGCATGGGGA
GGCGACCAACTCCATGGCTTCTGATAACCAGCCTGAAGGTATGATCTCAGA
GTCACTTGATAACTTGGAATCCATGATGCCCAACAAGGTCAGGAAGATTGG
CGAGCTCGTGAGATACCTGAACACCAACCCTGTGGGTGGCCTTTTGGAGTA
CGCCCGCTCCCATGGCTTTGCTGCTGAATTCAAGTTGGTCGACCAGTCCGG
ACCTCCTCACGAGCCCAAGTTCGTTTACCAAGCAAAAGTTGGGGGTCGCT
GGTTCCCAGCCGTCTGCGCACACAGCAAGAAGCAAGGCAAGCAGGAAGC
AGCAGATGCGGCTCTCCGTGTCTTGATTGGGGAGAACGAGAAGGCAGAAC
GCATGGGTTTCACAGAGGTAACCCCAGTGACAGGGGCCAGTCTCAGAAGA
ACTATGCTCCTCCTCTCAAGGTCCCCAGAAGCACAGCCAAAGACACTCCCT
CTCACTGGCAGCACCTTCCATGACCAGATAGCCATGCTGAGCCACCGGTGC
TTCAACACTCTGACTAACAGCTTCCAGCCCTCCTTGCTCGGCCGCAAGATT
CTGGCCGCCATCATTATGAAAAAAGACTCTGAGGACATGGGTGTCGTCGTC
AGCTTGGGAACAGGGAATCGCTGTGTAAAAGGAGATTCTCTCAGCCTAAA
AGGAGAAACTGTCAATGACTGCCATGCAGAAATAATCTCCCGGAGAGGCTT
CATCAGGTTTCTCTACAGTGAGTTAATGAAATACAACTCCCAGACTGCGAA
GGATAGTATATTTGAACCTGCTAAGGGAGGAGAAAAGCTCCAAATAAAAAA
GACTGTGTCATTCCATCTGTATATCAGCACTGCTCCGTGTGGAGATGGCGCC
CTCTTTGACAAGTCCTGCAGCGACCGTGCTATGGAAAGCACAGAATCCCGC
CACTACCCTGTCTTCGAGAATCCCAAACAAGGAAAGCTCCGCACCAAGGT
GGAGAACGGAGAAGGCACAATCCCTGTGGAATCCAGTGACATTGTGCCTA
CGTGGGATGGCATTCGGCTCGGGGAGAGACTCCGTACCATGTCCTGTAGTG
ACAAAATCCTACGCTGGAACGTGCTGGGCCTGCAAGGGGCACTGTTGACC
CACTTCCTGCAGCCCATTTATCTCAAATCTGTCACATTGGGTTACCTTTTCA
GCCAAGGGCATCTGACCCGTGCTATTTGCTGTCGTGTGACAAGAGATGGGA
GTGCATTTGAGGATGGACTACGACATCCCTTTATTGTCAACCACCCCAAGG
TTGGCAGAGTCAGCATATATGATTCCAAAAGGCAATCCGGGAAGACTAAGG
AGACAAGCGTCAACTGGTGTCTGGCTGATGGCTATGACCTGGAGATCCTGG
ACGGTACCAGAGGCACTGTGGATGGGCCACGGAATGAATTGTCCCGGGTC
TCCAAAAAGAACATTTTTCTTCTATTTAAGAAGCTCTGCTCCTTCCGTTACC
GCAGGGATCTACTGAGACTCTCCTATGGTGAGGCCAAGAAAGCTGCCCGT

GACTACGAGACGGCCAAGAACTACTTCAAAAAAGGCCTGAAGGATATGGG
CTATGGGAACTGGATTAGCAAACCCCAGGAGGAAAAGAACTTTTATCTCTG
CCCAGTA GATTACAAGGATGACGACGATAAG(Flag tag) TAG-3’

**ADAR1(p150) cDNA**
5’-
ATGAATCCGCGGCAGGGGTATTCCCTCAGCGGATACTACACCCATCCATTTC
AAGGCTATGAGCACAGACAGCTCAGATACCAGCAGCCTGGGCCAGGATCT
TCCCCCAGTAGTTTCCTGCTTAAGCAAATAGAATTTCTCAAGGGGCAGCTC
CCAGAAGCACCGGTGATTGGAAAGCAGACACCGTCACTGCCACCTTCCCT
CCCAGGACTCCGGCCAAGGTTTCCAGTACTACTTGCCTCCAGTACCAGAGG
CAGGCAAGTGGACATCAGGGGTGTCCCCAGGGGCGTGCATCTCGGAAGTC
AGGGGCTCCAGAGAGGGTTCCAGCATCCTTCACCACGTGGCAGGAGTCTG
CCACAGAGAGGTGTTGATTGCCTTTCCTCACATTTCCAGGAACTGAGTATC
TACCAAGATCAGGAACAAAGGATCTTAAAGTTCCTGGAAGAGCTTGGGGA
AGGGAAGGCCACCACAGCACATGATCTGTCTGGGAAACTTGGGACTCCGA
AGAAAGAAATCAATCGAGTTTTATACTCCCTGGCAAAGAAGGGCAAGCTAC
AGAAAGAGGCAGGAACACCCCCTTTGTGGAAAATCGCGGTCTCCACTCAG
GCTTGGAACCAGCACAGCGGAGTGGTAAGACCAGACGGTCATAGCCAAGG
AGCCCCAAACTCAGACCCGAGTTTGGAACCGGAAGACAGAAACTCCACAT
CTGTCTCAGAAGATCTTCTTGAGCCTTTTATTGCAGTCTCAGCTCAGGCTTG
GAACCAGCACAGCGGAGTGGTAAGACCAGACAGTCATAGCCAAGGATCCC
CAAACTCAGACCCAGGTTTGGAACCTGAAGACAGCAACTCCACATCTGCC
TTGGAAGATCCTCTTGAGTTTTTAGACATGGCCGAGATCAAGGAGAAAATC
TGCGACTATCTCTTCAATGTGTCTGACTCCTCTGCCCTGAATTTGGCTAAAA
ATATTGGCCTTACCAAGGCCCGAGATATAAATGCTGTGCTAATTGACATGGA
AAGGCAGGGGGATGTCTATAGACAAGGGACAACCCCTCCCATATGGCATTT
GACAGACAAGAAGCGAGAGAGGATGCAAATCAAGAGAAATACGAACAGT
GTTCCTGAAACCGCTCCAGCTGCAATCCCTGAGACCAAAAGAAACGCAGA
GTTCCTCACCTGTAATATACCCACATCAAATGCCTCAAATAACATGGTAACC
ACAGAAAAAGTGGAGAATGGGCAGGAACCTGTCATAAAGTTAGAAAACA
GGCAAGAGGCCAGACCAGAACCAGCAAGACTGAAACCACCTGTTCATTAC
AATGGCCCCTCAAAAGCAGGGTATGTTGACTTTGAAAATGGCCAGTGGGCC
ACAGATGACATCCCAGATGACTTGAATAGTATCCGCGCAGCACCAGGTGAG
TTTCGAGCCATCATGGAGATGCCCTCCTTCTACAGTCATGGCTTGCCACGGT
GTTCACCCTACAAGAAACTGACAGAGTGCCAGCTGAAGAACCCCATCAGC
GGGCTGTTAGAATATGCCCAGTTCGCTAGTCAAACCTGTGAGTTCAACATG
ATAGAGCAGAGTGGACCACCCCATGAACCTCGATTTAAATTCCAGGTTGTC
ATCAATGGCCGAGAGTTTCCCCCAGCTGAAGCTGGAAGCAAGAAAGTGGC
CAAGCAGGATGCAGCTATGAAAGCCATGACAATTCTGCTAGAGGAAGCCA
AAGCCAAGGACAGTGGAAAATCAGAAGAATCATCCCACTATTCCACAGAG
AAAGAATCAGAGAAGACTGCAGAGTCCCAGACCCCCACCCCTTCAGCCAC
ATCCTTCTTTTCTGGGAAGAGCCCCGTCACCACACTGCTTGAGTGTATGCA

CAAATTGGGGAACTCCTGCGAATTCCGTCTCCTGTCCAAAGAAGGCCCTGC
CCATGAACCCAAGTTCCAATACTGTGTTGCAGTGGGAGCCCAAACTTTCCC
CAGTGTGAGTGCTCCCAGCAAGAAAGTGGCAAAGCAGATGGCCGCAGAG
GAAGCCATGAAGGCCCTGCATGGGGAGGCGACCAACTCCATGGCTTCTGA
TAACCAGCCTGAAGGTATGATCTCAGAGTCACTTGATAACTTGGAATCCATG
ATGCCCAACAAGGTCAGGAAGATTGGCGAGCTCGTGAGATACCTGAACAC
CAACCCTGTGGGTGGCCTTTTGGAGTACGCCCGCTCCCATGGCTTTGCTGC
TGAATTCAAGTTGGTCGACCAGTCCGGACCTCCTCACGAGCCCAAGTTCGT
TTACCAAGCAAAAGTTGGGGGTCGCTGGTTCCCAGCCGTCTGCGCACACA
GCAAGAAGCAAGGCAAGCAGGAAGCAGCAGATGCGGCTCTCCGTGTCTTG
ATTGGGGAGAACGAGAAGGCAGAACGCATGGGTTTCACAGAGGTAACCCC
AGTGACAGGGGCCAGTCTCAGAAGAACTATGCTCCTCCTCTCAAGGTCCCC
AGAAGCACAGCCAAAGACACTCCCTCTCACTGGCAGCACCTTCCATGACC
AGATAGCCATGCTGAGCCACCGGTGCTTCAACACTCTGACTAACAGCTTCC
AGCCCTCCTTGCTCGGCCGCAAGATTCTGGCCGCCATCATTATGAAAAAAG
ACTCTGAGGACATGGGTGTCGTCGTCAGCTTGGGAACAGGGAATCGCTGT
GTAAAAGGAGATTCTCTCAGCCTAAAAGGAGAAACTGTCAATGACTGCCAT
GCAGAAATAATCTCCCGGAGAGGCTTCATCAGGTTTCTCTACAGTGAGTTA
ATGAAATACAACTCCCAGACTGCGAAGGATAGTATATTTGAACCTGCTAAG
GGAGGAGAAAAGCTCCAAATAAAAAAGACTGTGTCATTCCATCTGTATATC
AGCACTGCTCCGTGTGGAGATGGCGCCCTCTTTGACAAGTCCTGCAGCGA
CCGTGCTATGGAAAGCACAGAATCCCGCCACTACCCTGTCTTCGAGAATCC
CAAACAAGGAAAGCTCCGCACCAAGGTGGAGAACGGAGAAGGCACAATC
CCTGTGGAATCCAGTGACATTGTGCCTACGTGGGATGGCATTCGGCTCGGG
GAGAGACTCCGTACCATGTCCTGTAGTGACAAAATCCTACGCTGGAACGTG
CTGGGCCTGCAAGGGGCACTGTTGACCCACTTCCTGCAGCCCATTTATCTC
AAATCTGTCACATTGGGTTACCTTTTCAGCCAAGGGCATCTGACCCGTGCT
ATTTGCTGTCGTGTGACAAGAGATGGGAGTGCATTTGAGGATGGACTACGA
CATCCCTTTATTGTCAACCACCCCAAGGTTGGCAGAGTCAGCATATATGATT
CCAAAAGGCAATCCGGGAAGACTAAGGAGACAAGCGTCAACTGGTGTCTG
GCTGATGGCTATGACCTGGAGATCCTGGACGGTACCAGAGGCACTGTGGAT
GGGCCACGGAATGAATTGTCCCGGGTCTCCAAAAAGAACATTTTTCTTCTA
TTTAAGAAGCTCTGCTCCTTCCGTTACCGCAGGGATCTACTGAGACTCTCCT
ATGGTGAGGCCAAGAAAGCTGCCCGTGACTACGAGACGGCCAAGAACTAC
TTCAAAAAAGGCCTGAAGGATATGGGCTATGGGAACTGGATTAGCAAACCC
CAGGAGGAAAAGAACTTTTATCTCTGCCCAGTA
GATTACAAGGATGACGACGATAAG(Flag tag) TAG-3’

**ADAR2 cDNA**
5’-
ATGGATATAGAAGATGAAGAAAACATGAGTTCCAGCAGCACTGATGTGAAG
GAAAACCGCAATCTGGACAACGTGTCCCCCAAGGATGGCAGCACACCTGG
GCCTGGCGAGGGCTCTCAGCTCTCCAATGGGGGTGGTGGTGGCCCCGGCA
GAAAGCGGCCCCTGGAGGAGGGCAGCAATGGCCACTCCAAGTACCGCCTG

AAGAAAAGGAGGAAAACACCAGGGCCCGTCCTCCCCAAGAACGCCCTGA
TGCAGCTGAATGAGATCAAGCCTGGTTTGCAGTACACACTCCTGTCCCAGA
CTGGGCCCGTGCACGCGCCTTTGTTTGTCATGTCTGTGGAGGTGAATGGCC
AGGTTTTTGAGGGCTCTGGTCCCACAAAGAAAAAGGCAAAACTCCATGCT
GCTGAGAAGGCCTTGAGGTCTTTCGTTCAGTTTCCTAATGCCTCTGAGGCC
CACCTGGCCATGGGGAGGACCCTGTCTGTCAACACGGACTTCACATCTGAC
CAGGCCGACTTCCCTGACACGCTCTTCAATGGTTTTGAAACTCCTGACAAG
GCGGAGCCTCCCTTTTACGTGGGCTCCAATGGGGATGACTCCTTCAGTTCC
AGCGGGGACCTCAGCTTGTCTGCTTCCCCGGTGCCTGCCAGCCTAGCCCAG
CCTCCTCTCCCTGCCTTACCACCATTCCCACCCCCGAGTGGGAAGAATCCC
GTGATGATCTTGAACGAACTGCGCCCAGGACTCAAGTATGACTTCCTCTCC
GAGAGCGGGGAGAGCCATGCCAAGAGCTTCGTCATGTCTGTGGTCGTGGA
TGGTCAGTTCTTTGAAGGCTCGGGGAGAAACAAGAAGCTTGCCAAGGCCC
GGGCTGCGCAGTCTGCCCTGGCCGCCATTTTTAACTTGCACTTGGATCAGA
CGCCATCTCGCCAGCCTATTCCCAGTGAGGGTCTTCAGCTGCATTTACCGCA
GGTTTTAGCTGACGCTGTCTCACGCCTGGTCCTGGGTAAGTTTGGTGACCT
GACCGACAACTTCTCCTCCCCTCACGCTCGCAGAAAAGTGCTGGCTGGAG
TCGTCATGACAACAGGCACAGATGTTAAAGATGCCAAGGTGATAAGTGTTT
CTACAGGAACAAAATGTATTAATGGTGAATACATGAGTGATCGTGGCCTTGC
ATTAAATGACTGCCATGCAGAAATAATATCTCGGAGATCCTTGCTCAGATTT
CTTTATACACAACTTGAGCTTTACTTAAATAACAAAGATGATCAAAAAAGAT
CCATCTTTCAGAAATCAGAGCGAGGGGGGTTTAGGCTGAAGGAGAATGTC
CAGTTTCATCTGTACATCAGCACCTCTCCCTGTGGAGATGCCAGAATCTTCT
CACCACATGAGCCAATCCTGGAAGAACCAGCAGATAGACACCCAAATCGT
AAAGCAAGAGGACAGCTACGGACCAAAATAGAGTCTGGTGAGGGGACGA
TTCCAGTGCGCTCCAATGCGAGCATCCAAACGTGGGACGGGGTGCTGCAA
GGGGAGCGGCTGCTCACCATGTCCTGCAGTGACAAGATTGCACGCTGGAA
CGTGGTGGGCATCCAGGGATCCCTGCTCAGCATTTTCGTGGAGCCCATTTA
CTTCTCGAGCATCATCCTGGGCAGCCTTTACCACGGGGACCACCTTTCCAG
GGCCATGTACCAGCGGATCTCCAACATAGAGGACCTGCCACCTCTCTACAC
CCTCAACAAGCCTTTGCTCAGTGGCATCAGCAATGCAGAAGCACGGCAGC
CAGGGAAGGCCCCCAACTTCAGTGTCAACTGGACGGTAGGCGACTCCGCT
ATTGAGGTCATCAACGCCACGACTGGGAAGGATGAGCTGGGCCGCGCGTC
CCGCCTGTGTAAGCACGCGTTGTACTGTCGCTGGATGCGTGTGCACGGCAA
GGTTCCCTCCCACTTACTACGCTCCAAGATTACCAAACCCAACGTGTACCAT
GAGTCCAAGCTGGCGGCAAAGGAGTACCAGGCCGCCAAGGCGCGTCTGTT
CACAGCCTTCATCAAGGCGGGGCTGGGGGCCTGGGTGGAGAAGCCCACCG
AGCAGGACCAGTTCTCACTCACGCCC
GATTACAAGGATGACGACGATAAG(Flag tag) TAG-3’

**pLenti-MCS-mCherry backbone**

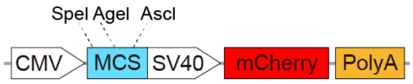

5’-
cgataagcttgggagttccgcgttacataacttacggtaaatggcccgcctggctgaccgcccaacgacccccgcccattg
acgtcaataatgacgtatgttcccatagtaacgccaatagggactttccattgacgtcaatgggtggagtatttacggtaaact
gcccacttggcagtacatcaagtgtatcatatgccaagtacgccccctattgacgtcaatgacggtaaatggcccgcctggc
attatgcccagtacatgaccttatgggactttcctacttggcagtacatctacgtattagtcatcgctattaccatggtgatgcgg
ttttggcagtacatcaatgggcgtggatagcggtttgactcacggggatttccaagtctccaccccattgacgtcaatgggag
tttgttttggcaccaaaatcaacgggactttccaaaatgtcgtaacaactccgccccattgacgcaaatgggcggtaggcgt
gtacggtgggaggtctatataagcagagctcgtttagtgaaccgtcagatcgcctggagacgccatccacgctgttttgacc
tccatagaagacaccgactctagaggatccgg**actagt**tt**accggt**gggggccccc**ggcgcgcc**ggtgtacaccctgca
ggggtttaaacccacgcgtcgaccagtggtcgaccctgtggaatgtgtgtcagttagggtgtggaaagtccccaggctccc
cagcaggcagaagtatgcaaagcatgcatctcaattagtcagcaaccaggtgtggaaagtccccaggctccccagcagg
cagaagtatgcaaagcatgcatctcaattagtcagcaaccatagtcccgcccctaactccgcccatcccgcccctaactccg
cccagttccgcccattctccgccccatggctgactaattttttttatttatgcagaggccgaggccgcctcggcctctgagctat
tccagaagtagtgaggaggcttttttggaggcctaggcttttgcaaaaagctatcgctagctcgagatggtgagcaagggc
gaggaggataacatggccatcatcaaggagttcatgcgcttcaaggtgcacatggagggctccgtgaacggccacgagtt
cgagatcgagggcgagggcgagggccgcccctacgagggcacccagaccgccaagctgaaggtgaccaagggtgg
ccccctgcccttcgcctgggacatcctgtcccctcagttcatgtacggctccaaggcctacgtgaagcaccccgccgacat
ccccgactacttgaagctgtccttccccgagggcttcaagtgggagcgcgtgatgaacttcgaggacggcggcgtggtga
ccgtgacccaggactcctccctgcaggacggcgagttcatctacaaggtgaagctgcgcggcaccaacttcccctccgac
ggccccgtaatgcagaagaagaccatgggctgggaggcctcctccgagcggatgtaccccgaggacggcgccctgaa
gggcgagatcaagcagaggctgaagctgaaggacggcggccactacgacgctgaggtcaagaccacctacaaggcca
agaagcccgtgcagctgcccggcgcctacaacgtcaacatcaagttggacatcacctcccacaacgaggactacaccatc
gtggaacagtacgaacgcgccgagggccgccactccaccggcggcatggacgagctgtacaagtaagctaagcacttc
gtggccgaggagcaggactgagaattccagtcgacaatcaacctctggattacaaaatttgtgaaagattgactggtattctt
aactatgttgctccttttacgctatgtggatacgctgctttaatgcctttgtatcatgctattgcttcccgtatggctttcattttctcct
ccttgtataaatcctggttgctgtctctttatgaggagttgtggcccgttgtcaggcaacgtggcgtggtgtgcactgtgtttgct
gacgcaacccccactggttggggcattgccaccacctgtcagctcctttccgggactttcgctttccccctccctattgccac
ggcggaactcatcgccgcctgccttgcccgctgctggacaggggctcggctgttgggcactgacaattccgtggtgttgtc
ggggaagctgacgtcctttccatggctgctcgcctgtgttgccacctggattctgcgcgggacgtccttctgctacgtccctt
cggccctcaatccagcggaccttccttcccgcggcctgctgccggctctgcggcctcttccgcgtcttcgccttcgccctca
gacgagtcggatctccctttgggccgcctccccgcctggaattcgagctcggtacctttaagaccaatgacttacaaggcag
ctgtagatcttagccactttttaaaagaaaaggggggactggaagggctaattcactcccaacgaagacaagatctgcttttt
gcttgtactgggtctctctggttagaccagatctgagcctgggagctctctggctaactagggaacccactgcttaagcctca
ataaagcttgccttgagtgcttcaagtagtgtgtgcccgtctgttgtgtgactctggtaactagagatccctcagacccttttagt
cagtgtggaaaatctctagcagtagtagttcatgtcatcttattattcagtatttataacttgcaaagaaatgaatatcagagagt
gagaggaacttgtttattgcagcttataatggttacaaataaagcaatagcatcacaaatttcacaaataaagcatttttttcact
gcattctagttgtggtttgtccaaactcatcaatgtatcttatcatgtctggctctagctatcccgcccctа-3’

**pLenti-arRNA-BFP backbone**

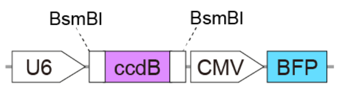

5’-
gagggcctatttcccatgattccttcatatttgcatatacgatacaaggctgttagagagataattagaattaatttgactgtaaa
cacaaagatattagtacaaaatacgtgacgtagaaagtaataatttcttgggtagtttgcagttttaaaattatgttttaaaatgga
ctatcatatgcttaccgtaacttgaaagtatttcgatttcttggctttatatatcttgtggaaaggacgaaacaccga**gagacg**ct
ggcttatcgaaattaatacgactcactatagggagacccaagctggctagttaagctatcaacaagtttgtacaaaaaagctg
aacgagaaacgtaaaatgatataaatatcaatatattaaattagattttgcataaaaaacagactacataatactgtaaaacaca
acatatccagtcactatgaatcaactacttagatggtattagtgacctgtagtcgaccgacagccttccaaatgttcttcgggtg
atgctgccaacttagtcgaccgacagccttccaaatgttcttctcaaacggaatcgtcgtatccagcctactcgctattgtcctc
aatgccgtattaaatcataaaaagaaataagaaaaagaggtgcgagcctcttttttgtgtgacaaaataaaaacatctacctat
tcatatacgctagtgtcatagtcctgaaaatcatctgcatcaagaacaatttcacaactcttatacttttctcttacaagtcgttcg
gcttcatctggattttcagcctctatacttactaaacgtgataaagtttctgtaatttctactgtatcgacctgcagactggctgtgt
ataagggagcctgacatttatattccccagaacatcaggttaatggcgtttttgatgtcattttcgcggtggctgagatcagcca
cttcttccccgataacggagaccggcacactggccatatcggtggtcatcatgcgccagctttcatccccgatatgcaccac
cgggtaaagttcacgggagactttatctgacagcagacgtgcactggccaggggggatcaccatccgtcgcccgggcgtgt
caataatatcactctgtacatccacaaacagacgataacggctctctcttttataggtgtaaaccttaaactgcatttcaccagc
ccctgttctcgtcagcaaaagagccgttcatttcaataaaccgggcgacctcagccatcccttcctgattttccgctttccagc
gttcggcacgcagacgacgggcttcattctgcatggttgtgcttaccagaccggagatattgacatcatatatgccttgagca
actgatagctgtcgctgtcaactgtcactgtaatacgctgcttcatagcatacctctttttgacatacttcgggtatacatatcagt
atatattcttataccgcaaaaatcagcgcgcaaatacgcatactgttatctggcttttagtaagccggatccacgcggcgttta
cgcccccccctgccactcatcgcagtactgttgtaattcattaagcattctgccgacatggaagccatcacaaacggcatgatg
aacctgaatcgccagcggcatcagcaccttgtcgccttgcgtataatatttgcccatggtgaaaacgggggcgaagaagtt
gtccatattggccacgtttaaatcaaaactggtgaaactcacccagggattggctgagacgaaaaacatattctcaataaacc
ctttagggaaataggccaggttttcaccgtaacacgccacatcttgcgaatatatgtgtagaaactgccggaaatcgtcgtgg
tattcactccagagcgatgaaaacgtttcagtttgctcatggaaaacggtgtaacaagggtgaacactatcccatatcaccag
ctcaccgtctttcattgccatacggaattccggatgagcattcatcaggcgggcaagaatgtgaataaaggccggataaaac
ttgtgcttatttttctttacggtctttaaaaaggccgtaatatccagctgaacggtctggttataggtacattgagcaactgactga
aatgcctcaaaatgttctttacgatgccattgggatatatcaacggtggtatatccagtgatttttttctccattttagcttccttagc
tcctgaaaatctcgataactcaaaaaatacgcccggtagtgatcttatttcattatggtgaaagttggaacctcttacgtgccga
tcaacgtctcattttcgccaaaagttggcccagggcttcccggtatcaacagggacaccaggatttatttattctgcgaagtga
tcttccgtcacaggtatttattcggcgcaaagtgcgtcgggtgatgctgccaacttagtcgactacaggtcactaataccatct
aagtagttgattcatagtgactggatatgttgtgttttacagtattatgtagtctgtttttttatgcaaaatctaatttaatatattgatatt
tatatcattttacgtttctcgttcagctttcttgtacaaagtggttgatctagagggcccgcggttcgaa**cgtctc**ttgatcatatg
gcgcgccctcgaggtcgacggtatcgataagctcgcttcacgagattccagcaggtcgagggacctaataacttcgtatag
catacattatacgaagttatattaagggttccaagcttaagcggccgcgtggataaccgtattaccgccatgcattagttattaa
tagtaatcaattacggggtcattagttcatagcccatatatggagttccgcgttacataacttacggtaaatggcccgcctggc
tgaccgcccaacgacccccgcccattgacgtcaataatgacgtatgttcccatagtaacgccaatagggactttccattgac
gtcaatgggtggagtatttacggtaaactgcccacttggcagtacatcaagtgtatcatatgccaagtacgccccctattgac
gtcaatgacggtaaatggcccgcctggcattatgcccagtacatgaccttatgggactttcctacttggcagtacatctacgta
ttagtcatcgctattaccatggtgatgcggttttggcagtacatcaatgggcgtggatagcggtttgactcacggggatttcca
agtctccaccccattgacgtcaatgggagtttgttttggcaccaaaatcaacgggactttccaaaatgtcgtaacaactccgc
cccattgacgcaaatgggcggtaggcgtgtacggtgggaggtctatataagcagagctggtttagtgaaccgtcagatccg
ctagcgccaccatgagcgagctgattaaggagaacatgcacatgaagctgtacatggagggcaccgtggacaaccatca
cttcaagtgcacatccgagggcgaaggcaagccctacgagggcacccagaccatgagaatcaaggtggtcgagggcg
gccctctccccttcgccttcgacatcctggctactagcttcctctacggcagcaagaccttcatcaaccacacccagggcatc

cccgacttcttcaagcagtccttccctgagggcttcacatgggagagagtcaccacatacgaagacgggggcgtgctgac
cgctacccaggacaccagcctccaggacggctgcctcatctacaacgtcaagatcagaggggtgaacttcacatccaacg
gccctgtgatgcagaagaaaacactcggctgggaggccttcaccgagactctgtaccccgctgacggcggcctggaagg
cagaaacgacatggccctgaagctcgtgggcgggagccatctgatcgcaaacatcaagaccacatatagatccaagaaa
cccgctaagaacctcaagatgcctggcgtctactatgtggactacagactggaaagaatcaaggaggccaacaacgaga
cctacgtcgagcagcacgaggtggcagtggccagatactgcgacctccctagcaaactggggcacaaactcaattaa-3’

**Coding sequence (CDS) of the disease-relevant genes**

COL3A1

5’-

atgatgagctttgtgcaaaaggggagctggctacttctcgctctgcttcatcccactattattttggcacaacaggaagctgttg
aaggaggatgttcccatcttggtcagtcctatgcggatagagatgtctggaagccagaaccatgccaaatatgtgtctgtga
ctcaggatccgttctctgcgatgacataatatgtgacgatcaagaattagactgccccaacccagaaattccatttggagaat
gttgtgcagtttgcccacagcctccaactgctcctactcgccctcctaatggtcaaggacctcaaggccccaagggagatcc
aggccctcctggtattcctgggagaaatggtgaccctggtattccaggacaaccagggtcccctggttctcctggcccccct
ggaatctgtgaatcatgccctactggtcctcagaactattctccccagtatgattcatatgatgtcaagtctggagtagcagta
ggaggactcgcaggctatcctggaccagctggcccccccaggccctcccggtcccccctggtacatctggtcatcctggttcc
cctggatctccaggataccaaggacccccctggtgaacctgggcaagctggtccttcaggccctccaggacctcctggtgct
ataggtccatctggtcctgctggaaaagatggagaatcaggtagacccggacgacctggagagcgaggattgcctggac
ctccaggtatcaaaggtccagctgggatacctggattccctggtatgaaaggacacagaggcttcgatggacgaaatgga
gaaaagggtgaaacaggtgctcctggattaaagggtgaaaatggtcttccaggcgaaaatggagctcctggacccatggg
tccaagaggggctcctggtgagcgaggacggccaggacttcctggggctgcaggtgctcggggtaatgacggtgctcga
ggcagtgatggtcaaccaggccctcctggtcctcctggaactgccggattccctggatcccctggtgctaagggtgaagtt
ggacctgcagggtctcctggttcaaatggtgcccctggacaaagaggagaacctggacctcagggacacgctggtgctc
aaggtcctcctggccctcctgggattaatggtagtcctggtggtaaaggcgaaatgggtcccgctggcattcctggagctcc
tggactgatgggagcccggggtcctccaggaccagccggtgctaatggtgctcctggactgcgaggtggtgcaggtgag
cctggtaagaatggtgccaaaggagagcccggaccacgtggtgaacgcggtgaggctggtattccaggtgttccaggag
ctaaaggcgaagatggcaaggatggatcacctggagaacctggtgcaaatgggcttccaggagctgcaggagaaaggg
gtgcccctgggttccgaggacctgctggaccaaatggcatcccaggagaaaagggtcctgctggagagcgtggtgctcc
aggccctgcagggcccagaggagctgctggagaacctggcagagatggcgtccctggaggtccaggaatgaggggca
tgcccggaagtccaggaggaccaggaagtgatgggaaaccagggcctcccggaagtcaaggagaaagtggtcgacca
ggtcctcctgggccatctggtccccgaggtcagcctggtgtcatgggcttccccggtcctaaaggaaatgatggtgctcctg
gtaagaatggagaacgaggtggccctggaggacctggccctcagggtcctcctggaaagaatggtgaaactggacctca

gggacccccagggcctactgggcctggtggtgacaaaggagacacaggacccccctggtccacaaggattacaaggctt
gcctggtacaggtggtcctccaggagaaaatggaaaacctggggaaccaggtccaaagggtgatgccggtgcacctgg
agctccaggaggcaagggtgatgctggtgcccctggtgaacgtggacctcctggattggcaggggccccaggacttaga
ggtggagctggtccccctggtcccgaaggaggaaagggtgctgctggtcctcctgggccacctggtgctgctggtactcc
tggtctgcaaggaatgcctggagaaagaggaggtcttggaagtcctggtccaaagggtgacaagggtgaaccaggcggt
ccaggtgctgatggtgtcccagggaaagatggcccaaggggtcctactggtcctattggtcctcctggcccagctggcca
gcctggagataagggtgaaggtggtgcccccggacttccaggtatagctggacctcgtggtagccctggtgagagaggt
gaaactggccctccaggacctgctggtttccctggtgctcctggacagaatggtgaacctggtggtaaaggagaaagagg
ggctccgggtgagaaaggtgaaggaggccctcctggagttgcaggacccccctggaggttctggacctgctggtcctcctg
gtccccaaggtgtcaaaggtgaacgtggcagtcctggtggacctggtgctgctggcttccctggtgctcgtggtcttcctgg
tcctcctggtagtaatggtaacccaggacccccaggtcccagcggttctccaggcaaggatgggcccccaggtcctgcgg
gtaacactggtgctcctggcagccctggagtgtctggaccaaaaggtgatgctggccaaccaggagagaagggatcgcc
tggtgcccagggcccaccaggagctccaggcccacttgggattgctgggatcactggagcacggggtcttgcaggacca
ccaggcatgccaggtcctaggggaagccctggccctcagggtgtcaagggtgaaagtgggaaaccaggagctaacggt
ctcagtggagaacgtggtccccctggaccccagggtcttcctggtctggctggtacagctggtgaacctggaagagatgg
aaaccctggatcagatggtcttccaggccgagatggatctcctggtggcaagggtgatcgtggtgaaaatggctctcctggt
gcccctggcgctcctggtcatccaggcccacctggtcctgtcggtccagctggaaagagtggtgacagaggagaaagtg
gccctgctggccctgctggtgctcccggtcctgctggttcccgaggtgctcctggtcctcaaggcccacgtggtgacaaag
gtgaaacaggtgaacgtggagctgctggcatcaaaggacatcgaggattccctggtaatccaggtgccccaggttctcca
ggccctgctggtcagcagggtgcaatcggcagtccaggacctgcaggccccagaggacctgttggacccagtggacctc
ctggcaaagatggaaccagtggacatccaggtcccattggaccaccagggcctcgaggtaacagaggtgaaagaggat
ctgagggctccccaggccacccagggcaaccaggccctcctggacctcctggtgcccctggtccttgctgtggtggtgttg
gagccgctgccattgctgggattggaggtgaaaaagctggcggttttgccccgtattatggagatgaaccaatggatttcaa
aatcaacaccgatgagattatgacttcactcaagtctgttaatggacaaatagaaagcctcattagtcctgatggttctcgtaaa
aaccccgctagaaactgcagagacctgaaattctgccatcctgaactcaagagtggagaatactgggttgaccctaaccaa
ggatgcaaattggatgctatcaaggtattctgtaatatggaaactggggaaacatgcataagtgccaatcctttgaatgttcca
cggaaacactggtggacagattctagtgctgagaagaaacacgtttggtttggagagtccatggatggtggttttcagtttag
ctacggcaatcctgaacttcctgaagatgtccttgatgtgcagctggcattccttcgacttctctccagccgagcttcccagaa
catcacatatcactgcaaaaatagcattgcatacatggatcaggccagtggaaatgtaaagaaggccctgaagctgatggg
gtcaaatgaaggtgaattcaaggctgaaggaaatagcaaattcacctacacagttctggaggatggttgcacgaaacacac
tggggaatggagcaaaacagtctttgaatatcgaacacgcaaggctgtgagactacctattgtagatattgcaccctatgac

attggtggtcctgatcaagaatttggtgtggacgttggccctgtttgctttttataa-3’

**BMPR2**

5’-

atgacttcctcgctgcagcggccctggcgggtgccctggctaccatggaccatcctgctggtcagcgctgcggctgcttcg
cagaatcaagaacggctatgtgcgtttaaagatccgtatcagcaagaccttgggataggtgagagtagaatctctcatgaaa
atgggacaatattatgctcgaaaggtagcacctgctatggcctttgggagaaatcaaaaggggacataaatcttgtaaaaca
aggatgttggtctcacattggagatccccaagagtgtcactatgaagaatgtgtagtaactaccactcctccctcaattcagaa
tggaacataccgtttctgctgttgtagcacagatttatgtaatgtcaactttactgagaattttccacctcctgacacaacaccac
tcagtccacctcattcatttaaccgagatgagacaataatcattgctttggcatcagtctctgtattagctgttttgatagttgcctt
atgctttggatacagaatgttgacaggagaccgtaaacaaggtcttcacagtatgaacatgatggaggcagcagcatccga
accctctcttgatctagataatctgaaactgttggagctgattggccgaggtcgatatggagcagtatataaaggctccttgga
tgagcgtccagttgctgtaaaagtgttttcctttgcaaaccgtcagaattttatcaacgaaaagaacatttacagagtgcctttg
atggaacatgacaacattgcccgctttatagttggagatgagagagtcactgcagatggacgcatggaatatttgcttgtgat
ggagtactatcccaatggatctttatgcaagtatttaagtctccacacaagtgact**g**ggtaagctcttgccgtcttgctcattctg
ttactagaggactggcttatcttcacacagaattaccacgaggagatcattataaacctgcaatttcccatcgagatttaaaca
gcagaaatgtcctagtgaaaaatgatggaacctgtgttattagtgactttggactgtccatgaggctgactggaaatagactg
gtgcgcccaggggaggaagataatgcagccataagcgaggttggcactatcagatatatggcaccagaagtgctagaag
gagctgtgaacttgagggactgtgaatcagctttgaaacaagtagacatgtatgctcttggactaatctattgggagatatttat
gagatgtacagacctcttcccaggggaatccgtaccagagtaccagatggcttttcagacagaggttggaaaccatcccac
ttttgaggatatgcaggttctcgtgtctagggaaaaacagagacccaagttcccagaagcctggaaagaaaatagcctggc
agtgaggtcactcaaggagacaatcgaagactgttgggaccaggatgcagaggctcggcttactgcacagtgtgctgagg
aaaggatggctgaacttatgatgatttgggaaagaaacaaatctgtgagcccaacagtcaatccaatgtctactgctatgcag
aatgaacgcaacctgtcacataataggcgtgtgccaaaaattggtccttatccagattattcttcctcctcatacattgaagact
ctatccatcatactgacagcatcgtgaagaatatttcctctgagcattctatgtccagcacacctttgactataggggaaaaaa
accgaaattcaattaactatgaacgacagcaagcacaagctcgaatccccagccctgaaacaagtgtcaccagcctctcca
ccaacacaacaaccacaaacaccacaggactcacgccaagtactggcatgactactatatctgagatgccatacccagat
gaaacaaatctgcataccacaaatgttgcacagtcaattgggccaacccctgtctgcttacagctgacagaagaagacttgg
aaaccaacaagctagacccaaaagaagttgataagaacctcaaggaaagctctgatgagaatctcatggagcactctctta
aacagttcagtggcccagacccactgagcagtactagttctagcttgctttacccactcataaaacttgcagtagaagcaact
ggacagcaggacttcacacagactgcaaatggccaagcatgtttgattcctgatgttctgcctactcagatctatcctctcccc

aagcagcagaaccttcccaagagacctactagtttgcctttgaacaccaaaaattcaacaaaagagccccggctaaaatttg
gcagcaagcacaaatcaaacttgaaacaagtcgaaactggagttgccaagatgaatacaatcaatgcagcagaacctcat
gtggtgacagtcaccatgaatggtgtggcaggtagaaaccacagtgttaactcccatgctgccacaacccaatatgccaat
gggacagtactatctggccaaacaaccaacatagtgacacatagggcccaagaaatgttgcagaatcagtttattggtgag
gacacccggctgaatattaattccagtcctgatgagcatgagcctttactgagacgagagcaacaagctggccatgatgaa
ggtgttctggatcgtcttgtggacaggagggaacggccactagaaggtggccgaactaattccaataacaacaacagcaat
ccatgttcagaacaagatgttcttgcacagggtgttccaagcacagcagcagatcctgggccatcaaagcccagaagagc
acagaggcctaattctctggatctttcagccacaaatgtcctggatggcagcagtatacagataggtgagtcaacacaagat
ggcaaatcaggatcaggtgaaaagatcaagaaacgtgtgaaaactccctattctcttaagcggtggcgcccctccacctgg
gtcatctccactgaatcgctggactgtgaagtcaacaataatggcagtaacagggcagttcattccaaatccagcactgctgt
ttaccttgcagaaggaggcactgctacaaccatggtgtctaaagatataggaatgaactgtctgtga-3’

**AHI1**

5’-

atgcctacagctgagagtgaagcaaaagtaaaaaccaaagttcgctttgaagaattgcttaagacccacagtgatctaatgc
gtgaaaagaaaaaactgaagaaaaaacttgtcaggtctgaagaaaacatctcacctgacactattagaagcaatcttcactat
atgaaagaaactacaagtgatgatcccgacactattagaagcaatcttccccatattaaagaaactacaagtgatgatgtaag
tgctgctaacactaacaacctgaagaagagcacgagagtcactaaaaacaaattgaggaacacacagttagcaactgaaa
atcctaatggtgatgctagtgtagaggaagacaaacaaggaaagccaaataaaaaggtgataaagacggtgccccagttg
actacacaagacctgaaaccggaaactcctgagaataaggttgattctacacaccagaaaacacatacaaagccacagcc
aggcgttgatcatcagaaaagtgagaaggcaaatgagggaagagaagagactgatttagaagaggatgaagaattgatg
caagcatatcagtgccatgtaactgaagaaatggcaaaggagattaagaggaaaataagaaagaaactgaaagaacagtt
gacttactttccctcagatactttattccatgatgacaaactaagcagtgaaaaaaggaaaaagaaaaaggaagttccagtctt
ctctaaagctgaaacaagtacattgaccatctctggtgacacagttgaaggtgaacaaaagaaagaatcttcagttagatca
gtttcttcagattctcatcaagatgatgaaataagctcaatggaacaaagcacagaagacagcatgcaagatgatacaaaac
ctaaaccaaaaaaaacaaaaaagaagactaaagcagttgcagataataatgaagatgttgatggtgatggtgttcatgaaat
aacaagccgagatagcccggtttatcccaaatgtttgcttgatgatgaccttgtcttgggagtttacattcaccgaactgatag
acttaagtcagattttatgatttctcacccaatggtaaaaattcatgtggttgatgagcatactggtcaatatgtcaagaaagatg
atagtggacggcctgtttcatcttactatgaaaaagagaatgtggattatattcttcctattatgacccagccatatgattttaaac
agttaaaatcaagacttccagagtgggaagaacaaattgtatttaatgaaaattttccctatttgcttcgaggctctgatgagag
tcctaaagtcatcctgttctttgagattcttgatttcttaagcgtggatgaaattaagaataattctgaggttcaaaaccaagaatg

tggctttcggaaaattgcctgggcatttcttaagcttctgggagccaatggaaatgcaaacatcaactcaaaacttcgcttgca
gctatattacccacctactaagcctcgatccccattaagtgttgttgaggcatttgaatggtggtcaaaatgtccaagaaatcat
tacccatcaacactgtacgtaactgtaagaggactgaaagttccagactgtataaagccatcttaccgctctatgatggctctt
caggaggaaaaaggtaaaccagtgcattgtgaacgtcaccatgagtcaagctcagtagacacagaacctggattagaaga
gtcaaaggaagtaataaagtggaaacgactccctgggcaggcttgccgtatcccaaacaaacacctcttctcactaaatgc
aggagaacgaggatgtttttgtcttgatttctcccacaatggaagaatattagcagcagcttgtgccagccgggatggatatc
caattattttatatgaaattccttctggacgtttcatgagagaattgtgtggccacctcaatatcatttatgatctttcctggtcaaa
agatgatcactacatccttacttcatcatctgatggcactgccaggatatggaaaaatgaaataaacaatacaaatactttcag
agttttacctcatccttcttttgtttacacggctaaattccatccagctgtaagagagctagtagttacaggatgctatgattccat
gatacggatat**g**gaaagttgagatgagagaagattctgccatattggtccgacagtttgacgttcacaaaagttttatcaactc
actttgttttgatactgaaggtcatcatatgtattcaggagattgtacaggggtgattgttgtttggaatacctatgtcaagattaat
gatttggaacattcagtgcaccactggactataaataaggaaattaaagaaactgagtttaagggaattccaataagttatttg
gagattcatcccaatggaaaacgtttgttaatccataccaaagacagtactttgagaattatggatctccggatattagtagca
aggaagtttgtaggagcagcaaattatcgggagaagattcatagtactttgactccatgtgggacttttctgtttgctggaagt
gaggatggtatagtgtatgtttggaacccagaaacaggagaacaagtagccatgtattctgacttgccattcaagtcacccat
tcgagacatttcttatcatccatttgaaaatatggttgcattctgtgcatttgggcaaaatgagccaattcttctgtatatttacgatt
tccatgttgcccagcaggaggctgaaatgttcaaacgctacaatggaacatttccattacctggaatacaccaaagtcaagat
gccctatgtacctgtccaaaactaccccatcaaggctcttttcagattgatgaatttgtccacactgaaagttcttcaacgaaga
tgcagctagtaaaacagaggcttgaaactgtcacagaggtgatacgttcctgtgctgcaaaagtcaacaaaaatctctcattt
acttcaccaccagcagtttcctcacaacagtctaagttaaagcagtcaaacatgctgaccgctcaagagattctacatcagttt
ggtttcactcagaccgggattatcagcatagaaagaaagccttgtaaccatcaggtagatacagcaccaacggtagtggct
ctttatgactacacagcgaatcgatcagatgaactaaccatccatcgcggagacattatccgagtgtttttcaaagataatgaa
gactggtggtatggcagcataggaaagggacaggaaggttattttccagctaatcatgtggctagtgaaacactgtatcaag
aactgcctcctgagataaaggagcgatcccctcctttaagccctgaggaaaaaactaaaatagaaaaatctccagctcctca
aaagcaatcaatcaataagaacaagtcccaggacttcagactaggctcagaatctatgacacattctgaaatgagaaaaga
acagagccatgaggaccaaggacacataatggatacacggatgaggaagaacaagcaagcaggcagaaaagtcactct
aatagagta-3’

**FANCC**

5’-

atggctcaagattcagtagatctttcttgtgattatcagttttggatgcagaagctttctgtatgggatcaggcttccactttggaa

acccagcaagacacctgtcttcacgtggctcagttccaggagttcctaaggaagatgtatgaagccttgaaagagatggatt
ctaatacagtcattgaaagattccccacaattggtcaactgttggcaaaagcttgttggaatccttttattttagcatatgatgaa
agccaaaaaattctaatatggtgcttatgttgtctaattaacaaagaaccacagaattctggacaatcaaaacttaactcctgga
tacagggtgtattatctcatatactttcagcactcagatttgataaagaagttgctcttttcactcaaggtcttgggtatgcaccta
tagattactatcctggtttgcttaaaaatatggttttatcattagcgtctgaactcagagagaatcatcttaatggatttaacactca
aaggcgaatggctcccgagcgagtggcgtccctgtcacgagtttgtgtcccacttattaccctgacagatgttgaccccctg
gtggaggctctcctcatctgtcatggacgtgaacctcaggaaatcctccagccagagttctttgaggctgtaaacgaggcca
ttttgctgaagaagatttctctccccatgtcagctgtagtctgcctctggcttcggcaccttcccagccttgaaaaagcaatgct
gcatctttttgaaaagctaatctccagtgagagaaattgtctgagaaggatcgaatgctttataaaagattcatcgctgcctcaa
gcagcctgccaccctgccatattccgggttgttgatgagatgttcaggtgtgcactcctggaaaccgatggggccctggaa
atcatagccactattcaggtgtttacgcagtgctttgtagaagctctggagaaagcaagcaagcagctgcggtttgcactcaa
gacctactttccttacacttctccatctcttgccatggtgctgctgcaagaccctcaagatatccctcggggacactggctcca
gacactgaagcatatttctgaactgctcagagaagcagttgaagaccagactcatgggtcctgcggaggtccctttgagag
ctggttcctgttcattcacttcggaggatgggctgagatggtggcagagcaattactgatgtcggcagccgaacccccac
ggccctgctgtggctcttggccttctactacggcccccgtgatgggaggcagcagagagcacagactatggtccaggtga
aggccgtgctgggccacctcctggcaatgtccagaagcagcagcctctcagcccaggacctgcagacggtagcaggac
agggcacagacacagacctcagagctcctgcacaacagctgatcaggcaccttctcctcaacttcctgctctgggctcctg
gaggccacacgatcgcctgggatgtcatcaccctgatggctcacactgctgagataactcacgagatcattggctttcttga
ccagaccttgtacagatggaatcgtcttggcattgaaagccctagatcagaaaaactggcccgagagctccttaaagagct
gcgaactcaagtctag-3’

**MYBPC3**

5’-

atgcctgagccggggaagaagccagtctcagcttttagcaagaagccacggtcagtggaagtggccgcaggcagccctg
ccgtgttcgaggccgagacagagcgggcaggagtgaaggtgcgctggcagcgcggaggcagtgacatcagcgccag
caacaagtacggcctggccacagagggcacacggcatacgctgacagtgcgggaagtgggccctgccgaccagggat
cttacgcagtcattgctggctcctccaaggtcaagttcgacctcaaggtcatagaggcagagaaggcagagcccatgctgg
cccctgcccctgcccctgctgaggccactggagcccctggagaagccccggccccagccgctgagctgggagaaagtg
ccccaagtcccaaagggtcaagctcagcagctctcaatggtcctacccctggagcccccgatgaccccattggcctcttcg
tgatgcggccacaggatggcgaggtgaccgtgggtggcagcatcaccttctcagcccgcgtggccggcgccagcctcct
gaagccgcctgtggtcaagtggttcaagggcaaatgggtggacctgagcagcaaggtgggccagcacctgcagctgca

cgacagctacgaccgcgccagcaaggtctatctgttcgagctgcacatcaccgatgcccagcctgccttcactggcagcta
ccgctgtgaggtgtccaccaaggacaaatttgactgctccaacttcaatctcactgtccacgaggccatgggcaccggaga
cctggacctcctatcagccttccgccgcacgagcctggctggaggtggtcggcggatcagtgatagccatgaggacactg
ggattctggacttcagctcactgctgaaaaagagagacagtttccggaccccgagggactcgaagctggaggcaccagc
agaggaggacgtgtgggagatcctacggcaggcacccccatctgagtacgagcgcatcgccttccagtacggcgtcact
gacctgcgcggcatgctaaagaggctcaagggcatgaggcgcgatgagaagaagagcacagcctttcagaagaagctg
gagccggcctaccaggtgagcaaaggccacaagatccggctgaccgtggaactggctgaccatgacgctgaggtcaaa
tggctcaagaatggccaggagatccagatgagcggcagcaagtacatctttgagtccatcggtgccaagcgtaccctgac
catcagccagtgctcattggcggacgacgcagcctaccagtgcgtggtgggtggcgagaagtgtagcacggagctctttg
tgaaagagccccctgtgctcatcacgcgccccttggaggaccagctggtgatggtggggcagcgggtggagtttgagtgt
gaagtatcggaggagggggcgcaagtcaaatggctgaaggacggggtggagctgacccgggaggagaccttcaaata
ccggttcaagaaggacgggcagagacaccacctgatcatcaacgaggccatgctggaggacgcggggcactatgcact
gtgcactagcgggggccaggcgctggctgagctcattgtgcaggaaaagaagctggaggtgtaccagagcatcgcaga
cctgatggtgggcgcaaaggaccaggcggtgttcaaatgtgaggtctcagatgagaatgttcggggtgtgtggctgaaga
atgggaaggagctggtgcccgacagccgcataaaggtgtcccacatcgggcgggtccacaaactgaccattgacgacgt
cacacctgccgacgaggctgactacagctttgtgcccgagggcttcgcctgcaacctgtcagccaagctccacttcatgga
ggtcaagattgacttcgtacccaggcaggaacctcccaagatccacctggactgcccaggccgcataccagacaccattg
tggttgtagctggaaataagctacgtctggacgtccctatctctggggaccctgctcccactgtgatctggcagaaggctatc
acgcaggggaataaggccccagccaggccagccccagatgccccagaggacacaggtgacagcgatgagtgggtgtt
tgacaagaagctgctgtgtgagaccgagggccgggtccgcgtggagaccaccaaggaccgcagcatcttcacggtcga
gggggcagagaaggaagatgagggcgtctacacggtcacagtgaagaaccctgtgggcgaggaccaggtcaacctca
cagtcaaggtcatcgacgtgccagacgcacctgcggcccccaagatcagcaacgtgggagaggactcctgcacagtac
agtgggagccgcctgcctacgatggcgggcagcccatcctgggctacatcctggagcgcaagaagaagaagagctacc
ggtggatgcggctgaacttcgacctgattcaggagctgagtcatgaagcgcggcgcatgatcgagggcgtggtgtacga
gatgcgcgtctacgcggtcaacgccatcggcatgtccaggcccagccctgcctcccagcccttcatgcctatcggtccccc
cagcgaacccacccacctggcagtagaggacgtctctgacaccacggtctccctcaagtggcggcccccagagcgcgt
gggagcaggaggcctggatggctacagcgtggagtactgcccagagggctgctcagagtgggtggctgccctgcaggg
gctgacagagcacacatcgatactggtgaaggacctgcccacgggggcccggctgcttttccgagtgcgggcacacaat
atggcagggcctggagcccctgttaccaccacggagccggtgacagtgcaggagatcctgcaacggccacggcttcag
ctgcccaggcacctgcgccagaccattcagaagaaggtcggggagcctgtgaaccttctcatccctttccagggcaagcc
ccggcctcaggtgacctggaccaaagaggggcagcccctggcaggcgaggaggtgagcatccgcaacagccccaca

gacaccatcctgttcatccgggccgctcgccgcgtgcattcaggcacttaccaggtgacggtgcgcattgagaacatgga

ggacaaggccacgctggtgctgcaggttgttgacaagccaagtcctccccaggatctccgggtgactgacgcctggggtc

ttaatgtggctctggagtggaagccaccccaggatgtcggcaacacggagctct**g**gggtacacagtgcagaaagccga

caagaagaccatggagtggttcaccgtcttggagcattaccgcgcacccactgcgtggtgccagagctcatcattggcaa

tggctactacttccgcgtcttcagccagaatatggttggctttagtgacagagcggccaccaccaaggagcccgtctttatcc

ccagaccaggcatcacctatgagccacccaactataaggccctggacttctccgaggccccaagcttcacccagcccctg

gtgaaccgctcggtcatcgcgggctacactgctatgctctgctgtgctgtccggggtagccccaagcccaagatttcctggt

tcaagaatggcctggacctgggagaagacgcccgcttccgcatgttcagcaagcagggagtgttgactctggagattaga

aagccctgcccctttgacgggggcatctatgtctgcagggccaccaacttacagggcgaggcacggtgtgagtgccgcct

ggaggtgcgagtgcctcagtga-3’

**IL2RG**

5’-

atgttgaagccatcattaccattcacatccctcttattcctgcagctgcccctgctgggagtggggctgaacacgacaattctg

acgcccaatgggaatgaagacaccacagctgatttcttcctgaccactatgcccactgactccctcagtgtttccactctgcc

cctcccagaggttcagtgttttgtgttcaatgtcgagtacatgaattgcacttggaacagcagctctgagccccagcctacca

acctcactctgcattattggtacaagaactcggataatgataaagtccagaagtgcagccactatctattctctgaagaaatca

cttctggctgtcagttgcaaaaaaaggagatccacctctaccaaacatttgttgttcagctccaggacccacgggaacccag

gagacaggccacacagatgctaaaactgcagaatctggtgatcccctgggctccagagaacctaacacttcacaaactga

gtgaatcccagctagaactgaactggaacaacagattcttgaaccactgtttggagcacttggtgcagtaccggactgactg

ggaccacagctggactgaacaatcagtggattatagacataagttctccttgcctagtgtggatgggcagaaacgctacac

gtttcgtgttcggagccgctttaacccactctgtggaagtgctcagcatt**g**gagtgaatggagccacccaatccactggggg

agcaatacttcaaaagagaatcctttcctgtttgcattggaagccgtggttatctctgttggctccatgggattgattatcagcct

tctctgtgtgtatttctggctggaacggacgatgccccgaattcccaccctgaagaacctagaggatcttgttactgaatacca

cgggaacttttcggcctggagtggtgtgtctaagggactggctgagagtctgcagccagactacagtgaacgactctgcct

cgtcagtgagattcccccaaaaggaggggcccttggggaggggcctggggcctccccatgcaaccagcatagcccctа

ctgggcccccccatgttacaccctaaagcctgaaacctga-3’
