## Supplementary Table 1 for "Leveraging Endogenous ADAR for Programmable Editing on RNA"

**Supplementary Table 1 |** LbuCas13 crRNA sequences.

| **Name** | **Sequence** | **Source** |
| --- | --- | --- |
| LbuCas13/Cas13a crRNA scaffold | ggaccaccccaaaaaugaaggggacuaaaac | Extended Data Figure 1 |
| Ctrl crRNA_70_ | aaaccgagggaucauaggggacugaauccaccauucuucucccaaucccugcaacuccuucuuccccugc | Extended Data Figure 1 |
| Spacer of crRNA_15_ | gcagagccucCagc | Extended Data Figure 1 |
| Spacer of crRNA_22_ | cucacuggcagagccucCagc | Extended Data Figure 1 |
| Spacer of crRNA_28_ | cccuugcucacuggcagagccucCagc | Extended Data Figure 1 |
| Spacer of crRNA_35_ | cucucgcccuugcucacuggcagagccucCagc | Extended Data Figure 1 |
| Spacer of crRNA_40_ | cucucgcccuugcucacuggcagagccucCagcaucgc | Extended Data Figure 1 |
| Spacer of crRNA_47_ | ugaacagcucucgcccuugcucacuggcagagccucCagcaucgc | Extended Data Figure 1 |
| Spacer of crRNA_70_ | ugaacagcuccucgcccuugcucacuggcagagcccucCagcaucgcgagcaggcgcugccuccuccgcc | Extended Data Figure 1 |
