## Supplementary Table 3 for "Leveraging Endogenous ADAR for Programmable Editing on RNA"

**Supplementary Table** **3 |** Disease-related cDNAs used in this study.

| **Candidate** | **Disease** | **Mutant Adenosine** |
| --- | --- | --- |
| NM_000090.3 (*COL3A1*) | Ehlers-Danlos syndrome, type 4 | c.3833G>A (p.Trp1278Ter) |
| NM_001204.6 *(BMPR2*) | Primary pulmonary hypertension | c.893G>A (p.Trp298Ter) |
| NM_017651.4 (*AHI1*) | Joubert syndrome 3 | c.2174G>A (p.Trp725Ter) |
| NM_000136.2 (*FANCC*) | Fanconi anemia, complementation group C | c.1517G>A (p.Trp506Ter) |
| NM_000256.3 (*MYBPC3*) | Primary familial hypertrophic cardiomyopathy | c.3293G>A (p.Trp1098Ter) |
| NM_000206.2 (*IL2RG*) | X-linked severe combined immunodeficiency | c.710G>A  (p.Trp237Ter) |
